# Parental-effect gene-drive elements under partial selfing, or why do *Caenorhabditis* genomes have hyperdivergent regions?

**DOI:** 10.1101/2024.07.23.604817

**Authors:** Matthew V. Rockman

## Abstract

Self-fertile *Caenorhabditis* nematodes carry a surprising number of *Medea* elements, alleles that act in heterozygous mothers and cause death or developmental delay in offspring that don’t inherit them. At some loci, both alleles in a cross operate as independent *Medeas*, affecting all the homozygous progeny of a selfing heterozygote. The genomic coincidence of *Medea* elements and ancient, deeply coalescing haplotypes, which pepper the otherwise homogeneous genomes of these animals, raises questions about how these apparent gene-drive elements persist for long periods of time. Here I investigate how mating system affects the evolution of *Medeas*, and their paternal-effect counterparts, *peels*. Despite an intuition that antagonistic alleles should induce balancing selection by killing homozygotes, models show that, under partial selfing, antagonistic elements experience positive frequency dependence: the common allele drives the rare one extinct, even if the rare one is more penetrant. Analytical results for the threshold frequency required for one allele to invade a population show that a very weakly penetrant allele, one whose effects would escape laboratory detection, could nevertheless prevent a much more penetrant allele from invading under high rates of selfing. Ubiquitous weak antagonistic *Medeas* and *peels* could then act as localized barriers to gene flow between populations, generating genomic islands of deep coalescence. Analysis of gene expression data, however, suggest that this cannot be the whole story. A complementary explanation is that ordinary ecological balancing selection generates ancient haplotypes on which *Medeas* can evolve, while high homozygosity in these selfers minimizes the role of gene drive in their evolution.

## INTRODUCTION

Populations harbor genetic variation, and its distribution is a function of mutation, recombination, natural selection, and many aspects of population biology. Models that explain heterogeneity in variation along a genome invoke various combinations of these factors. Recently, researchers have found that the genomes of some *Caenorhabditis* nematodes are mosaics of some regions that are hyperdivergent and consistent with very ancient coalescence amid other regions that are depauperate in variation and likely share very recent ancestry across the species (Thompson *et al*. 2015; Lee *et al*. 2021; Noble *et al*. 2021; Stevens *et al*. 2022). Efforts to understand these patterns have focused on balancing selection as a mechanism that preserves diversity in hyperdivergent regions while allowing homogenization of the rest of the genome by drift and selective sweeps.

Experimental crosses among wild isolates revealed a surprising candidate mechanism for balancing selection. Studies showed that each of the three partially-selfing species of *Caenorhabditis* carry polymorphic postzygotic gene-drive elements. *Medea* elements are alleles that cause reduced fitness in offspring that lack them but whose mother carried such an allele: Maternal-Effect Dominant Embryonic Arrest (Beeman *et al*. 1992). Analogous *peel* elements depend on the father’s genotype: Paternal-effect Epistatic Embryonic Lethal (Seidel *et al*. 2008). *Medea* and *peel* are typically conceptualized as toxin-antidote systems (Figure 1A), and in a few cases this sort of mechanism has been experimentally confirmed at the molecular level (Seidel *et al*. 2011; BEN-DAVID *et al*. 2017; Caro *et al*. 2024). They have also been synthetically generated and validated as gene-drive systems (Chen *et al*. 2007). Each of the mapped *Medea* and *peel* elements in *Caenorhabditis* falls within a region of genome characterized by hyperdivergent, ancient haplotypes.

**Figure 1.**
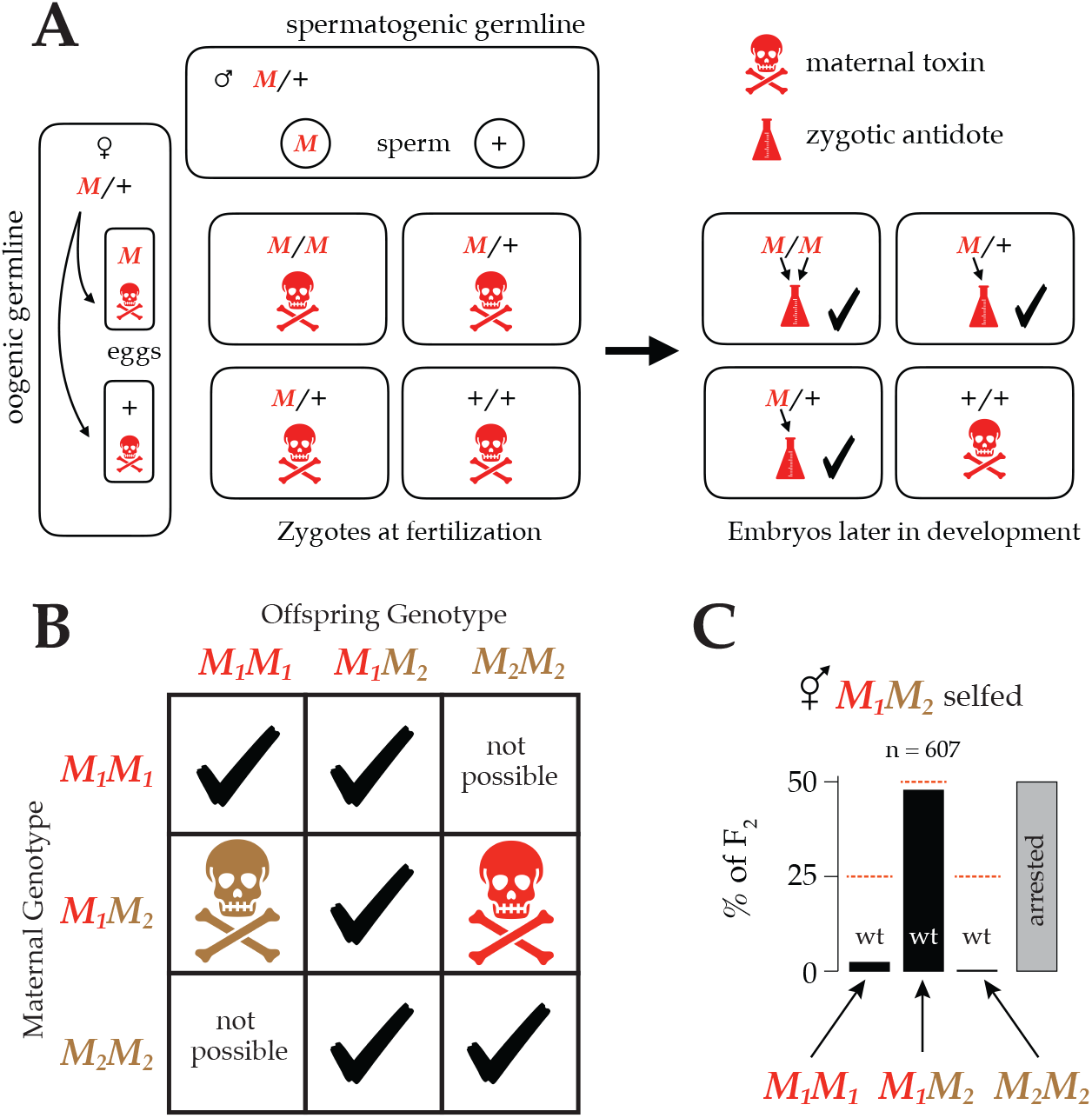
**A**. A cross between two heterozygotes (or a self-fertilization by one) results in a Punnett Square with Mendelian segregation, *M/M, M*/+, and +/+ zygotes in a 1:2:1 ratio. A *Medea* allele (*M*) deposits a toxin (skull and crossbones) in every egg. Later, when zygotic transcription starts, embryos carrying an *M* allele synthesize an antidote (beaker). The quarter of embryos that lack an *M* allele suffer the effects of the maternal toxin. **B**. In a population segregating *Medea* alleles, the phenotype of an embryo depends on its own genotype and that of its mother. There are seven possible combinations of maternal and zygotic genotype at one locus. If both alleles at the locus have *Medea* activity, one combination suffers the effects of the *M*_*1*_ allele (red) and another suffers the effects of the *M*_*2*_ allele (brown). **C**. An example of antagonistic *Medea* elements killing the majority of both classes of F_2_ homozygous offspring of a heterozygous hermaphrodite. These data are from self-fertilization of *C. tropicalis* hermaphrodites heterozygous at a locus on chromosome V with antagonistic *Medeas, M*_*1*_ and *M*_*2*_, adapted from Figure 6 of Noble *et al*. (2021). Nearly all the wild-type F_2_s are heterozygotes; the homozygotes mostly exhibit arrested development.

Remarkably, several studies found that *C. tropicalis* harbors so many such elements that they are sometimes allelic to one another: the two alleles at a single locus each behave as a *Medea* (BEN-DAVID *et al*. 2021; Noble *et al*. 2021; Pliota *et al*. 2024). In such cases, a heterozygous mother deposits two independent toxins into each egg, and only zygotes that inherit a copy of each allele escape the effects of the toxins (Figure 1B-C). Because only these heterozygous offspring retain wild-type fitness, antagonistic *Medea* elements plausibly induce balancing selection at these loci via overdominance: heterozygote advantage.

Despite appearances, forward simulations showed that antagonistic *Medea* elements actually do not cause overdominant balancing selection, and instead they induce positive frequency dependence: the allele that is initially more common in the population drives the rarer allele to extinction (Noble *et al*. 2021). A key requirement for this frequency dependence is a mixed-mating system, with a combination of self-fertilizations and outcrosses. Here I develop mathematical models to better understand the fate of *Medea* and *peel* alleles in partially-selfing populations, with particular attention to the question of natural variation in *Caenorhabditis* populations. I also review the assumptions about molecular and population biology that go into these models.

I find that selfing has dramatic effects on the evolution of these alleles, and that antagonistic alleles experience strong positive frequency dependence. These analytical results raise the possibility that hyperdivergent haplotypes evolve and persist not via overdominance but via an underdominance-like mechanism, where alleles fixed in subpopulations can exclude alleles from other subpopulations, while the rest of the genome experiences gene flow. The positive-frequency dependence is so strong that very weakly penetrant *Medea* and *peel* alleles could generate the required selective force, and experimental studies would have overlooked them. The overall picture then is that ubiquitous weak parental-by-zygotic genetic interactions could contribute to the mosaic pattern of genetic variation in partially selfing *Caenorhabditis*, and perhaps in other species as well.

## METHODS

I modeled the evolution of *Medea* and *peel* genotype frequencies in a deterministic framework with discrete generations, as detailed below in the results. Figures illustrating allele frequency dynamics were produced by iterating recursion equations in R (v.4.0.2) (R_CORE_TEAM 2020). The ternary plots make use of the package *Ternary* (v1.2.3) (SMITH 2017). To approximate unstable equilibria numerically for antagonistic *Medea/peel* elements under androdioecy (Figure 7 and Supplementary Figures 1 and 2), I used the recursion equations to ask, for each starting frequency from 0.001 to 0.999, with steps of size 0.001, whether the allele frequency was higher or lower than the starting frequency after 100 generations. I performed this analysis for selfing rates from 0 to 0.99 with steps of size 0.01. Code to replicate and extend the analyses is provided in the associated R package, *MedeaFight*.

**Figure 2.**
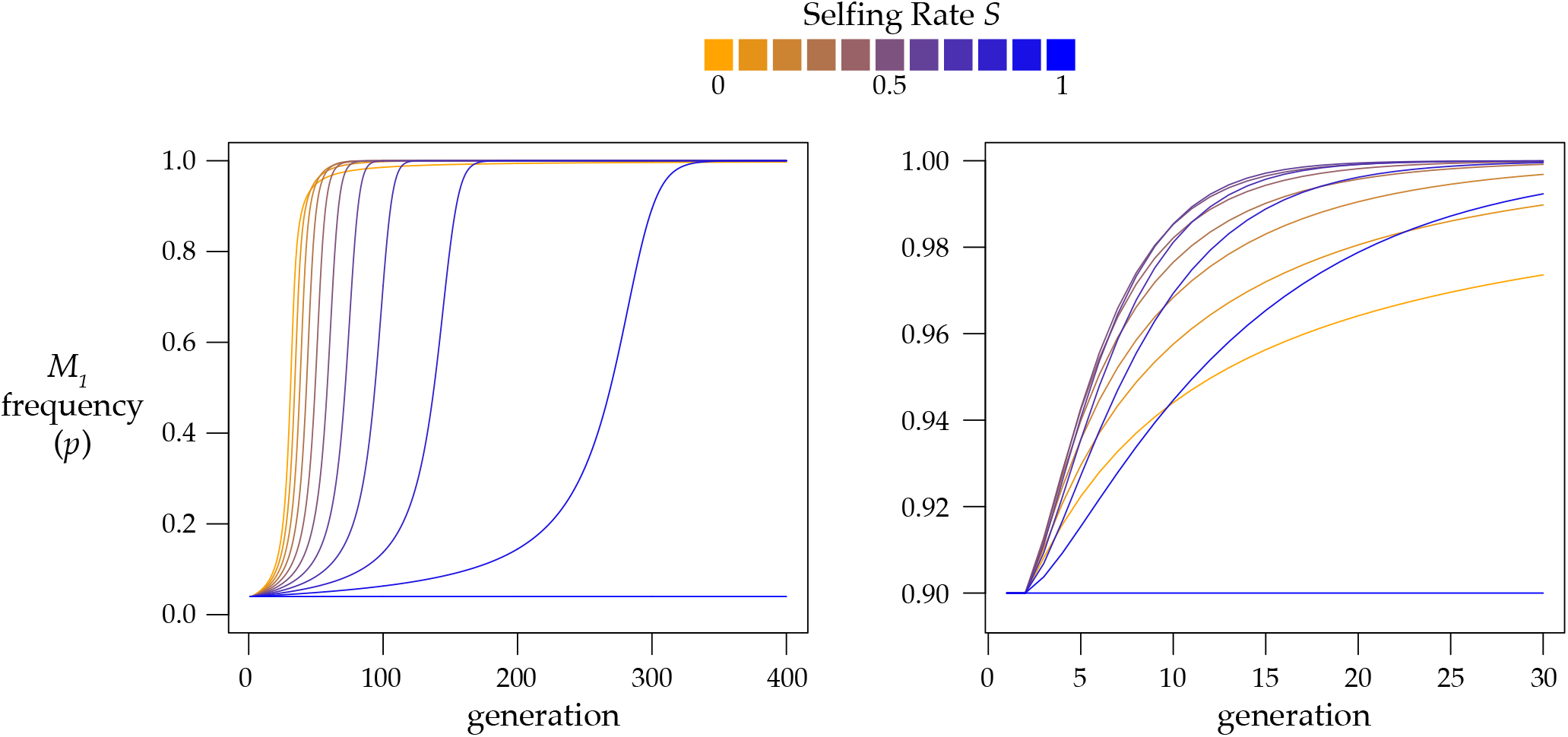
With outcrossing or partial selfing, a *Medea* element sweeps to fixation. These plots show results found by iterating equation 2 for eleven values of selfing rate *S* from 0 to 1, with penetrance *k*_*1*_ = 1. At left, initial allele frequency *p* = 0.04, and selfing slows the spread of the *Medea*. At right, initial allele frequency *p* = 0.90, and partial selfing hastens the fixation of the *Medea*.

Gene expression analysis used functions *merge* and *intersect* from bedtools v2.30.0 (QUINLAN AND HALL 2010) to process positions of hyperdivergent regions from CeNDR release 20220216 (Cook *et al*. 2017), and positions from WormBase ws283 (Davis *et al*. 2022) of genes from table S11 from Tzur et al. (2018), table S1 from Billi et al. (2013), table S1 from Bezler et al (2019).

## RESULTS

Consider a monoecious population, consisting of hermaphrodites capable of both selfing and outcrossing. This mating system is very different from that of androdioecious *Caenorhabditis* nematodes, but it provides a simple starting point, and in later sections of the paper I extend the analysis to the *Caenorhabditis* mating system.

In the monoecious model, hermaphrodites produce progeny by selfing with fixed frequency *S* and by randomly mating with frequency 1-*S*. At a single biallelic locus, each allele has the potential to be a *Medea* element. A *Medea* acts in the oogenic germline to deposit a toxin into every egg, and embryos that do not inherit the *Medea* allele die with frequency *k*, the killing penetrance of the element. If both alleles at the locus have *Medea* activity, allele *M*_*1*_ kills *M*_*2*_ homozygotes with penetrance *k*_*1*_ and *M*_*2*_ kills *M*_*1*_ homozygotes with penetrance *k*_*2*_. The three genotypes, *M*_*1*_*M*_*1*_, *M*_*1*_*M*_*2*_, and *M*_*2*_*M*_*2*_, have frequencies *X, Y*, and *Z*, which sum to 1. From this model, we can write down the frequencies of the various self-fertilizations and outcrosses and the expected genotypes of the surviving progeny of each cross after Mendelian segregation and *Medea*-mediated killing.

| Egg-parent | Sperm-parent | Frequency | Surviving Progeny Genotypes |  |  |
| --- | --- | --- | --- | --- | --- |
| | | | $M_1M_1$ | $M_1M_2$ | $M_2M_2$ |
| | $M_1M_1$ | $SX$ | 1 | | |
| | $M_1M_2$ | $SY$ | $(1-k_2)/4$ | $1/2$ | $(1-k_1)/4$ |
| | $M_2M_2$ | $SZ$ | | | 1 |
| $M_1M_1$ | $M_1M_1$ | $(1-S)X^2$ | 1 | | |
| $M_1M_1$ | $M_1M_2$ | $(1-S)XY$ | $1/2$ | $1/2$ | |
| $M_1M_1$ | $M_2M_2$ | $(1-S)XZ$ | | 1 | |
| $M_1M_2$ | $M_1M_1$ | $(1-S)XY$ | $(1-k_2)/2$ | $1/2$ | |
| $M_1M_2$ | $M_1M_2$ | $(1-S)Y^2$ | $(1-k_2)/4$ | $1/2$ | $(1-k_1)/4$ |
| $M_1M_2$ | $M_2M_2$ | $(1-S)YZ$ | | $1/2$ | $(1-k_1)/2$ |
| $M_2M_2$ | $M_1M_1$ | $(1-S)XZ$ | | 1 | |
| $M_2M_2$ | $M_1M_2$ | $(1-S)YZ$ | | $1/2$ | $1/2$ |
| $M_2M_2$ | $M_2M_2$ | $(1-S)Z^2$ | | | 1 |

This table shows, for example, that zygotes arising from a cross between an *M*_*1*_*M*_*2*_ egg-parent and *M*_*2*_*M*_*2*_ sperm-parent will be (1-*S*)*YZ* of all zygotes. By Mendelian segregation, 1/2 of such zygotes will be *M*_*2*_*M*_*2*_ homozygotes. Fraction *k*_1_ of these will be killed by the *M*_*1*_ *Medea* in their mother’s genome, and fraction 1-*k*_1_ of them will survive.

With a bit of tedious book keeping we can turn this table into recursion equations for the frequencies of the three genotypes in succeeding generations. The *M*_*1*_*M*_*1*_ homozygotes in the next generation arise from six different parental types, as shown in the *M*_*1*_*M*_*1*_ column of the table. The frequency of *M*_*1*_*M*_*1*_ homozygotes in the next generation (written *X’*) is the sum of the progeny from these six types normalized by the sum of the whole table, which is the mean fitness of the population, abbreviated 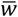.

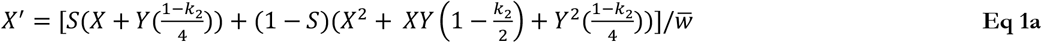

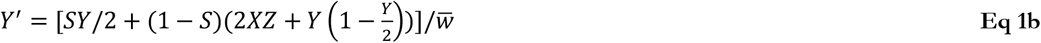

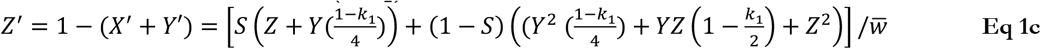

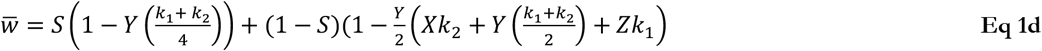

For a paternal-effect *peel* element, the table is slightly different – only the heterozygotes acting as males in crosses have *peel*-affected offspring – but the resulting recursions are the same, even in the case of antagonism between one haplotype that bears a *Medea* and one that bears a *peel*.

Consequently, for a monoecious species with self- and cross-compatible hermaphrodites, *Medea* and *peel* elements behave identically in this model.

The frequency of *M*_*1*_ allele is *X* + *Y*/2 = *p*. This relation lets us recast the recursions more succinctly in terms of *M*_*1*_ allele frequency *p* and heterozygosity *Y*:

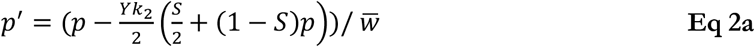

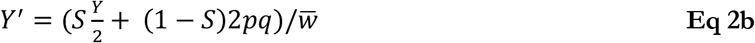

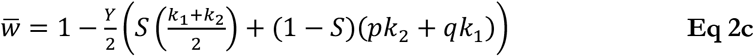

where *q* = 1-*p* is the frequency of the *M*_2_ allele.

Here the recursions have been written to highlight the relative contributions of selfing and outcrossing. The effects of the former depend only on heterozygosity, while those of the latter depends also on the allele frequencies. The system as a whole is at equilibrium when *p* = 0 or when *p* = 1 (in both cases heterozygosity *Y* is necessarily 0). A special case of the parameters is complete selfing in the total absence of heterozygotes (*S*=1, *Y*=0). In this case every value of *p* is an equilibrium; allele frequencies remain constant because selection requires heterozygotes and without outcrossing none are generated. However, if 0<*p*<1, any amount of outcrossing will restore heterozygotes in the next generation (that is, when *Y*=0,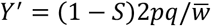).

Next, I will find internal equilibria for allele frequency *p* and heterozygosity *Y*. The results are not conducive to easy intuitions, but I will detail several specific scenarios to show how these equations provide insight into the behavior of *Medea* elements in populations with partial selfing.

If we rearrange Eqq 2 to emphasize the relative contributions of the two *Medea* alleles, the equations take the following forms:

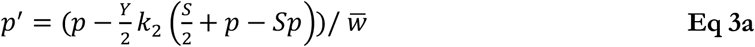

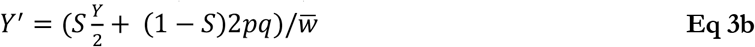

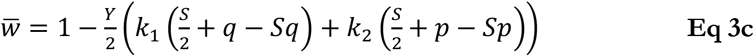

These versions of the recursions make it easier to find internal allele-frequency equilibria 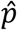 by setting *p = p’* and rearranging to find this implicit solution:

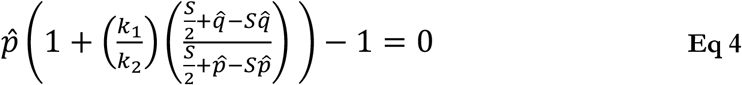

This equation reveals that internal equilibria for allele frequency *p* are independent of heterozygosity *Y*, and further that the *Medea* penetrances *k*_*1*_ and *k*_*2*_ affect the equilibria only through their ratio. An explicit equilibrium for *p* can be found by rewriting equation 4 as a polynomial and then determining its roots in the range 0 to 1:

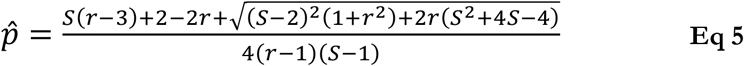

where *r* = *k*_*1*_*/k*_*2*_, the ratio of the *Medea* killing penetrances. At 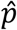, the equilibrium heterozygosity is given by

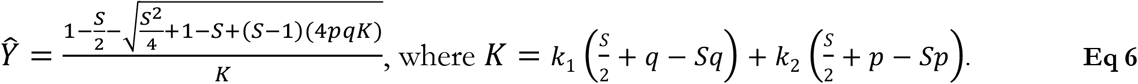

### Case 1: a single *Medea*

Wade and Beeman (WADE AND BEEMAN 1994) showed that a *Medea* element will invade and sweep to fixation in a randomly mating population. That result holds under partial selfing.

With *k*_*2*_ = 0, so that only allele *M*_*1*_ has *Medea* activity, the recursion equation for *M*_*1*_ frequency *p* (eqq. 3) becomes 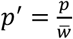, with 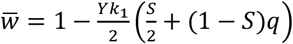. Because 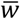. is always less than or equal to 1, the change in *M*_*1*_ allele frequency (*p’-p* = Δ*p*) is always positive (when *Y* >0) or 0 (when *Y* = 0). Whenever *S*<1 and 0<*p*<1, outcrossing will generate heterozygotes, keeping Δ*p* positive and driving *M*_*1*_ to fixation.

When rare, *M*_*1*_ alleles increase in frequency most quickly under high rates of outcrossing (Figure 2). This is due both to the higher rate of production of heterozygotes and to the high probability that outcrossing heterozygotes will mate with the prevalent *M*_*1*_-susceptible *M*_*2*_ homozygotes, inducing the selective deaths of *k*_*1*_/2 of the cross progeny. When the *M*_*1*_ allele becomes very common, however, outcrossing is less efficient, as heterozygotes will tend to mate with the now-common *M*_*1*_homozygotes, resulting in 100% viability of the cross. If a heterozygote instead reproduces by self-fertilization, it will reliably generate and kill off *k*_*1*_/4 of its progeny. The result is that partial selfing actually hastens the fixation of the *Medea* allele (Figure 2). This result was previously shown by stochastic simulation (Noble *et al*. 2021).

### Case 2: Equal-penetrance antagonistic *Medea* alleles

When the ratio of penetrances *r* = 1 (i.e., *k*_*1*_ *= k*_*2*_ = *k)*, 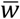. is simply 1-*Yk*/2 and the recursion for *M*_*1*_ frequency *p* works out to 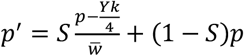. The second term shows that under random mating (*S*=0), allele frequencies do not change; *p’* = *p* for all *p*. This bizarre result, that alleles are neutral despite very strong selection at the genotypic level and dramatically reduced population mean fitness, was initially discovered by Hedrick (HEDRICK 1997), who was analyzing an instance of maternal-fetal interaction analogous to *Medea* elements (WADE 2000). That this phenomenon yields genetic drift in finite populations was shown by stochastic simulation (Noble *et al*. 2021).

The difference equation is 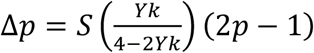. Recalling that *p* = 0 and *p* = 1 are equilibria, we find an additional equilibrium at 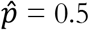. This equilibrium is unstable under partial selfing. When 0>*S*>1 and 0<*p*<0.5, Δ*p* is negative and the rarer allele is eliminated as *p* goes to 0. When *M*_*1*_ is the common allele, 0.5>*p*>1, Δ*p* is positive and again the rarer allele is eliminated as *p* goes to 1. This pattern of positive frequency dependence was previously discovered by Wade (WADE 2000), who used a model with a fixed inbreeding parameter *F* to extend Hedrick’s (HEDRICK 1997) results.

The dynamics are complicated (Figure 3A), as they depend on interactions among *S, k, Y*, and *p*. In general, the rarer allele is eliminated most rapidly at intermediate selfing rates, while both high and low rates of selfing result in a slower elimination of the rarer allele.

**Figure 3.**
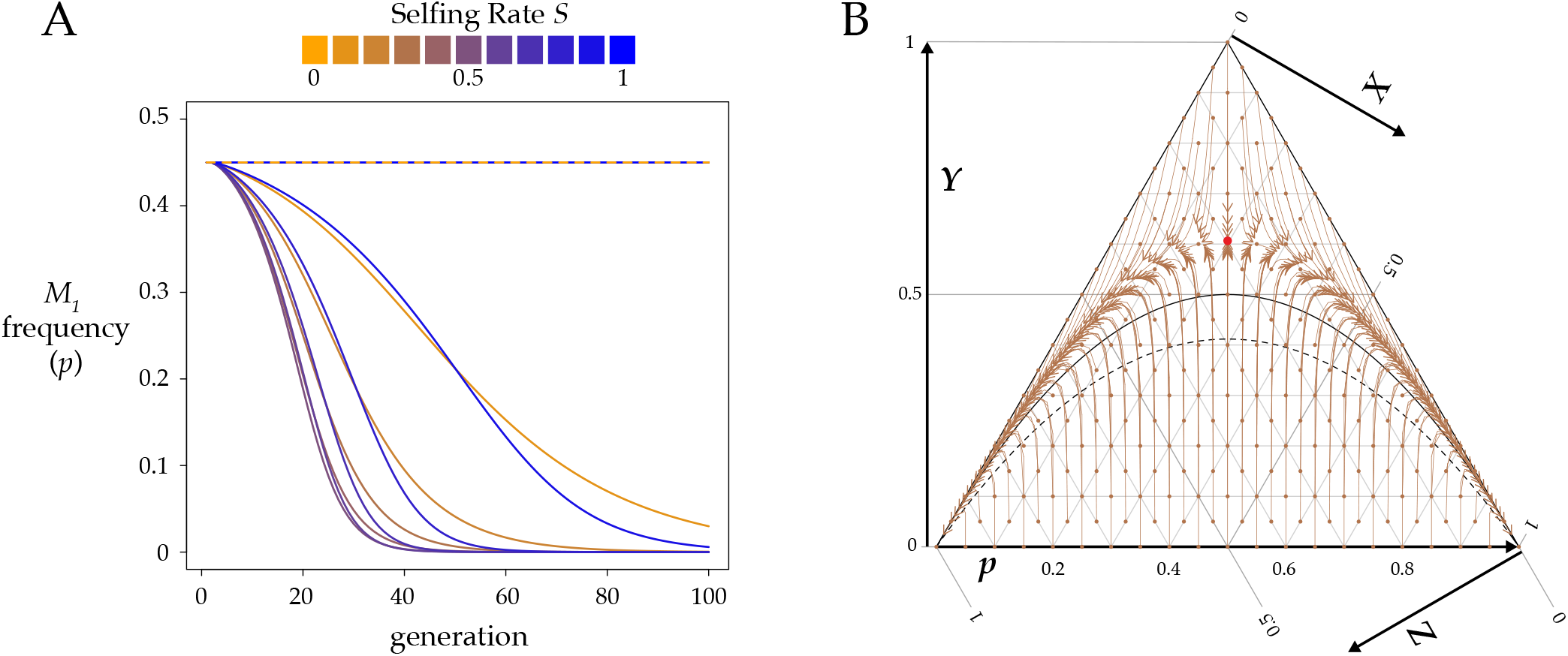
Partial selfing causes the elimination of the rarer of two equally penetrant *Medeas*. **A**. This plot shows allele frequencies found by iterating equation 2 for eleven values of *S* from 0 to 1, with *k* = 0.9 and starting from allele frequency *p* = 0.45 and heterozygosity *Y* =0. The dashed orange and blue line represents results for both *S*=0 and *S*=1. **B**. This ternary (De Finetti) plot shows evolution of this system (*k* = 0.9, *S* = 0.3) in genotype frequency space (equation 1). Each segment starts from a small dot, a combination of the three genotype frequencies (points spaced every 0.05). Each segment then plots genotype frequencies through six generations, with the arrows pointing to the position of the population in generation 6. The solid black curve shows genotype frequencies at Hardy-Weinberg equilibrium, and the dashed curve shows genotype frequencies under the neutral extended Hardy Weinberg equilibrium with selfing. The red dot shows the unstable internal equilibrium at 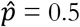 and *Ŷ* given by equation 7 (approximately 0.61 for these parameter values). The stable equilibria are in the lower corners, at *Z*=1 (*M*_*2*_ fixed) and *X*=1 (*M*_*1*_ fixed).

Equilibrium heterozygosity under obligate outcrossing was found by Hedrick (HEDRICK 1997). The general version with selfing is given by

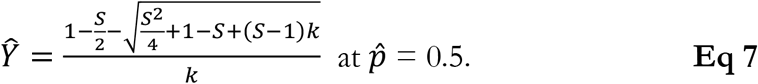

As shown in figure 3B, heterozygosity is elevated considerably above the value expected under neutrality 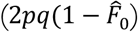, where the equilibrium neutral fixation index 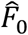 is *S*/(2-*S*)). The elevation is greater for high *k* and low *S*. The fixation index (*F* = 1 − *Ŷ*/2*pq*) is actually negative for a large part of the parameter space (at *p*=0.5, *F*<0 when *S* < *k*/2). In other words, there is an absolute excess of heterozygotes, even in the face of selfing.

### Case 3: Antagonistic *Medeas* with unequal penetrance

The discovery of multiple segregating *Medea* alleles within *Caenorhabditis tropicalis* raises the question of whether a *Medea* can invade a population that already carries a *Medea* at the same locus. The dynamics of the system depend on interactions among *S, k*_*1*_, *k*_*2*_, *Y*, and *p*. The equilibria are somewhat simpler, as there are only three for *p*: 0, 1, and as shown in equation 5, an internal equilibrium that depends on selfing rate *S* and *Medea* penetrance ratio *r* = *k*_*1*_*/k*_*2*_. As seen in figure 4, the internal equilibrium is unstable, and the system evolves to *p*=0 or *p*=1. A representative vector field is plotted in Figure 4B.

**Figure 4.**
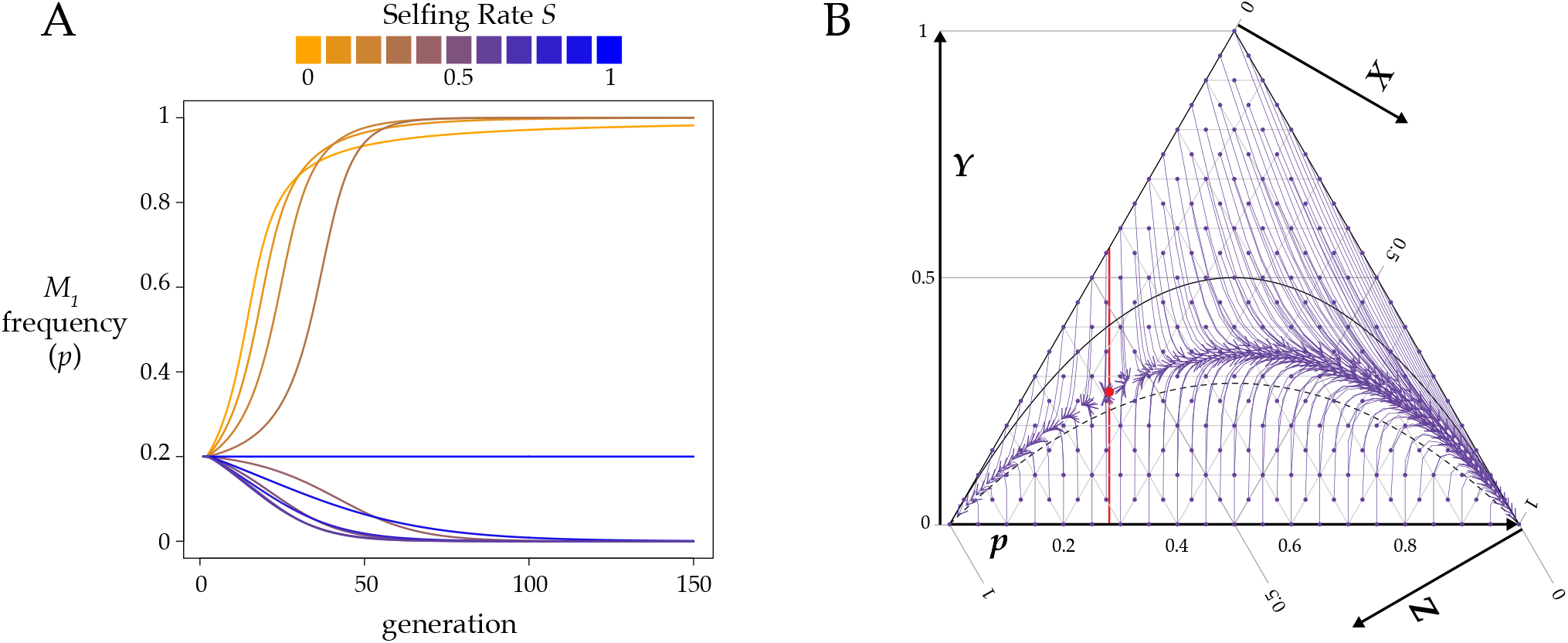
Partial selfing prevents a strong *Medea* from invading a population with a weaker resident *Medea*. **A**. This plot shows allele frequencies found by iterating equation 2 for eleven values of *S* from 0 to 1, with initial *M*_*1*_ frequency *p* = 0.2 and heterozygosity *Y* =0. The resident *Medea M*_*2*_ has penetrance *k*_*2*_ = 0.5 and the invading *Medea M*_*1*_ has penetrance *k*_*1*_ = 0.9. Under high outcrossing *M*_*1*_ sweeps to fixation but under high selfing it is eliminated from the population. Equation 8 gives the critical value of *S* for these conditions as approximately 0.37. **B**. A ternary plot for the same *k*_*1*_ and *k*_*2*_ as in panel A, here with *S* = 0.6. The red line marks the internal 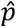. To the right of this line, *M*_*1*_ fixes, and to the left, *M*_*1*_ is eliminated. The red dot marks the unstable equilibrium given by equations 5 and 6 (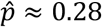, Ŷ ≈ 0.27).

The internal 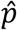 represents the critical allele frequency threshold above which *M*_*1*_ will spread and fix and below which *M*_*1*_ will be eliminated from the population. A *Medea* can never invade from low frequency when its penetrance is lower than that of the resident allele (i.e, *r* < 1, *p* < 0.5). However, even a stronger *Medea* can only invade and displace a weaker resident if its initial frequency exceeds the critical 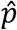. Figure 5 shows this threshold as a function of selfing rate *S* for a range of *r* values. A high ratio of invader-to-resident *k* lowers the threshold, as does a lower rate of selfing. But with selfing, even a very weak resident *Medea* can prevent invasion by a very strong *Medea* when the invader starts at low frequency (e.g., from a new mutation or rare migrant). And at high selfing rates, a resident *Medea* basically holds its position against all comers. Note that under obligate outcrossing (*S* = 0) and *r* ≠ 1, there is no intermediate equilibrium; the stronger *Medea* sweeps to fixation. This result shows that partial selfing flips the behavior of antagonistic *Medeas* from the Fisherian realm into the world of bistable dynamics (Barton and Turelli 2011).

**Figure 5.**
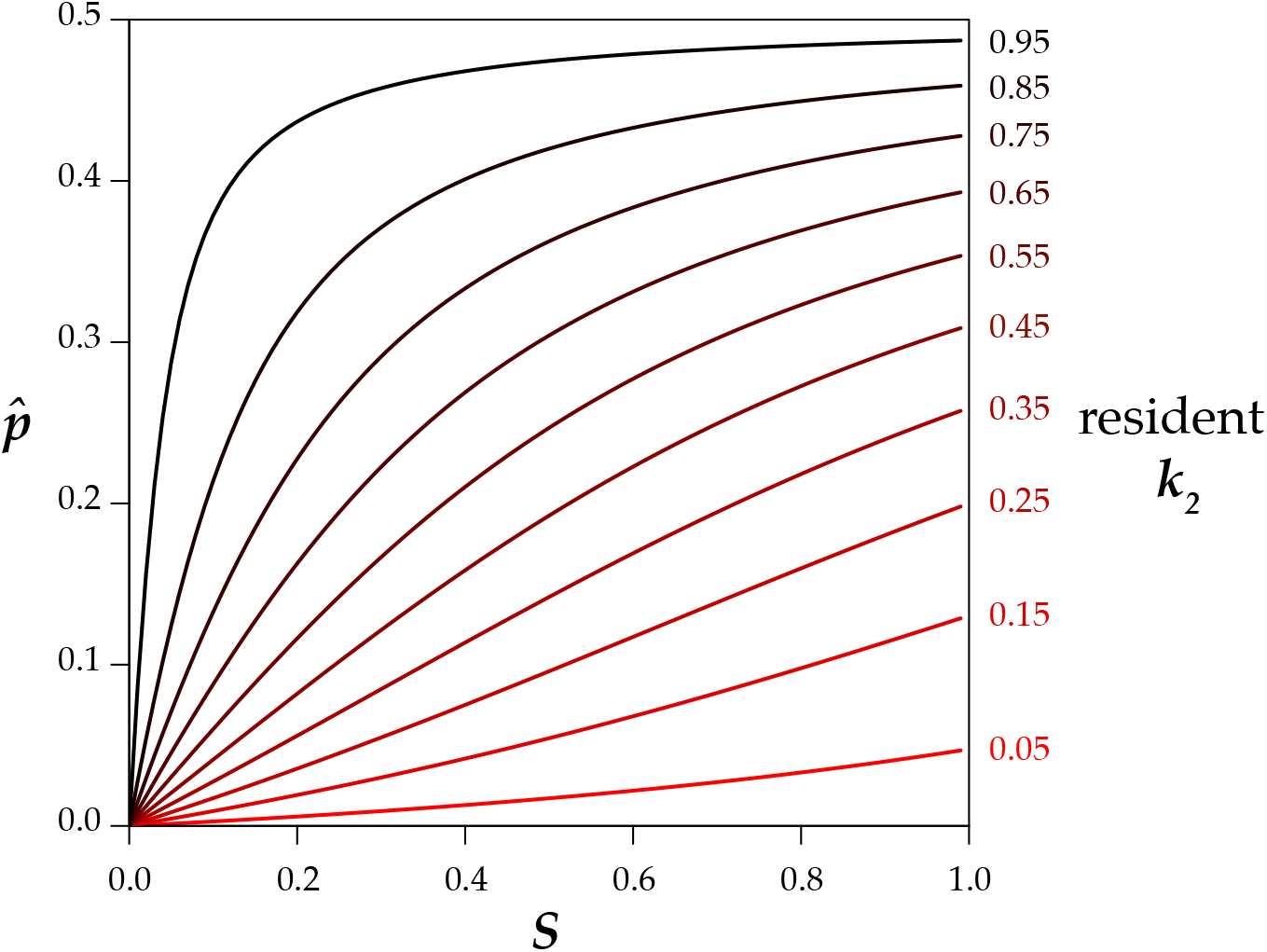
In a population with a resident *Medea* with the specified penetrance (*k*_*2*_), a completely penetrant *Medea* (*k*_*1*_=1) can invade and sweep to fixation only if its initial frequency *p* is above the relevant line at the indicated selfing rate (*S*). Below the line, the resident *Medea* excludes the invader. Note that the lines hold for any *k*_*1*_ and *k*_*2*_ with the same ratio; for example, the 0.25 line applies to any invader with four times the penetrance of a resident *Medea*.

Equation 5 allows for the calculation of the critical value of any one of *p, r*, or *S*, given values for the other two. If the frequency *p* of an invading allele with penetrance ratio *r* relative to the resident *Medea* is low enough that high values of selfing will prevent it from invading, the critical value *S*_*C*_ above which the *Medea* cannot invade is (Figure 4A)

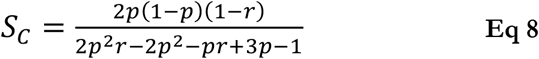

For example, if an invading *Medea* has a frequency of 0.01 and is fifty times more penetrant than the resident *Medea*, it will nevertheless be eliminated from the population if the selfing rate exceeds 0.67.

Similarly, a *Medea* with frequency *p* can invade a population with selfing rate *S* only if its penetrance relative to the resident *Medea* is greater than the critical value *r*_*C*_.

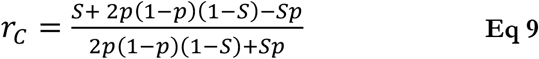

If an invading *Medea* has a frequency of 0.01 in a population with selfing rate *S* = 0.9, the invader will sweep only if it is more than 81 times more penetrant than the resident.

### Effect of androdioecy

Next, I will adapt the model to the idiosyncratic version of androdioecy observed in *Caenorhabditis* nematodes. In brief, hermaphrodites can self to make hermaphrodites, or they can mate with males to make a 50:50 mix of males and hermaphrodites, but hermaphrodites cannot mate with one another (Corsi *et al*. 2015). Hermaphrodites have an XX sex-chromosome complement, and so all self progeny are XX hermaphrodites. Males are X0, and their sperm are half X-bearing and half 0, lacking a sex chromosome. Consequently, cross progeny are half XX hermaphrodites and half X0 males. Males can also arise at low frequencies by X-chromosome nondisjunction in selfing hermaphrodites, but this occurs at a sufficiently low rate that we ignore its effects in the model. Overall, the genetic effect of this mating system is that hermaphrodites come from both selfing and outcrossing but males come exclusively from outcrossing, and males therefore tend to have much greater heterozygosity than hermaphrodites. We can accommodate this complication by allowing for sex-specific genotype frequencies, *X*_*h*_, *Y*_*h*_, and *Z*_*h*_ for the hermaphrodites and *X*_*m*_, *Y*_*m*_, and *Z*_*m*_ for the males.

Androdioecy introduces an additional complication – the two-fold cost of males (Cutter *et al*. 2019). If a hermaphrodite has a fixed brood size, outcrossing will produce half as many hermaphrodites as selfing will; the other half of the outcross brood will be male. Outcrossing therefore depletes the hermaphrodite population of heterozygosity even more than the mere presence of selfing would, by halving the contribution of each outcross to the hermaphrodite gene pool. In actual *Caenorhabditis* nematodes, selfing hermaphrodites do indeed have a fixed brood size – they are limited by the number of sperm that they make before the germline switches to oogenesis. But outcrossing removes that limit, as males can supply additional sperm, and can do so repeatedly. We address this issue by adding a new parameter, *b*, the ratio of hermaphrodite offspring from an outcross to hermaphrodite offspring from a self fertilization. If a hermaphrodite will have the same number of total offspring by selfing or by outcrossing, *b* is ½. If a hermaphrodite has twice as many offspring when it outcrosses, *b* is 1. From *C. elegans* data (WARD AND CARREL 1979; Hughes *et al*. 2007), *b* is probably close to 1, and biologically plausible values fall roughly in the range from ½ to 2.

Recursion equations for *p*_*h*_, *p*_*m*_, *Y*_*h*_, and *Y*_*m*_, derived and presented in the Supplementary Material, now have separate 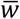. for hermaphrodites and males, as the relative frequencies of the male genotypes depend only on the outcrossing fraction of the population. Unlike the case under monoecy, the recursions are not identical for *Medea* and *Peel* elements, and antagonism between a *Medea* and a *Peel* introduces additional complications.

When all reproduction is by selfing, males are absent and androdioecy reduces to monoecy with *S*=1. When all reproduction is random mating, the mating system is dioecious and the recursion equations are the same as those for monoecy with *S*=0.

Superficially, the androdioecious equations are very similar to equation 3, as highlighted in table S1. As in the monoecious case, a single *Medea* or *peel* will sweep to fixation under androdioecy.

Androdioecy either accelerates or decelerates the spread and fixation of a *Medea* element as a function of the cost of males; when males are free (*b* = 1), androdioecy hastens the spread, but a two-fold cost of males (*b* = ½) slows it (Figure 6). The *b* value at the transition depends on the selfing rate.

**Figure 6.**
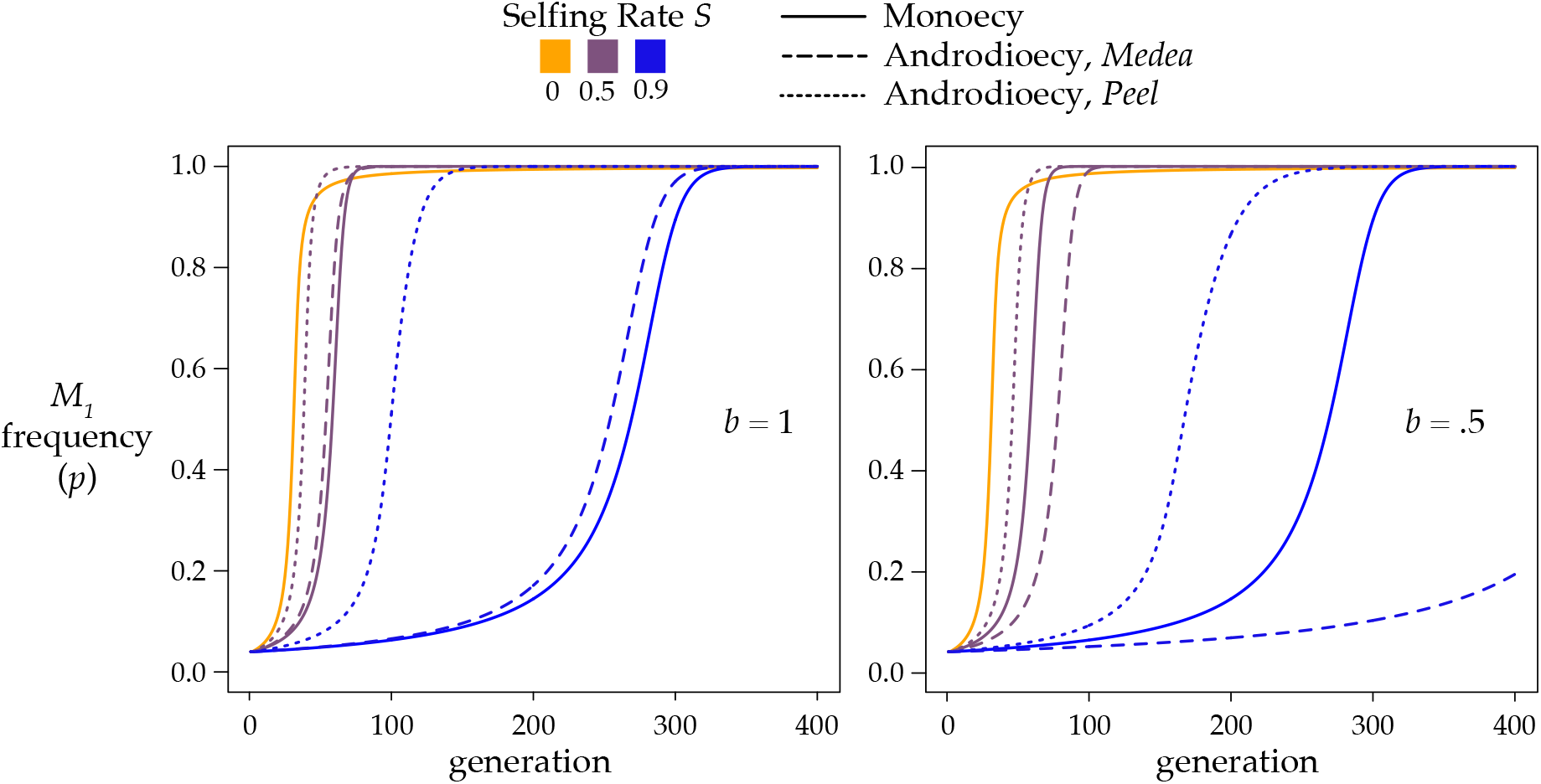
Androdioecy alters the rate of spread and fixation of *Medea* (dashed lines) and *peel* (dotted lines) elements. Under monoecy (solid lines, replotted from Figure 2), *Medea* and *peel* elements behave identically. In addition, only one line is plotted for *S*=0 because without selfing androdioecy and monoecy behave identically. This plot shows results for penetrance *k*_*1*_ = 1 and initial frequencies *p*_*h*_ = *p*_*m*_ =0.04, *Y*_*h*_ = *Y*_*m*_ = 0. The plotted *p* on the y-axis is the sex-weighted population average. The left panel shows the case of *b* = 1, where the number of hermaphrodite progeny from and mating is the same as the number from a self fertilization. In the right panel, *b* = 0.5, and mating events produce half as many hermaphrodites as selfings (the two-fold cost of males).

For *Medea* alleles, male heterozygosity *Y*_*m*_ turns out to play no role in the equations for allele frequencies, and so allele frequency dynamics can be described with just three variables. For models that include *peel* alleles, both *Y*_*h*_ and *Y*_*m*_ are required – hermaphrodite heterozygosity governs the effects of the *peels* under selfing, male heterozygosity under outcrossing – and so the allele frequency dynamics involve all four variables.

Under androdioecy, a *peel* element spreads more rapidly than a *Medea* element (Figure 6). With *peel* elements, male heterozygosity *Y*_*m*_ takes the place of hermaphrodite heterozygosity *Y*_*h*_ in the outcrossing components of the recursions. Because androdioecy causes *Y*_*m*_ to be greater than *Y*_*h*_ under partial selfing, *peel* elements are exposed to selection more than *Medea* elements.

In the monoecious model, antagonistic elements have an unstable internal equilibrium allele frequency 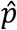 that depended only on selfing rate *S* and penetrance ratio *r*. All of that simplicity disappears under androdioecy. The equilibrium now also depends on the magnitudes of the two penetrances, *k*_1_ and *k*_2_, not just their ratio, on the cost of males, and on heterozygosity *Y*_*h*_ (and *Y*_*m*_ for models with *peel* elements). Despite the added complexity, the fundamental behavior of antagonistic *Medea* and *peel* alleles under androdioecy is the same as under monoecy: 0 and 1 are stable allele frequency equilibria, and partial selfing causes the system to exhibit positive frequency dependence.

Genotype frequency dynamics in this model are complicated, as male and hermaphrodite heterozygosities diverge from one another (Figure S1); in some cases male and female allele frequencies evolve transiently in opposite directions, before converging at the stable equilibria (loss or fixation). Selection generally keeps heterozygosities well above the extended Hardy-Weinberg expectations, even as the population evolves toward ultimate homogeneity.

I estimated 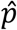 numerically for the biologically relevant case where individuals from a population fixed for one allele migrate into a population fixed for the alternate allele. For antagonistic *Medeas*, the threshold frequencies required for an allele to sweep to fixation differ only slightly from the monoecious case (Figure 7A).

In the case of antagonistic *peel* alleles, the critical frequency for a stronger allele to invade is somewhat lower than in the case of antagonistic *Medea* alleles (Figure 7B). In addition, 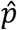 is no longer a monotonic function of selfing as it is in the other cases considered so far. When the invading allele is only slightly more penetrant than the resident, the critical frequency 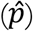 required for the allele to sweep is highest at an intermediate selfing rate, and declines again as selfing increases further.

*Medea-peel* antagonism introduces new surprises (Figure 7C, D): under androdioecy, a weaker *peel* can displace a stronger *Medea*! The *peel* alleles are strongly affected by male heterozygosity, which is always higher than hermaphrodite heterozygosity in this model. Consequently, for equally penetrant antagonistic alleles, the *peel* has an advantage, and the unstable equilibrium is no longer 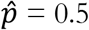.Nevertheless, the general behavior is preserved: under partial selfing, an invading allele, whether *peel* or *Medea*, must exceed a minimum frequency threshold in order to spread.

### Some relevant empirical data

Each of the three species of androdioecious *Caenorhabditis* carries blocks of hyperdivergent haplotypes, and the known *Medea* and *peel* alleles fall within these intervals (Seidel *et al*. 2008; BEN-DAVID *et al*. 2017; BEN-DAVID *et al*. 2021; Lee *et al*. 2021; Noble *et al*. 2021; Zdraljevic *et al*. 2024). For these blocks to be maintained by the model we have described, positive frequency-dependence due to antagonistic *Medea/peel* alleles, hyperdivergent regions should contain genes that could act as *Medeas* or *peels*. Such genes are necessarily expressed in the maternal or paternal germlines. As a simple test of plausibility, I asked whether hyperdivergent regions in *C. elegans* include germline-expressed genes. Comparable analyses for *C. briggsae* and *C. tropicalis* are not currently feasible due to the lack of analogous germline gene-expression datasets.

Genomic analysis of 1,384 *C. elegans* wild isolates from around the world (Cook *et al*. 2017; Lee *et al*. 2021) identified 312 hyperdivergent regions averaging 65kb in length and spanning 20.188 Mb of the reference genome (∼20% of the genome). Of these regions, 82% contain all or part of at least one protein-coding or lncRNA gene expressed in the germline of the reference strain, N2, based on the dataset of 9,345 such genes (Tzur *et al*. 2018).

Small RNAs – piRNAs, miRNAs, and siRNAs – play important roles in the maternal-zygotic transition (Quarato *et al*. 2021) and are demonstrably capable of acting as *Medea* toxins (Chen *et al*. 2007). When we add germline-expressed small RNAs (Billi *et al*. 2013; Bezler *et al*. 2019) to the analysis, the total fraction of hyperdivergent regions containing germline-expressed genes rises to 89%.

The two known elements active in the N2 strain, *Medea* locus *sup-35* and *peel* locus *peel-1*, occur, as expected, in the hermaphrodite-specific and male-specific subsets of the Tzur *et al*. dataset, respectively. When we restrict the analysis to these 3,803 sex-specific germline genes, we find that 54% of hyperdivergent regions contain such a gene. When we add sex-biased germline-expressed small RNAs (Billi *et al*. 2013; Bezler *et al*. 2019) to the analysis, the total fraction of hyperdivergent regions containing sex-biased germline-expressed genes rises to 63%.

The third known element in *C. elegans*, the *Medea mll-1* (*B0250*.*8*), recently discovered by (Zdraljevic *et al*. 2024), is a more complicated story. The reference strain (N2) carries an allele that lacks *Medea* activity, and, as expected, Tzur et al. (2018) did not find the gene to be germline expressed in N2. Strain XZ1516, which carries the *Medea* allele, does express it in the maternal germline, as required for its activity (Zdraljevic *et al*. 2024). Moreover, the reference strain N2 seems to express it in the germline too, but then the transcript is rapidly degraded through small RNA pathways. The hyperdivergent haplotypes that differentiate N2 and XZ1516 across this region contain other genes that appear in the Tzur et al. germline expression set, indicating that N2 also carries a haplotype with *Medea* potential.

## DISCUSSION

Despite the intuition that antagonistic *Medea* or *peel* elements should cause overdominant balancing selection, a simple model shows that they do not. Under obligate outcrossing, the stronger allele fixes. Under partial selfing, the system exhibits bistability: a common allele will drive a rare one to extinction, with the frequency threshold depending on the ratio of penetrances and the selfing rate.

What causes this positive frequency dependence under partial selfing? The situation is easiest to see in the case of equally penetrant *Medeas*. In this case, outcrossing causes no change in allele frequency at all, as first shown by Hedrick (1997). In the selfing part of the population, only the homozygous offspring of heterozygous mothers are affected by the *Medea*. With Mendelian segregation and equal penetrance, both classes of homozygous offspring are affected in equal numbers. However, the rare alleles present in the heterozygous mothers are a larger fraction of all rare alleles than the common alleles in these mothers are of the common alleles. The rare allele takes a much larger hit to its frequency when heterozygous mothers affect both homozygous classes equally. Meanwhile, outcrossing serves to generate the heterozygotes whose selfing then reduces the frequency of the rare allele.

Extension of the model to the idiosyncratic mating system of androdioecious worms reveals a number of new wrinkles, with maternal and paternal-effect elements now behaving differently and the frequency threshold now sensitive to additional parameters. Notably, *peel* elements drive more effectively than *Medeas*. The paternal-effect elements depend on male heterozygosity, which is higher than hermaphrodite heterozygosity because males are always the product of outcrosses. But the fundamental phenomenology is unchanged: the only stable equilibria are zero and one.

*Medea* and *peel* elements are reasonably modeled as toxin-antidote systems (Burga *et al*. 2020), and consequently they are often discussed in the context of other such systems, including spore killers, sperm killers, and pollen killers. It is therefore striking how different the population-genetic behavior of these systems is. Lopez-Hernandez et al (Lopez Hernandez *et al*. 2024) found that antagonistic spore killers at a single locus generate positive frequency dependence even under random mating, in contrast to our findings for elements that involve parental-zygotic interactions. Elements that kill sperm or pollen are sensitive to another variable, male gametic redundancy: a flower that kills half its pollen can nevertheless fertilize all its ovules with the remaining half. This phenomenon can cause drive to be more effective under partial selfing than under obligate outcrossing (Wang *et al*. 2024).

### Do *Medea/peel* elements cause selection for increased selfing?

As Noble et al. (2021) note, segregating *Medeas* impose a substantial fitness cost on the species, depressing the mean fitness. This creates a selection for mating-system modifiers, which can reduce the fitness costs of *Medeas* by reducing the frequency of heterozygotes. This concept is well established in the broader gene drive literature, and several studies have investigated the potential for invasion by alleles that increase inbreeding in the context of other (non-*Medea/peel*) gene-drive systems (BULL 2017; Bull *et al*. 2019 ; BEAGHTON AND BURT 2022). These studies find that alleles that favor inbreeding can invade under a range of plausible parameters. The fitness gains from suppressing drive, coupled with the elimination of the two-fold cost of males, provide the selective push to overcome the fitness cost of inbreeding depression.

These models could potentially extend to parental-zygotic *Medea/peel* systems. Most obligately outcrossing species of *Caenorhabditis* carry an enormous load of recessive deleterious alleles (Dolgin *et al*. 2007; BarriÈre *et al*. 2009; Fierst *et al*. 2015), and the magnitude of inbreeding depression may be sufficient to prevent invasion by alleles favoring selfing. But if ancestors of the androdioecious species had managed to reduce their levels of inbreeding depression, perhaps through an extended interval of small populations sizes, selection against *Medea/peel* elements could have helped in the origin of selfing. In addition, simultaneous segregation of antagonistic *Medea/peel* haplotypes at multiple genomic loci could impose such a large cost that even moderate levels of inbreeding depression could be overcome.

Among the three androdioecious *Caenorhabditis* species, where *Medea* and *peel* elements are known to occur, population genetic models of inbreeding modifiers as suppressors of drive face a difficulty: it is not clear what selective force would oppose increases in inbreeding. *C. elegans, C. tropicalis*, and *C. briggsae* occur in nature as completely inbred lines, having purged themselves of segregating recessive deleterious variants. Consequently, inbreeding depression does not provide a pressure that would oppose increased selfing rates, and a model along the lines already established will always favor increased selfing in the face of *Medea/peel* elements. Efforts to understand levels of outcrossing in these species will require consideration of benefits of recombination, including effects on the rate of adaptation and the amelioration of mutation accumulation (Cutter *et al*. 2019).

### Simplifying assumptions of these models

The models treated here consider a narrow slice of *Medea/peel* phenomena. Additional aspects of both molecular and population genetics provide opportunities for future work. Here I enumerate some of the model’s assumptions and review the outstanding questions they raise in relation to relevant empirical data.

### Molecular genetic assumptions

#### No recombination

We modeled a single locus with antagonistic alleles. If antagonistic elements are not truly allelic, and instead can recombine with one another, then selection will favor a recombinant haplotype that carries both driving alleles (BEN-DAVID *et al*. 2021). In the one case of antagonistic elements that have been mapped and validated at the level of molecular genes, the elements proved to be fully allelic (Pliota *et al*. 2024), and the general coincidence of drive elements and hyperdivergent regions in *Caenorhabditis* implies ample opportunity for antagonistic elements to occur within recombination-suppressed intervals. At the same time, *Medeas* occur at many separable loci within *Caenorhabditis* and *Triblium* genomes, both linked on the same chromosome and segregating independently on different chromosomes, and the behavior of such multilocus systems remains for future analysis.

#### No costs of the driving alleles

The model imposes no fitness cost on individuals that carry both toxin and antidote. It is easy to imagine that leaky toxin expression, or imperfect antidote efficacy, or even simple metabolic costs of transcription and translation, could be costly. Costs associated with drive elements are well known in segregating meiotic drive and gamete-killing systems such as mouse *t-* alleles and *Drosophila* segregation distorter (DUNN 1956; HARTL 1969), and are also very important for applications of engineered gene drive. Under obligate outcrossing, a *Medea* with a maternal fitness cost can be deterministically fixed, eliminated, or maintained at a stable equilibrium, depending on the nature and magnitude of the cost, the driver penetrance, and the initial allele frequencies (WADE AND BEEMAN 1994; Ward *et al*. 2011). Recently, (Long *et al*. 2023) found that monoecious partial selfing reduces the parameter space under which the polymorphic equilibrium is stable, such that driver costs are implausible as an explanation for the long-term maintenance of *Medea/peel* polymorphism in selfing *Caenorhabditis*.

Antagonistic *Medea/peel* elements with costs have not been studied, under partial selfing or otherwise. Such models would require many additional parameters and assumptions, including the costs of each allele, their dominance coefficients, and the question of whether the costs are born by mothers, fathers, or both, or by the offspring (Ward *et al*. 2011). For example, a leaky maternal-effect toxin would impose a fitness cost on mothers, while antidote translation costs would be born by zygotes independent of sex. The cell-biological details of these mechanisms make for an exceptional diversity of potential cost models (MADGWICK AND WOLF 2021).

But *Medea/peel* elements need not impose meaningful costs. For example, an engineered *Medea* in *Drosophila melanogaster* showed that no fitness cost through 20 generations of experimental competition (Chen *et al*. 2007). At present, there is no evidence that *Caenorhabditis Medea* or *peel* elements impose costs on their bearers, and indeed *peel-1* appears to be homozygous beneficial under laboratory conditions (Long *et al*. 2023).

#### No intragenic modifier mutations

The model ignores the molecular details by which the *Medea* or *peel* element acts and it does not allow for mutations that alter those details. In the molecularly characterized cases, each driving haplotype carries distinct molecular genes that act as toxin and antidote. Mutations that disrupt toxin function in a driving haplotype generate drive-resistant alleles, and these are known to segregate at low frequency in the *C. elegans peel-1* (*peel* toxin) and *sup-35* and *mll-1* (*Medea* toxin) genes (Seidel *et al*. 2008; Seidel *et al*. 2011; BEN-DAVID *et al*. 2017; Zdraljevic *et al*. 2024). Segregation of driving, drive-susceptible, and drive-resistant alleles can generate stable limit cycles under some assumptions (SMITH 1998), when the driver imposes a cost on individuals that carry it.

Mutations can also increase or decrease the penetrance of the toxin. Assuming that the linked antidote is capable of detoxifying an increased toxin effect, penetrance-increasing mutations should be able to invade and replace their own ancestral allele (THOMSON AND FELDMAN 1974; CROW 1991). While a locus is polymorphic for antagonistic elements, selection would favor penetrance-increasing mutations in each one.

The model assumes that each *Medea/peel* haplotype has an independent toxin-antidote mechanism, such that each allele’s antidote is powerless against the other allele’s toxin. An allele with a cross-reactive antidote would be able to invade and replace the alternate allele. An example of such a mutation is an insertion of a new toxin-antidote pair within an existing toxin-antidote locus; the two antidotes carried by the new haplotype give it resistance to the old and new toxins, and it can invade and replace the old haplotype. This scenario has been studied as a method for increasing the efficacy of engineered elements (Chen *et al*. 2007; Hay *et al*. 2010). Cross-active antidotes have also been studied in the spore-killing case (Lopez Hernandez *et al*. 2024).

Each *Medea* in *C. elegans, C. tropicalis* and *Tribolium castaneum* appears to have a completely independent toxin and antidote, with no antidotes that rescue the effects of unlinked toxins (Beeman *et al*. 1992; BEEMAN AND FRIESEN 1999; BEN-DAVID *et al*. 2017; BEN-DAVID *et al*. 2021; Noble *et al*. 2021; Zdraljevic *et al*. 2024). In *C. briggsae*, remarkably, a single *Medea* element occurs in different genomic loci in different wild isolates, the result of independent transposon-mediated insertions (Widen *et al*. 2023). In this case, the alleles in the non-reference location are pseudogenized; if they were functional, progeny of a mother heterozygous at one locus but homozygous *Medea* at a second would suffer no *Medea*-mediated killing, with the second locus zygotically protecting against the maternal effects of the first.

#### No sex linkage

The model treats autosomal *Medea/peel* elements. Sex-linked elements are not currently known in *Caenorhabditis* or elsewhere, but there is no obstacle to their occurrence. Hemizygosity affects *Medea* and *peel* elements differently, and as a function of which sex is heterogametic. In an XY or X0 male-heterogametic species, half of the sons of a mother heterozygous for an X-linked *Medea* will be hemizygous for the non-*Medea* allele, and so will suffer its effects no matter of the genotype of the father. On the other hand, *peel* drive is dependent on paternal heterozygosity, and an X-linked *peel* element would be incapable of driving at all in a dioecious male-heterogametic species. Under androdioecy, only the selfing part of the model would apply to an X-linked *peel*. Generally, for both *Medea* and *peel* elements, greater rates of selfing reduce the difference between autosomal and sex-linked elements. X-linked *Medeas* have been studied in the obligate outcrossing case (Ward *et al*. 2011; MARSHALL AND HAY 2012b), along with a whole Pantheon of X-, Y-, and Z-linked parental-zygotic synthetic drive systems: *Semele, Medusa*, inverse *Medea*, and others (MARSHALL AND HAY 2012b; MARSHALL AND HAY 2014; Verma *et al*. 2021).

#### No sex-specific effects

The model assumes both that embryos are affected by *Medea/peel* toxins in a sex-independent manner, although sex-dependent effects are easily imagined and have been studied for single *Medeas* in the obligate outcrossing case (MARSHALL AND HAY 2012b). In addition, the model here assumes that the penetrance of elements is the same whether they are transmitted by selfing or outcrossing. In fact, *peel-1* has higher penetrance when the toxin protein is passed to the embryo by a male than when passed by a hermaphrodite, probably due to differences in protein dosage between male and hermaphrodite sperm (Seidel *et al*. 2008; Seidel *et al*. 2011). As a consequence, the selfing and outcrossing components of the model should have different values for the penetrance parameters.

#### No cytoplasmic genetic and epigenetic modifiers

The genetics of *Medea* and *peel* elements involve interactions between parental and offspring genotypes, but recent studies have also revealed that the grandparental genotypes also matter. In *C. tropicalis, Medea*-heterozygous mothers from reciprocal crosses often have different effects on their offspring (Noble *et al*. 2021). In the best characterized case (Pliota *et al*. 2024), the pattern reflects an epigenetic parent-of-origin effect mediated by small RNAs.

### Population Biology Assumptions

#### No reproductive compensation

The model implies that matings that result in affected zygotes will yield fewer adult progeny than matings that do not. An alternative scenario is that the zygotes lost to *Medea* or *peel* activity will be replaced by additional siblings, so that each parent contributes the same number of children to the next generation’s gene pool. This pattern is known variously as reproductive compensation, local density regulation within families, and family-level soft selection (WADE AND BEEMAN 1994). One mechanism for reproductive compensation is that a family that loses offspring to a *Medea* has more resources for the remaining offspring, so losses that might otherwise occur due to density-dependent competition are ameliorated. The result, a transfer of resources from affected zygotes to their *Medea*-bearing siblings, increases the rate of spread of a single *Medea* under obligate outcrossing (WADE AND BEEMAN 1994; SMITH 1998). Though family-level soft selection is likely to be important in many cases, it seems unlikely to play a role in *Caenorhabditis* nematodes.

#### No group selection

Absent reproductive compensation, families affected by *Medeas* have smaller numbers of viable progeny than unaffected families do. This same pattern extends to any kind of population structure in which some groups have more descendants than other groups due to heritable factors that differ among them. For *Caenorhabditis* nematodes, which have very strong metapopulation dynamics, selection among groups is likely to be quite important. Small numbers of founder animals colonize ephemeral habitat patches, where they proliferate (Ferrari *et al*. 2017; Richaud *et al*. 2018; Sloat *et al*. 2022). When a patch reaches high densities, it generates dispersers to found new patches. Patches strongly affected by *Medeas* or *peels*, as a result of the genotype distribution among its founders, will have lower rates of proliferation and will take longer to generate dispersers. Simulations (Noble *et al*. 2021) show that the high burden of zygote loss imposed by antagonistic *Medeas* is sufficient to suppress patch growth and enhance the positive frequency-dependent selection against the rarer allele under androdioecy, or to induce this form of selection in the obligate outcrossing case, where it would otherwise not occur. Group selection models are sensitive to many details and further analysis is required (Bull *et al*. 2019).

*Infinite population size*. The model is deterministic and does not allow allele or genotype frequencies to change by drift. Wade and Beeman (1994) showed that the initial spread of a single *Medea* is slow, and consequently a rare *Medea* is very susceptible to loss by drift (WADE AND BEEMAN 1994; Hay *et al*. 2021). This risk is exacerbated by selfing. From our treatment of the monoecious single *Medea* case, Δ*p* approaches 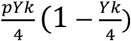 under selfing and 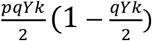 under outcrossing. When the *Medea* allele is rare, selfing approximately halves its rate of increase, extending it susceptibility to loss. At the same time, selfing decreases the effective population size, further magnifying the effects of drift (POLLAK 1987). As a consequence, low-frequency *Medea* or *peel* alleles in small populations with selfing behave as though they were neutral. Stochastic simulations in Noble *et al*. (2021) effectively illustrate this point (see their figure 10B).

While drift works against the deterministic spread of single *Medea* or *peel* alleles, it also allows invading antagonistic alleles to escape deterministic elimination. Allele frequencies bouncing around by drift can bounce across the unstable equilibrium (Barton and Turelli 2011). Stochastic fluctuations can allow otherwise prohibited transitions to proceed, and selfing facilitates the requisite fluctuations.

#### No spatial structure

Spatial structure, like drift, can cause local variation in genotype frequencies. In partially isolated patches, or on the leading edges of range expansions, *Medea* or *peel* alleles can exceed their threshold frequencies, particularly when the alleles have effects on the demographic properties of their local populations. This issue has been widely studied in the context of engineered gene drives (MARSHALL AND HAY 2012a; Bull *et al*. 2019; Champer *et al*. 2020; Dhole *et al*. 2020;

Champer *et al*. 2021; TURELLI AND BARTON 2022). For antagonistic *Medea* and *peel* alleles, selfing increases the potential for spatial effects to come into play; under obligate outcrossing, the stronger allele can always invade and sweep to fixation, independent of structure, but with partial selfing, the system flips to positive frequency dependence. In other words, an antagonistic allele can spread via pulled wave under obligate outcrossing but only by pushed wave under partial selfing.

The model results presented here are specific to the case of selfing, but the ephemeral, patchy population structure of obligately outcrossing *Caenorhabditis* induces biparental inbreeding that is likely to yield comparable patterns of positive frequency dependence (WADE 2000). When, imminently, *Medeas* are reported in obligately outcrossing *Caenorhabditis*, the results for monoecious partial selfing are likely to be more germane than those for obligate outcrossing.

*Fixed selfing rate*. We treat selfing rate as a fixed parameter of the model. Natural populations violate this assumption in two important ways. First, selfing rate varies heritably within populations and can evolve. *Medea* and *peel* alleles are expected to generate selection favoring increased selfing rates (BULL 2017; Drury *et al*. 2017; Bull *et al*. 2019; Noble *et al*. 2021). Both heritable variation and adaptive evolution of selfing rate are well documented in *Caenorhabditis* nematodes (Teotonio *et al*. 2006; Morran *et al*. 2009b; Teotonio *et al*. 2012; Noble *et al*. 2021). Variation in selfing rate is likely to be oligo- or polygenic in most species, and a full model of *Medea* dynamics would therefore have to incorporate linkage disequilibrium between the *Medea* and modifiers of the selfing rate. Second, selfing rates can be highly variable even in the absence of heritable variation. In particular, in androdioecious *Caenorhabditis*, a successful outcross event results in a transient increase in male frequency, purely as a side effect of the chromosomal sex-determination system. Transient elevations in male frequency can facilitate adaptive evolution, after which selfing rates can return to their previous levels (Morran *et al*. 2009a; Anderson *et al*. 2010; Cutter *et al*. 2019).

*Offspring are independent*. We assume that each outcross zygote draws its parents independently. This assumption would be violated by monogamy or similar mating system. Under some models of meiotic drive, polyandry reduces the efficacy of a driving allele by forcing sperm competition (HAIG AND BERGSTROM 1995). The criterion for such effects is that the father’s genotype at the drive locus affects the competitive ability of its sperm, which is probably quite typical for sperm-killing prezygotic drive mechanisms but is not presently known for *Medea* or *peel* alleles. For most drive models, polyandry affects dynamics but not equilibria (Verma *et al*. 2023).

A second violation of the random mating assumption would arise if individuals mate assortatively with respect to *Medea* genotype. Because *Medeas* and *peels* require heterozygosity to drive, positive assortative mating would reduce their efficacy.

Finally, the analysis of androdioecious models focused exclusively on cases where starting genotype frequencies were identical between males and hermaphrodites. This mimics the case of sex-independent migration, where individuals from a population fixed for one allele are introduced into a population fixed for the other. Models of gene drive engineered to modify populations often consider sex-limited introductions (e.g., transgenic male mosquitos), which would cause initial post-introduction outcrossing to be genotype-dependent; the introduced sex cannot mate with others of its own genotype. The effects of sex-limited introductions in androdioecious populations remains for future analysis.

### Do antagonistic *Medeas* explain ancient haplotypes in androdioecious *Caenorhabditis*?

This study was motivated by three observations: first, *Medea* and *peel* elements are abundant in *Caenorhabditis*, so abundant that they sometimes occur as the alternate alleles at a single locus. Second, *Caenorhabditis* genomes are mosaics of homogeneity and hyperdivergence. And third, the *Medea* and *peel* elements occur within the hyperdivergent regions. What explains this coincidence? There are three models (Figure 8).

**Figure 7.**
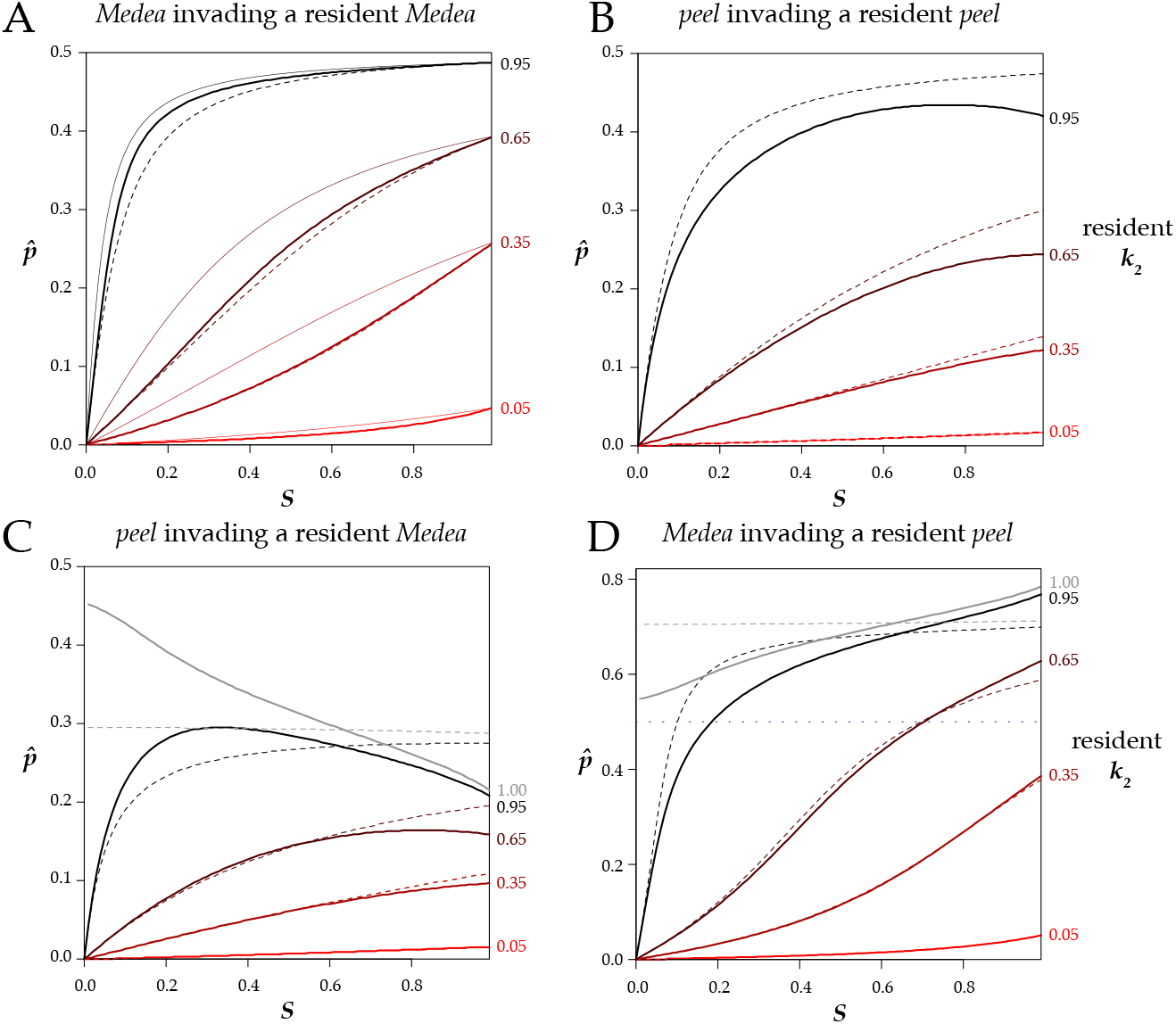
Frequency thresholds for invasion. **A**. In an androdioecious population with a resident *Medea* with the specified penetrance (*k*_*2*_), a completely penetrant *Medea* (*k*_*1*_=1) can invade and sweep to fixation if its frequency *p* is above the relevant thick solid line at the indicated selfing rate (*S*). Below the line, the resident *Medea* excludes the invader. The thin solid lines reproduce those in figure 5, showing results for a monoecious population. The dashed lines describe the results for an androdioecious population where the *k* values are ten-fold lower. For example, the topmost dashed line represents the case of *k*_*2*_ = 0.095 and *k*_*1*_ = 0.1. Under androdioecy, the absolute values of the *k*s matter, and not just their ratio as in monoecy. **B**. A *peel* allele invading a population with a resident *peel*. **C**. A *peel* allele invading a population with a resident *Medea*. The gray line at resident *k* = 1.00 shows that a *peel* allele can displace an equally penetrant (or even more penetrant) *Medea* allele under androdioecy. (For models that include only *Medea* or only *peel* alleles, the resident *k* = 1 line is a constant 0.5 and is not shown on the plots.) **D**. A *Medea* allele invading a population with a resident *peel*. Note that the y-axis has a larger range in this panel, as a fully penetrant invading *Medea* often requires an initial frequency above 0.5 (dotted line) to invade a population with a resident *peel*. The androdioecious curves are numerical estimates for the case of starting populations that consist exclusively of homozygotes, with *b* =1. Results for *b* = ½ are shown in Figure S2.

**Figure 8.**
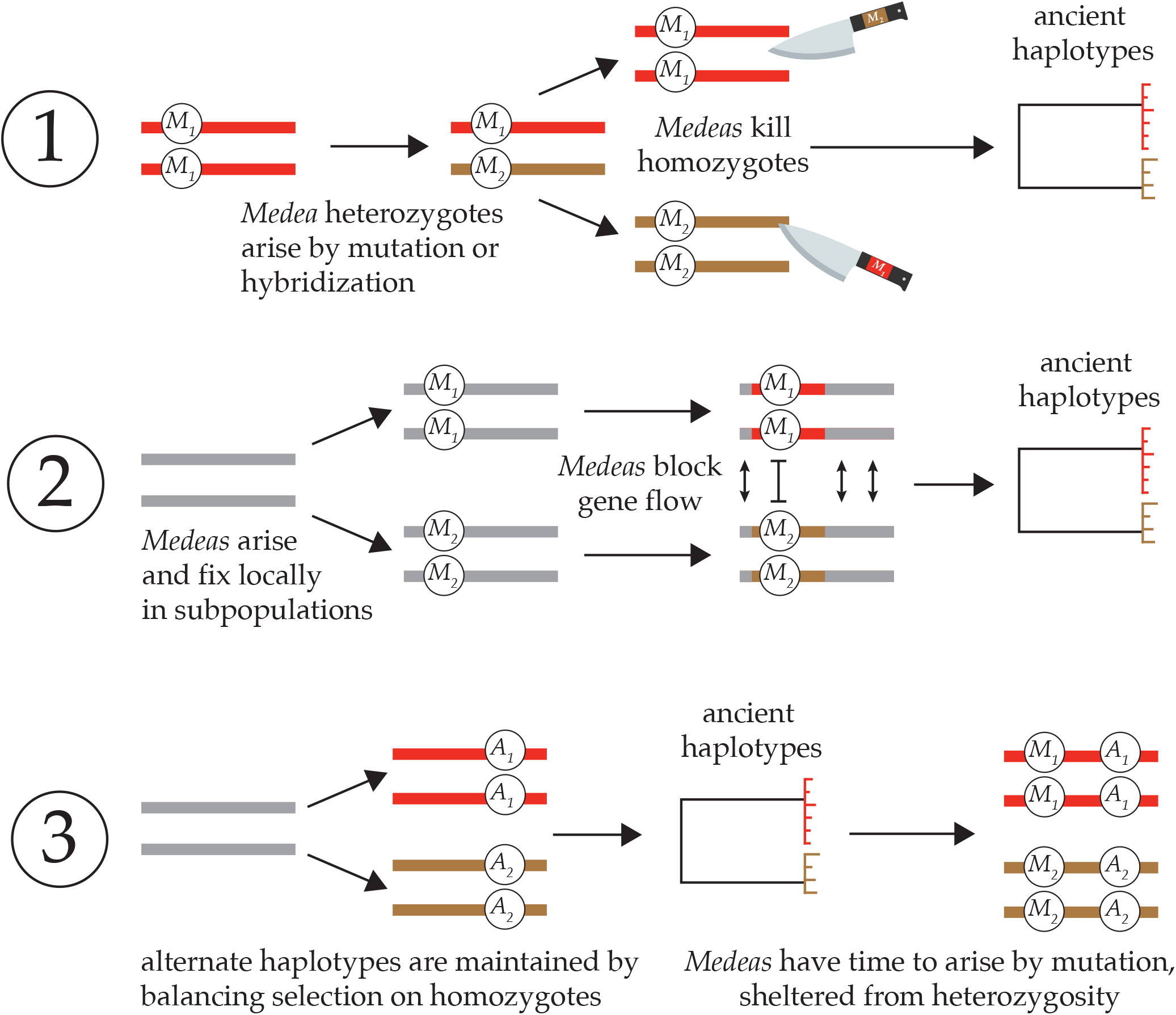
Three models for the occurrence of *Medea* elements on ancient haplotypes. 1. *Overdominance*. Alternate *Medeas* cause ancient haplotypes through *Medea*-mediated overdominant balancing selection. Genetic analysis rules this out. 2. *Positive Frequency Dependence*. Antagonistic *Medeas* cause ancient haplotypes by acting as a localized barrier to homogenizing gene flow between populations fixed for the alternative alleles. 3. *Incompatibility Trap*. Ancient haplotypes arise through ordinary ecological balancing selection that maintains the two alternate homozygotes; because of high selfing rates, heterozygotes are vanishingly rare. The long-term maintenance of alternative homozygotes provides a genomic environment in which *Medeas* can arise and fix neutrally, never exercising their gene-drive effects.

In the absence of any apparent costs to *Caenorhabditis Medeas* and *peels*, two models identify antagonistic *Medeas* as the cause of the hyperdivergence. The first, balancing selection via overdominance, is rejected by the results in this paper. The second is suggested by those results: *Medeas* cause ancient haplotypes via positive frequency dependence acting as a barrier to coalescence in a spatially structured population.

The basic conclusion from the analysis is that the alternate alleles exclude one another under partial selfing. Consequently, the occurrence of these alleles on ancient haplotypes may result from the local fixation of alternate alleles in separate populations, connected by gene flow. While most loci can move freely from one population to another, *Medea* alleles block one another from such movements. As a result, separated populations will become mosaics of genomic regions with shallow coalescence (non-*Medea*) and deep coalescence (antagonistic *Medeas*).

An important feature of the model results is that very weak elements can act as very effective obstacles. *C. elegans* clearly does not have highly penetrant *Medeas* in each of its hyperdivergent regions – only three elements have been identified genomewide in this species where we are best positioned to have discovered them. But the model does not require highly penetrant *Medeas*.

Elements that result in subtle developmental delays in a small fraction of offspring would almost certainly be invisible to our current experimental protocols. Instead, we would predict a general and diffuse outbreeding depression that manifests in the progeny of F_1_ animals, and which is congruent with the evidence (Dolgin *et al*. 2007), though many other models can also account for that pattern.

Another prediction is that hyperdivergent regions should carry genes expressed in the germline, capable of acting as parental-effect toxins. In the N2 strain, most hyperdivergent regions do carry such genes, but not all. These results suggest that antagonistic *Medea* or *peel* elements are unlikely to be the exclusive cause for hyperdivergent regions. If they were, we would have to invoke an explanation for the absence of germline-expressed genes in a subset of the hyperdivergent blocks. One explanation is imperfect discovery of germline-expressed genes. Another possible explanation is turnover of *Medea*/*peel* alleles through accumulation of disabling mutations. Under this model, some of the divergent haplotypes arose via antagonistic *Medea/peel* elements, but the germline toxin genes have become pseudogenized (at least in the reference strain N2). This scenario is documented for *peel-1*, where several wild isolates carry pseudogenized *peel-1* alleles as a result of point mutations (Seidel *et al*. 2008; Seidel *et al*. 2011), and apparently in both *mll-1* (Zdraljevic *et al*. 2024) and *sup-35* (BEN-DAVID *et al*. 2017), though the sequences of evolutionary events are less clear. Invasion of toxin-disabling mutations is expected once a *Medea* has reached high frequency, conditional on the toxin imposing a cost on the individual that expresses it (SMITH 1998). As noted above, there is not currently evidence that these elements impose a cost in *Caenorhabditis* or in a well-studied synthetic *Medea* in *Drosophila* (Chen *et al*. 2007). In addition, once a disabling mutation is present, the element will no longer act as a barrier to gene flow.

The simplest interpretation of the *C. elegans* germline gene expression data is that *Medea*/*peel* elements are not the primary cause of hyperdivergent regions. That leaves the third model for the coincidence of hyperdivergence and *Medeas*, which inverts the causality. In this incompatibility trap model (Seidel *et al*. 2008; Seidel *et al*. 2011), hyperdivergence comes first, and is the cause of the *Medeas*.

Ancient haplotypes are typically interpreted as evidence for long-term balancing selection mediated by ecological factors. This explanation is perfectly sufficient to explain the hyperdivergent haplotypes in *Caenorhabditis*, and indeed is supported by gene enrichment analyses (Lee *et al*. 2021). In selfing species, the balanced haplotypes are almost always present in homozygotes. Alleles with effects only in heterozygotes – *Medeas* and *peels* – could then gradually arise and evolve neutrally, rarely or never expressing their effects, prevented by recombination suppression from jumping to other haplotypes. This is the mirror image of the usual version of sheltered load – recessive alleles that evolve neutrally on haplotypes that are maintained in heterozygous states (Seidel *et al*. 2011). The incompatibility trap model is compatible with the emerging view that many apparent gene drive elements exhibit few or none of the expected population genetic signatures of gene drive, and may have evolved without relying on their segregation-distorting effects (Seidel *et al*. 2008; Sweigart *et al*. 2019; Long *et al*. 2023; Wang *et al*. 2024). Phenotypic effects of the *Medea* and *peel* loci unrelated to drive, as demonstrated for the case of *peel-1 in C. elegans*, could even be responsible for ecological balancing selection themselves, without the need for tight linkage to other loci under selection (Long *et al*. 2023).

Once antagonistic *Medea* and *peel* elements arise, they can reinforce the isolation of the divergent haplotypes via the positive frequency dependence described here. This model allows the *Medea/peel* elements to contribute to the accumulation of divergence but removes them from the driver’s seat.

### Implications of the models for engineered gene drives

We found that partial selfing slows the spread of *Medea* or *peel* elements and creates barriers to their spread in the presence of resident elements. These results are relevant to efforts to use engineered gene drive elements for public health applications. Though the major focus of such efforts has been gonochoristic species that vector diseases — mosquitoes most notably — there are several partial selfers that could be targets for engineered gene drives. The *Biomphalaria* snails that serve as intermediate hosts for *Schistosoma mansoni* fit the monoecious partial-selfing model (KENGNE-FOKAM *et al*. 2016), and most trematode and cestode parasites also have mixed mating systems. Our models show that mating system variation can cause *Medea* and *peel* to have different behaviors, and where possible the more potent driver should be selected. In addition, it may be useful to include mating-system modifiers into gene-drive constructs. *C. elegans* is a natural proving ground for such gene drive systems.

## Supporting information

MedeaFight R package and guide

## ACKNOWLEDGMENTS

This work was supported by a grant from the National Institutes of Health (GM141906). Thanks to Sarah Rankin and members of the Rockman Lab for helpful feedback.

## SUPPLEMENTARY FILES

### Supplementary text: Recursion equations under androdioecy

### R package *MedeaFight*

This zipped directory also contains an html guide to using the package and an R script, *PlotMedeaFigures*, that allows for reconstruction of the figures from the manuscript. To use the package, download and unzip the file and then in R, `install.packages`(“<path to directory> `/MedeaFightAndGuide/MedeaFight”, repos = NULL, type = “source”`). Finally, `library(“MedeaFight”)`. For some functions it is necessary to also have the Ternary package, which is available from CRAN, at cran.r-project.org.

## Supplementary text: Recursion equations under androdioecy

### Androdioecy, *Medea*

Taking into account the chromosomal sex determination mechanism in *Caenorhabditis*, each selfing results in hermaphrodite progeny only (XX), while each outcross results in 50% hermaphrodite, 50% male progeny. An outcrossing results in *b* hermaphrodite progeny for every 1 hermaphrodite that arises from a selfing; *b* is 0.5 if a cross results in the same number of progeny as a selfing, while it can be greater than 0.5 if a cross yields more progeny than a selfing.

Tracking the frequencies of different matings and the genotypes of their progeny, we can write *Medea* recursions in *X, Y, Z*, with *h* and *m* subscripts to specify hermaphrodite and male values respectively:

| Egg-parent | Sperm-parent | Frequency | Surviving Progeny Genotypes |  |  |  |  |  |
| --- | --- | --- | --- | --- | --- | --- | --- | --- |
|  |  |  | Hermaphrodites |  |  | Males |  |  |
| | | | $M_1M_1$ | $M_1M_2$ | $M_2M_2$ | $M_1M_1$ | $M_1M_2$ | $M_2M_2$ |
| | $M_1M_1$ | $SX_b$ | 1 | | | | | |
| | $M_1M_2$ | $SY_b$ | $(1-k_2)/4$ | $1/2$ | $(1-k_1)/4$ | | | |
| | $M_2M_2$ | $SZ_b$ | | | 1 | | | |
| $M_1M_1$ | $M_1M_1$ | $(1-S)X_bX_m$ | $b$ | | | $b$ | | |
| $M_1M_1$ | $M_1M_2$ | $(1-S)X_bY_m$ | $b/2$ | $b/2$ | | $b/2$ | $b/2$ | |
| $M_1M_1$ | $M_2M_2$ | $(1-S)X_bZ_m$ | | $b$ | | | $b$ | |
| $M_1M_2$ | $M_1M_1$ | $(1-S)Y_bX_m$ | $b(1-k_2)/2$ | $b/2$ | | $b(1-k_2)/2$ | $b/2$ | |
| $M_1M_2$ | $M_1M_2$ | $(1-S)Y_bY_m$ | $b(1-k_2)/4$ | $b/2$ | $b(1-k_1)/4$ | $b(1-k_2)/4$ | $b/2$ | $b(1-k_1)/4$ |
| $M_1M_2$ | $M_2M_2$ | $(1-S)Y_bZ_m$ | | $b/2$ | $b(1-k_1)/2$ | $b/2$ | $b(1-k_1)/2$ | |
| $M_2M_2$ | $M_1M_1$ | $(1-S)Z_bX_m$ | | $b$ | | | $b$ | |
| $M_2M_2$ | $M_1M_2$ | $(1-S)Z_bY_m$ | | $b/2$ | $b/2$ | | $b/2$ | $b/2$ |
| $M_2M_2$ | $M_2M_2$ | $(1-S)Z_bZ_m$ | | | $b$ | | | $b$ |

The recursions for hermaphrodites:

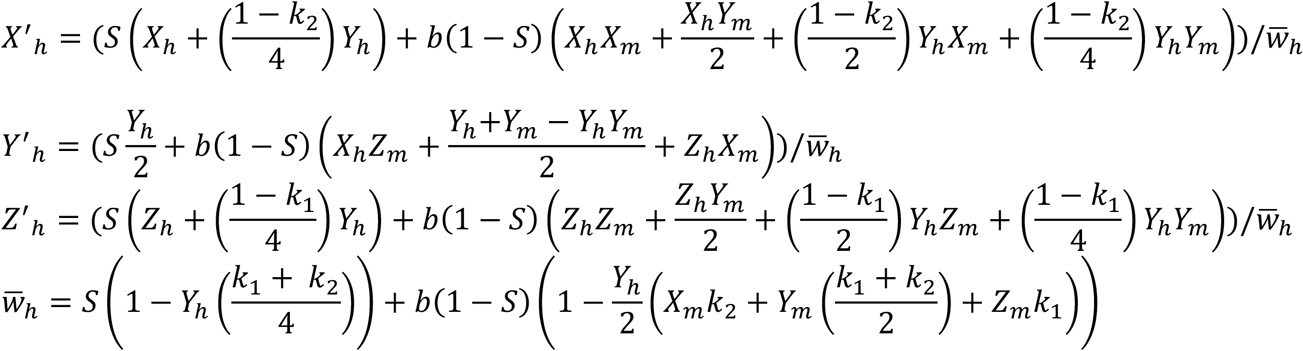

The recursions for males are simply the outcrossing part of the numerators above, normalized to the male-specific 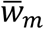, which is the outcrossing part of the denominator.

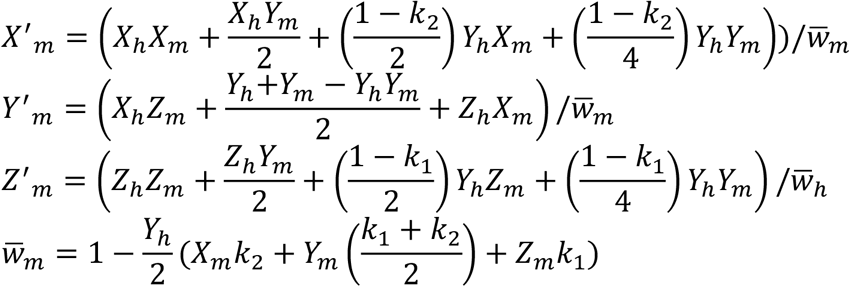

These equations can be consolidated to a system of equations for the four sex-specific allele frequencies *p* and heterozygosities *Y*:

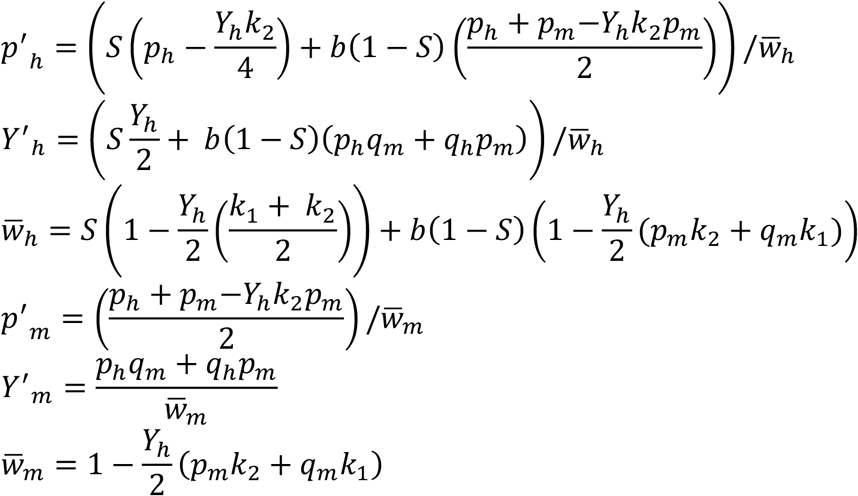

These recursions can be rewritten to highlight the effects of the *Medea* alleles, as shown in Table S1:

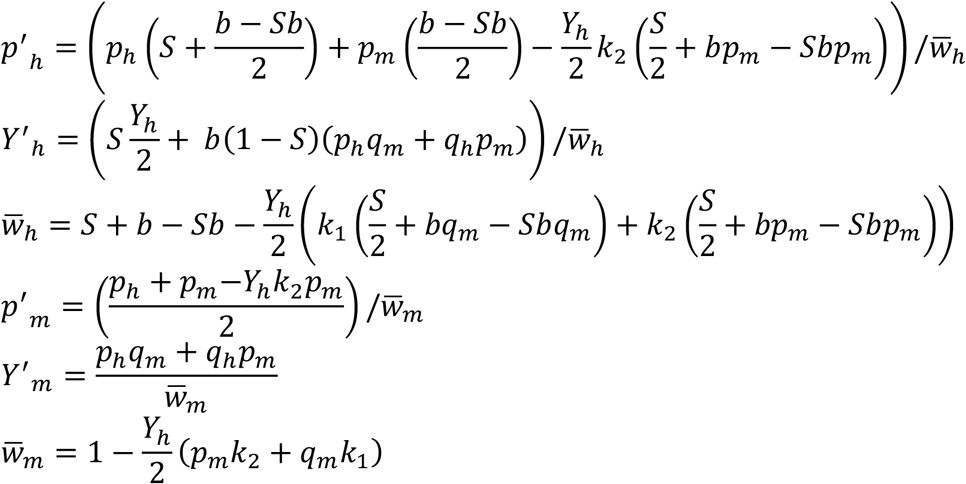

### Androdioecy, *peel*

The mating table below shows the effects of *peel* alleles (*P*_*1*_ and *P*_*2*_) in an androdioecious population.

| Egg-parent | Sperm-parent | Frequency | Surviving Progeny Genotypes |  |  |  |  |  |
| --- | --- | --- | --- | --- | --- | --- | --- | --- |
|  |  |  | Hermaphrodites |  |  | Males |  |  |
| | | | $P_1P_1$ | $P_1P_2$ | $P_2P_2$ | $P_1P_1$ | $P_1P_2$ | $P_2P_2$ |
| | $P_1P_1$ | $SX_b$ | 1 | | | | | |
| | $P_1P_2$ | $SY_b$ | $(1-k_2)/4$ | 1/2 | $(1-k_1)/4$ | | | |
| | $P_2P_2$ | $SZ_b$ | | | 1 | | | |
| $P_1P_1$ | $P_1P_1$ | $(1-S)X_bX_P$ | $b$ | | | $b$ | | |
| $P_1P_1$ | $P_1P_2$ | $(1-S)X_bY_P$ | $b(1-k_2)/2$ | $b/2$ | | $b(1-k_2)/2$ | $b/2$ | |
| $P_1P_1$ | $P_2P_2$ | $(1-S)X_bZ_P$ | | $b$ | | | $b$ | |
| $P_1P_2$ | $P_1P_1$ | $(1-S)Y_bX_P$ | $b/2$ | $b/2$ | | $b/2$ | $b/2$ | |
| $P_1P_2$ | $P_1P_2$ | $(1-S)Y_bY_P$ | $b(1-k_2)/4$ | $b/2$ | $b(1-k_1)/4$ | $b(1-k_2)/4$ | $b/2$ | $b(1-k_1)/4$ |
| $P_1P_2$ | $P_2P_2$ | $(1-S)Y_bZ_P$ | | $b/2$ | $b/2$ | | $b/2$ | $b/2$ |
| $P_2P_2$ | $P_1P_1$ | $(1-S)Z_bX_P$ | | $b$ | | | $b$ | |
| $P_2P_2$ | $P_1P_2$ | $(1-S)Z_bY_P$ | | $b/2$ | $b(1-k_1)/2$ | | $b/2$ | $b(1-k_1)/2$ |
| $P_2P_2$ | $P_2P_2$ | $(1-S)Z_bZ_P$ | | | $b$ | | | $b$ |

This table yields the following recursions for genotype frequencies:

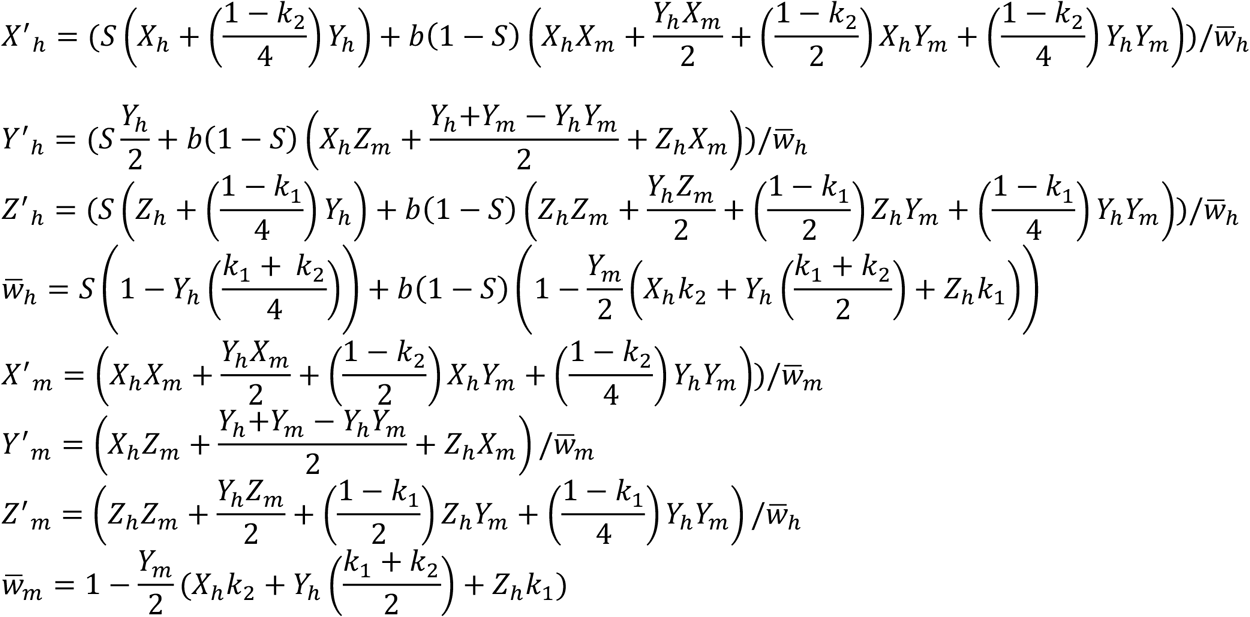

These equations can be consolidated to a system of equations for the four sex-specific allele frequencies *p* and heterozygosities *Y*:

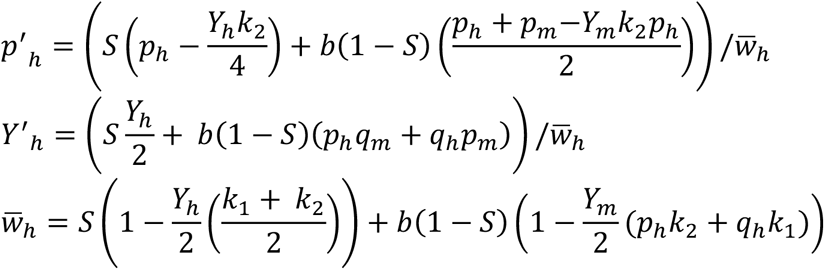

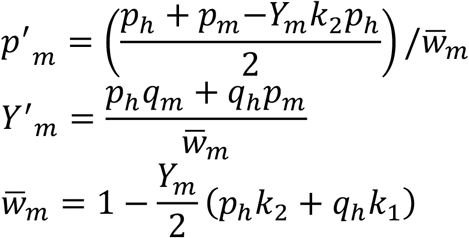

And these can be rewritten to highlight the effects of the *peel* alleles, as shown in Table S1:

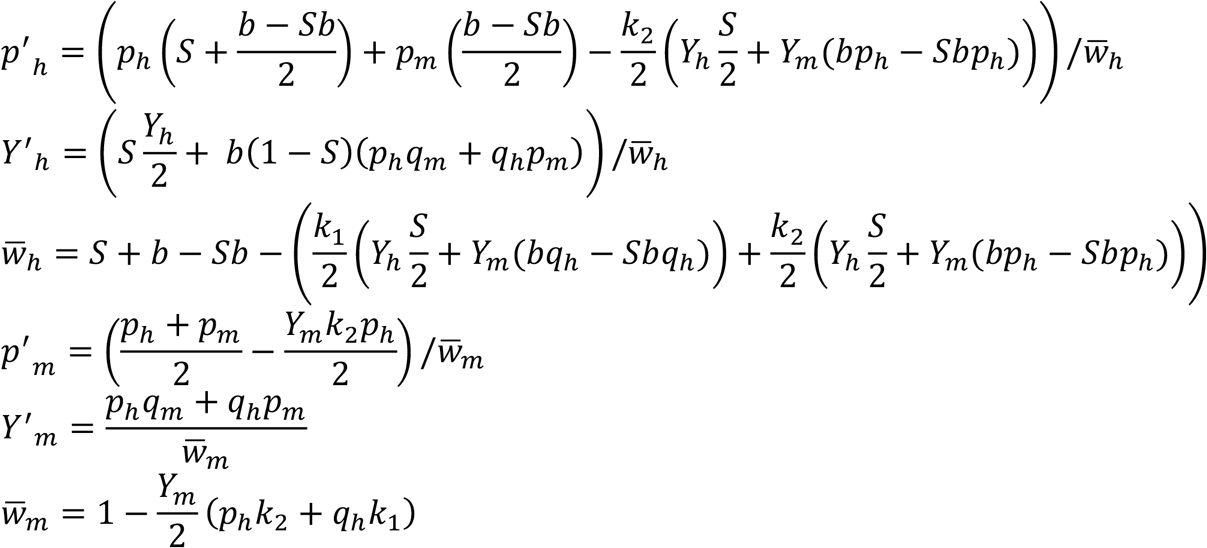

### Androdioecious *peel-Medea* Antagonism

The table below represents the case of antagonistic *Medea* and *peel* alleles (*M* and *P*) in an androdioecious population, where the penetrances of *Medea* and *peel* are *k*_*M*_ and *k*_*P*_ respectively. Note that this model is not symmetrical the way the other models are. Here we let *X, Y*, and *Z* be the genotype frequencies of *Medea* homozygotes, heterozygotes, and *peel* homozygotes, respectively.

Consequently, *p* is the *Medea* allele frequency and 1-*p* = *q* is the *peel* allele frequency.

| Herm | Male | Frequency | Surviving Progeny Genotypes |  |  |  |  |  |
| --- | --- | --- | --- | --- | --- | --- | --- | --- |
|  |  |  | Hermaphrodites |  |  | Males |  |  |
| | | | $M_1M_1$ | $M_1P_2$ | $P_2P_2$ | $M_1M_1$ | $M_1P_2$ | $P_2P_2$ |
| *MM* | | $SX_b$ | 1 | | | | | |
| *MP* | | $SY_b$ | $(1-k_P)/4$ | $1/2$ | $(1-k_M)/4$ | | | |
| *PP* | | $SZ_b$ | | | 1 | | | |
| *MM* | *MM* | $(1-S)X_bX_m$ | $b$ | | | $b$ | | |
| *MM* | *MP* | $(1-S)X_bY_m$ | $b(1-k_P)/2$ | $b/2$ | | $b(1-k_P)/2$ | $b/2$ | |
| *MM* | *PP* | $(1-S)X_bZ_m$ | | $b$ | | | $b$ | |
| *MP* | *MM* | $(1-S)Y_bX_m$ | $b/2$ | $b/2$ | | $b/2$ | $b/2$ | |
| *MP* | *MP* | $(1-S)Y_bY_m$ | $b(1-k_P)/4$ | $b/2$ | $b(1-k_M)/4$ | $b(1-k_P)/4$ | $b/2$ | $b(1-k_M)/4$ |
| *MP* | *PP* | $(1-S)Y_bZ_m$ | | $b/2$ | $b(1-k_M)/2$ | | $b/2$ | $b(1-k_M)/2$ |
| *PP* | *MM* | $(1-S)Z_bX_m$ | | $b$ | | | $b$ | |
| *PP* | *MP* | $(1-S)Z_bY_m$ | | $b/2$ | $b/2$ | | $b/2$ | $b/2$ |
| *PP* | *PP* | $(1-S)Z_bZ_m$ | | | $b$ | | | $b$ |

This table yields the following recursions for genotype frequencies:

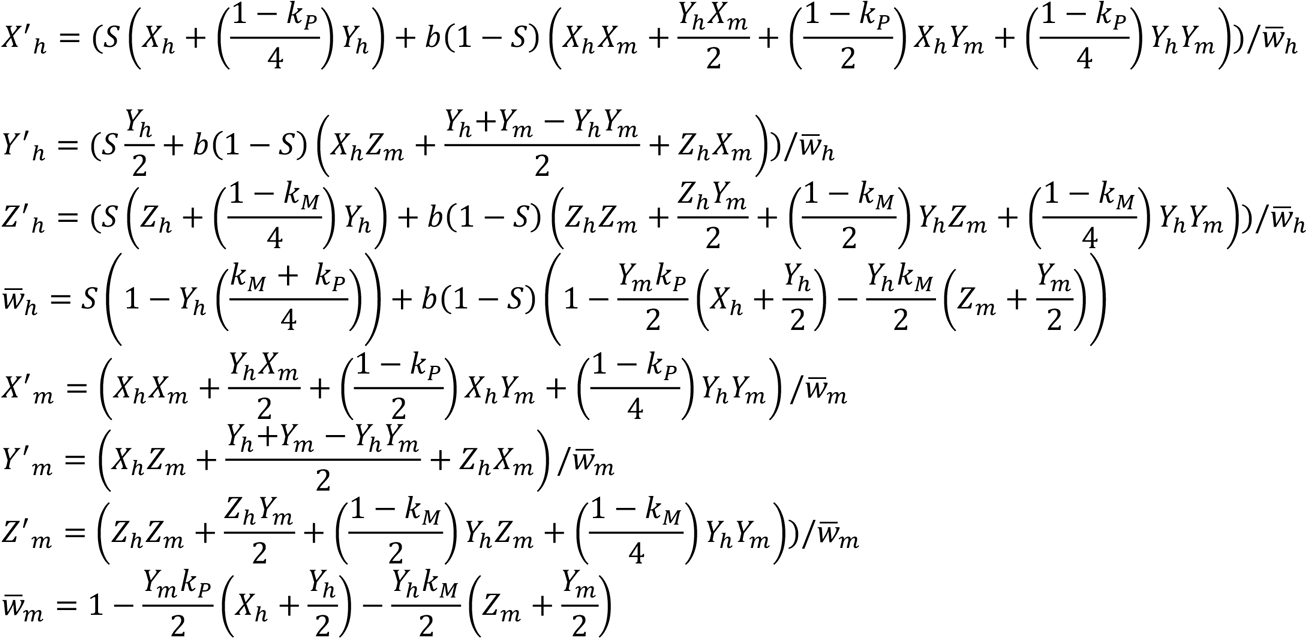

These equations can be consolidated to a system of equations for the four sex-specific allele frequencies *p* and heterozygosities *Y*:

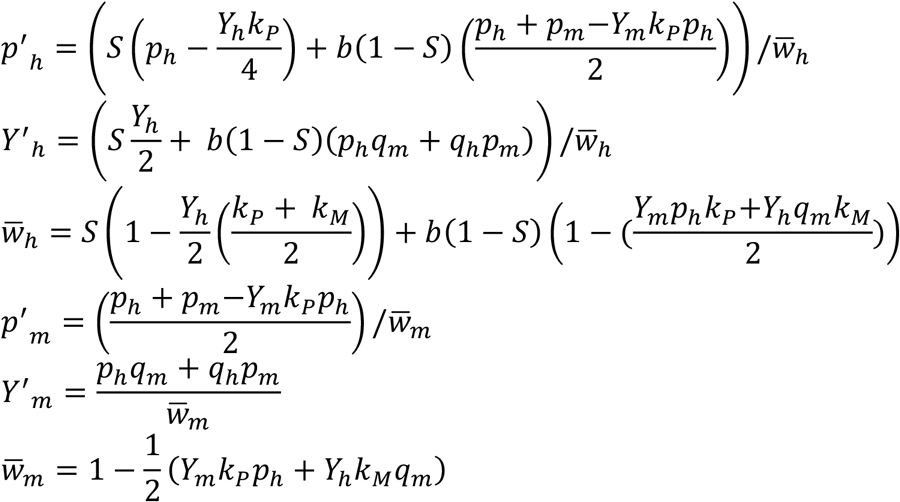

And these recursions can be rewritten to highlight the effects of the *Medea* and *peel* alleles, as shown in Table S1:

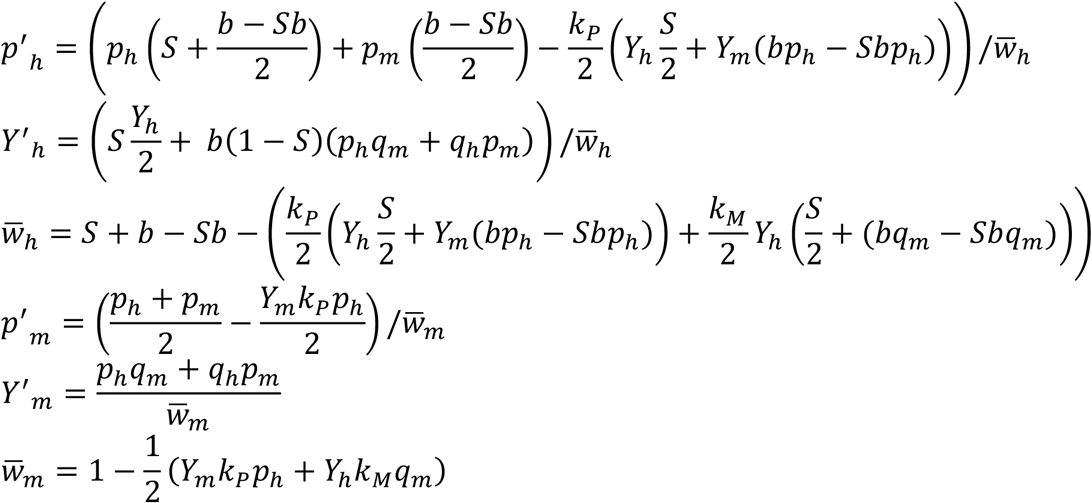

**Table S1.**
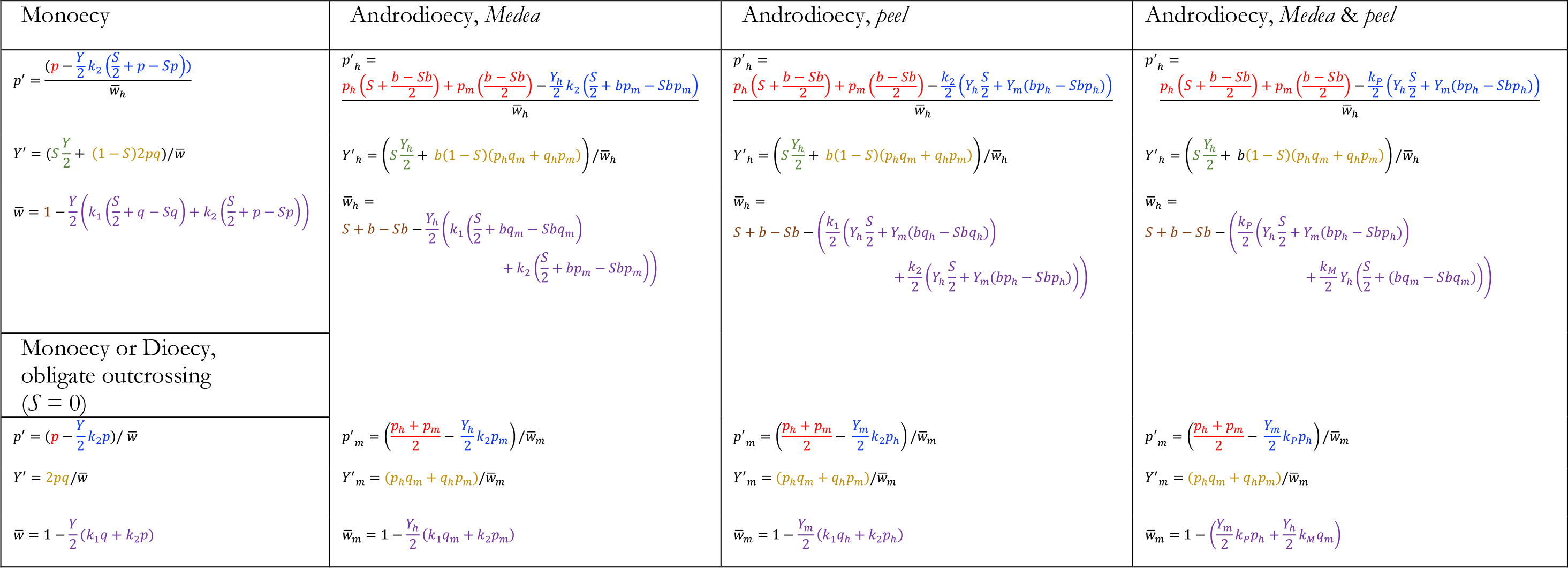
Comparison of recursion equations for different mating systems.Sex-specific recursion equations under androdioecy resemble the monoecious case for hermaphrodites and the obligately outcrossing case for males. Important differences are that the allele frequency *p* is replaced by its sex-weighted average in hermaphrodites and its unweighted average in males, and the generation of heterozygotes by outcrossing, 2*pq* under monoecy or dioecy, is *p*_*h*_*q*_*m*_ + *q*_*h*_*p*_*m*_ under androdioecy, influenced by sex differences in allele frequencies. Finally, *Medea* and *peel* elements differ in which sex’s heterozygosity and allele frequency influence the effects of the alleles. **Variables and parameters:** *p M*_*1*_ allele frequency *q M*_*2*_ allele frequency, 1-*p* *Y M*_*1*_*M*_*2*_ heterozygote genotype frequency 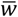 Population mean fitness, the proportion of zygotes that survive to reproduce *S* Selfing rate *b* Ratio of hermaphrodite progeny from an outcrossing to hermaphrodite progeny from a selfing *k*_*1*_ penetrance of the *M*_*1*_ allele *k*_*2*_ penetrance of the *M*_*2*_ allele *k*_*M*_ penetrance of the *Medea* allele in the *Medea-peel* antagonism model *k*_*P*_ penetrance of the *peel* allele in the *Medea-peel* antagonism model *h* subscript for hermaphrodite-specific variables *m* subscript for male-specific variables

**Figure S1.**
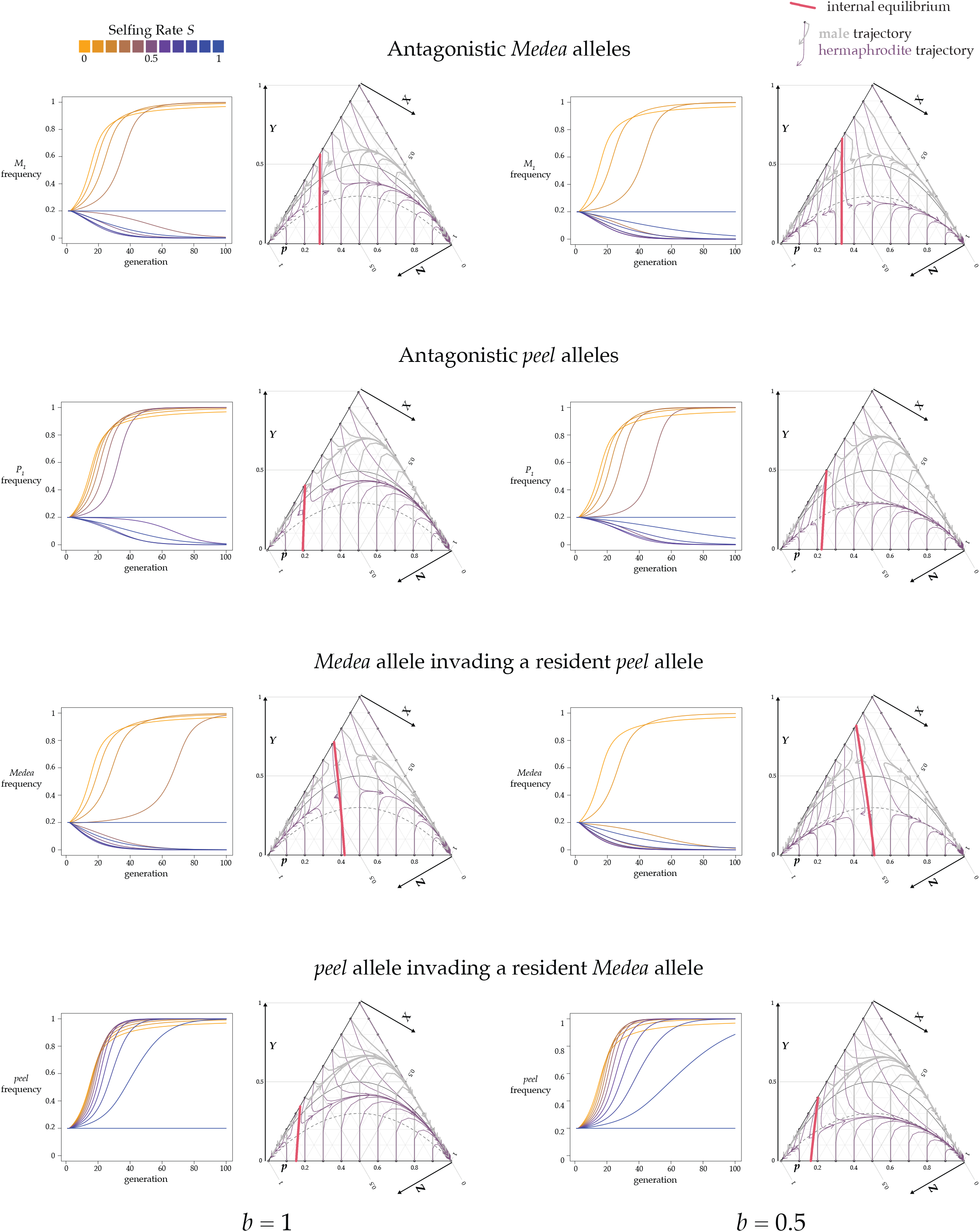
The effects of *Caenorhabditis*-type androdioecy on genotype and allele-frequency evolution differ between *Medea* and *peel* alleles, and are driven by large differences in heterozygosity between males and hermaphrodites. The four rows show the dynamics for antagonistic *Medeas*, antagonistic peels, a *Medea* invading a population with a resident *peel*, and a *peel* invading a population with a resident *Medea*. The left half of the figure shows results for *b* =1, and the right half *b* = 0.5. For each of the eight situations, there are two plots. The left one shows allele frequencies found by iterating the relevant equations (Supplementary File 1) for eleven values of *S* from 0 to 1, with initial *M*_*1*_ frequency *p* = 0.2 and heterozygosity *Y* = 0. The resident *Medea M*_*2*_ has penetrance *k*_*2*_ = 0.65 and the invading *Medea M*_*1*_ has penetrance *k*_*1*_ = 1. The right panel in each pair shows the dynamics in genotype space via De Finetti plot. These examples show again the case of *k*_*1*_ = 1 and *k*_*2*_ = 0.65, here with fixed selfing rate *S* = 0.57. Each trajectory starts from one position at the periphery of the plot and represents the male- or hermaphrodite-specific genotype frequencies through 15 generations. The red line shows the unstable internal equilibrium, conditional on identical starting frequencies for males and females. Under androdioecy, the equilibrium is sensitive to the initial heterozygosity (as seen in its departures from the vertical). To the right of this line, the invading allele fixes, and to the left, it is eliminated. The solid black curve shows genotype frequencies at Hardy-Weinberg equilibrium, and the dashed curve shows genotype frequencies under the neutral extended Hardy Weinberg equilibrium with selfing.

**Figure S2.**
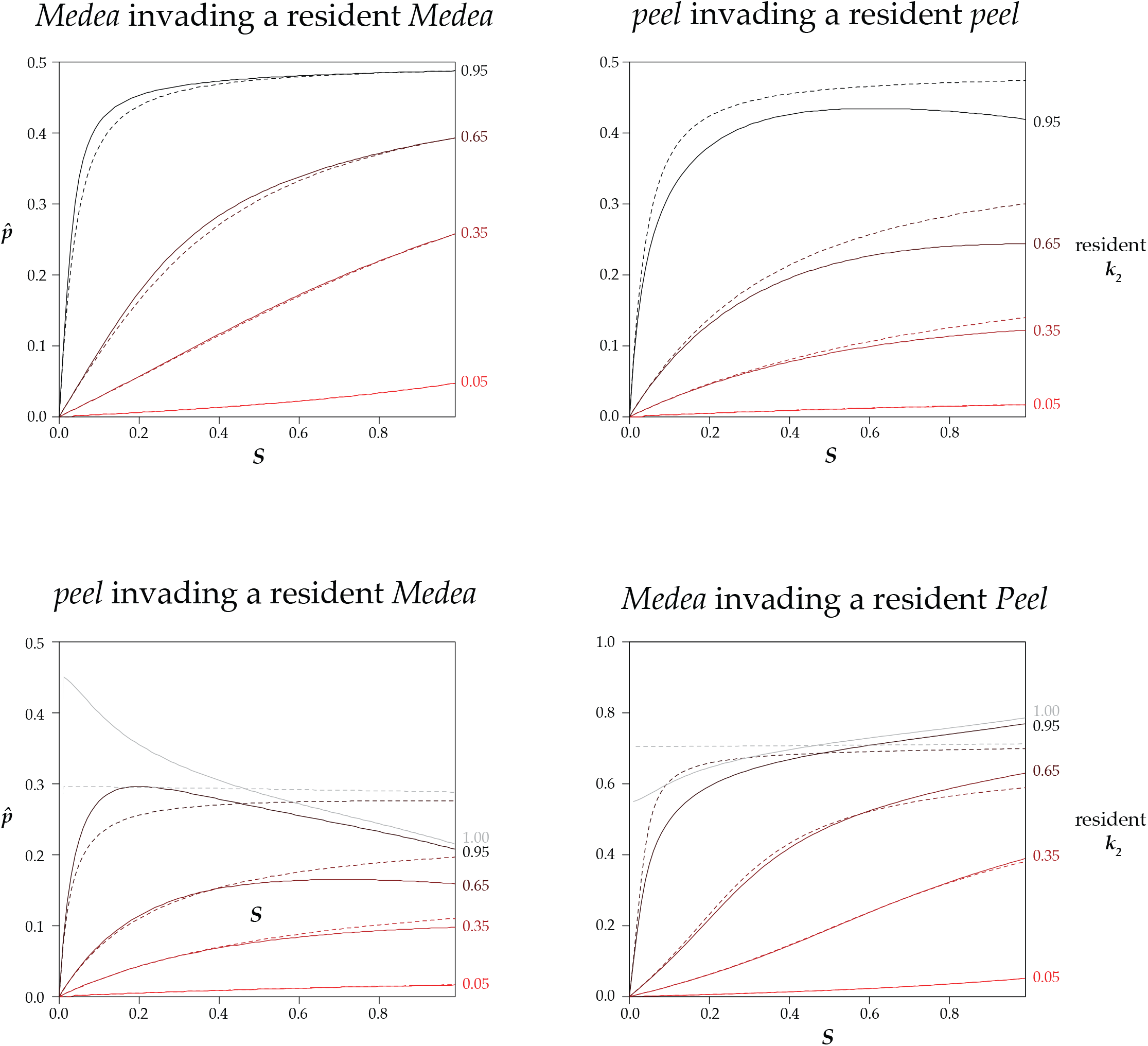
Frequency thresholds for invasion, with *b* = 0.5. This figure differs from Figure 5 only in the value of *b*. In an androdioecious population with a resident allele with the specified penetrance (*k*_*2*_), a completely penetrant allele (*k*_*1*_=1) can invade and sweep to fixation if its frequency *p* is above the relevant thick solid line at the indicated selfing rate (*S*). Below the line, the resident excludes the invader. The dashed lines describe the results for an androdioecious population where the *k* values are ten-fold lower. For example, the topmost dashed line represents the case of *k*_*2*_ = 0.095 and *k*_*1*_ = 0.1.

