## Supplementary material for "Parental-effect gene-drive elements under partial selfing, or why do *Caenorhabditis* genomes have hyperdivergent regions?": MedeaFight R package and guide: HowToUseMedeaFight.html

Using the MedeaFight Package


### Using the `MedeaFight` Package

```
library(MedeaFight)
```

Use this package to study the evolution of *Medea* and *peel* alleles under different kinds of mating systems, with the possibility that both alleles at a single locus harbor *Medea* and *peel* activity.

The package accompanies a paper on this topic. You can find the latest version of the paper at https://rockmanlab.bio.nyu.edu/publications/.

The package is written in base R and has not undergone extensive testing in Rstudio.

### Recursions

Most of the functions in the package use recursion equations to find genotype frequencies at some future generation given a set of initial genotype frequencies. To see a typical example, try `?AndroMedeaXYZ`. Some of the functions track the genotype frequencies, *X, Y,* and *Z*, while others track only the allele frequency *p* and heterozygosity *Y* (*e.g.*, `MedeaYp`). These functions use different systems of equations, but they are completely interconvertible: *p* is just *X + Y*/2. Each function requires a set of starting frequencies, a selfing rate (which can be any value from 0 to 1), and penetrances for the *Medea/peel* activity of each allele.

#### Mating Systems

There are recursions for two general classes of mating system: monoecy and androdioecy. The monoecious recursion equations, and their derivations, are in the main text of the paper (equations 1-3). The monoecious model imagines a population of hermaphroditic organisms that are capable of self fertilization and also cross fertilization. Each zygote arises from a selfing with probability *S* and an outcross with probability 1-*S*.

The androdioecious model imagines a *Caenorhabditis elegans*-type mating system (the equations are in the supplement to the paper). Hermaphrodites can self but they cannot cross with one another. A second sex, males, allows for crosses. Selfing results in hermaphrodite progeny, while crossing results in a 50:50 mix of hermaphrodites and males. Androdioecious models need to separately track genotype frequencies in hermaphrodites and males.

Monoecious functions (`MedeaYp` and `MedeaXYZ`) return single values each generation for allele and genotype frequencies and population mean fitness, while the androdioecious functions return two values for each: one for hermaphrodites and one for males.

Here’s an example where we’ll track genotype frequencies through 100 generations. The two alleles start with frequencies 0.05 and 0.95, all present in homozygotes. The first allele – the one at low frequency – has a killing penetrance of 0.8. That is, of the common-allele homozygotes that are offspring of a heterozygous mother, 80% will die. The other allele – the common one – has a lower killing penetrance, only 0.4. The selfing rate is 0.4. Will the rare allele increase or decrease in frequency?

```
library(MedeaFight)
MedeaXYZ(start = c(0.05, 0, 0.95), selfing = 0.4, k1 = 0.8, k2 = 0.4, n.gens = 100)[c(1,100),]
#>                X            Y         Z            p         W
#> 1   5.000000e-02 0.0000000000 0.9500000 0.0500000000 1.0000000
#> 100 1.859314e-05 0.0001656036 0.9998158 0.0001013949 0.9999404
```

This shows that the rare allele decreased in frequency, even though it has a higher penetrance. Try it again with a lower selfing rate, say, `selfing = 0.1`.

```
MedeaXYZ(start = c(0.05, 0, 0.95), selfing = 0.1, k1 = 0.8, k2 = 0.4, n.gens = 100)[c(1,100),]
#>               X          Y         Z          p         W
#> 1   0.050000000 0.00000000 0.9500000 0.05000000 1.0000000
#> 100 0.003384546 0.09593089 0.9006846 0.05134999 0.9634736
```

With less selfing, the rare but penetrant allele increases in frequency.

#### Allele Types

Under androdioecy, *Medea* and *peel* alleles behave differently (they are totally equivalent to one another under both monoecy and dioecy, curiously). The package includes separate functions for four different cases under androdioecy: both alleles at the locus are *Medea*s (`AndroMedeaYp` and `AndroMedeaXYZ`), both alleles are *peel*s (`AndroPeelYp` and `AndroPeelXYZ`), the allele of interest is a *Medea* and the alternate allele is *peel* (`MedPeelYp` and `MedPeelXYZ`), or vice versa (`PeelMedYp` and `PeelMedXYZ`).

### Plotting

Many functions in the package make it easy to visualize evolution of *Medea* and *peel* alleles through plots.

#### Dynamics

Two functions, `FreqByTimePlot` and `DeFinetti`, plot allele or genotype frequencies through time. `FreqByTimePlot` is good for seeing the effect of variation in selfing rate, holding all else constant:

```
FreqByTimePlot(start = c(0.2, 0), k1 = 1, k2 = 0.5, S = seq(0, 1, 0.1), weights = 2, mod = MedeaYp, plotgens = 100)
```

As in the paper, the colors represent the selfing rate, from 0 (orange) to 1 (blue). With these parameter values, under the monoecious model, the less common allele sweeps to fixation under low selfing but is swept to extinction with higher selfing.

Note that we specified the start frequencies with a vector of two numbers, (p, Y), in this case, instead of three (X, Y, Z). because we used the MedeaYp model. `FreqByTimePlot` expects a model parameterized in p and Y.

DeFinetti plots represent genotypes (X, Y, Z) in a triangular state-space where the three frequencies have to add up to 1. The `DeFinetti` function therefore requires a model in XYZ. The function is good for looking at how evolution depends on initial frequencies, holding all else constant.

```
DeFinetti(startingpoints = "Boundary", k1 = 1, k2 = 0.65, S = 0.6, sexes = "both", plotgens = 15, weights = 2, mod = MedeaXYZ)
```

The horizontal axis represents the allele frequency, *p*, and the vertical axis is the heterozygote frequency, *Y*. The curved lines represent the expected Hardy-Weinberg equilibrium genotype frequencies, with (dashed) or without (solid) selfing. What we see is that, for the parameters specified, genotype frequencies evolve to *p* = 0 or *p* = 1, depending on the starting frequencies, and that the trajectories follow a manifold with higher heterozygosity that we’d see absent selection.

#### Equilibria

What is the initial frequency above which the allele frequency increases, and below which it decreases? For the monoecious model, we can find it using `AddTheoryEquilibrium`. This will add the equilibrium for p and Y to the plot, and also output them to a list.

```
DeFinetti(startingpoints = "Boundary", k1 = 1, k2 = 0.65, S = 0.6, sexes = "both", plotgens = 15, weights = 2, mod = MedeaXYZ)
hats <- AddTheoryEquilibrium(k1 = 1, k2 = 0.65, S = 0.6)
```

```
hats$p_hat
#> [1] 0.3312831
hats$Y_hat
#> [1] 0.3123028
```

For the androdioecious models, there is no simple formula for the equilibrium values, but they can be found numerically, using the very slow `AddNumericEquilibrium` function.

Finally, we can plot the equilibrium allele frequency, above which the allele increases and below which it decreases, as a function of selfing rate. For the monoecious model, we can use the analytical results from the paper.

```
thresholds <- TheoryPhat(residents = c(0.1,0.5,0.9))
```

Here we see that the equilibrium value of *p* is higher at higher rates of selfing, and that it varies as a function of the relative penetrances. In this case, we’re looking at resident alleles that are 10%, 50%, and 90% as penetrant as the invading allele. These values are represented by color, from red (invader much more penetrant) to black (invader only slightly more penetrant).

Under androdioecy, these equilibria have to be found numerically, which is slow. But `NumericPhat` will do it.

### Summary

*Medea* and *peel* alleles have unintuitive behaviors under partial selfing. Playing around with the relevant parameters - starting frequencies, penetrances, selfing rates, allele types, and mating systems - makes it possible to develop some intuitions.
