## Supplementary material for "Parental-effect gene-drive elements under partial selfing, or why do *Caenorhabditis* genomes have hyperdivergent regions?": MedeaFight R package and guide: DESCRIPTION

Package: MedeaFight
Title: Model the evolution of Medea and peel alleles in populations of partial selfers
Version: 1.0.0
Authors@R: 
           comment = c(ORCID = "0000-0001-6492-8906"))
Description: Track Medea and peel allele and genotype frequencies through time under monoecy, dioecy, and Caenorhabditis-type androdioecy with selfing. The package also includes functions for plotting frequencies through time and calculating or estimating frequency equilibria and plotting those. 
License: `use_mit_license()`
Imports:
	Ternary (>= 2.1.2)
Encoding: UTF-8
Roxygen: list(markdown = TRUE)
RoxygenNote: 7.3.2.9000
