## Supplementary material for "Parental-effect gene-drive elements under partial selfing, or why do *Caenorhabditis* genomes have hyperdivergent regions?": MedeaFight R package and guide: FunctionsForPackage.R

```
#' TheoryPhat
#'
#' @description
#' `TheoryPhat` plots the value of p_hat, the unstable internal equilibrium for antagonistic Medeas or peels,
#' under monoecy, as a function of selfing rate, for one or more ratios of the penetrances of the alleles.
#'
#' @details
#' This simply calculates the value of p_hat from equation 5 of Rockman 2024 and plots it as a function 
#' of selfing rate. It does this for each ratio of resident/invading Medea. Note that under monoecy, the
#' equilibrium is identical whether we're dealing with Medeas or peels or both. This function calculates
#' p_hat every 0.01 from 0 to 1 and connects the values with a line.
#' 
#' 
#' @param residents This is a vector of values for the penetrance of the resident Medea, relative to an 
#' invading Medea with penetrance 1. So, a value of 0.2 means that the resident has a penetrance 1/5 that 
#' of the invader. For the monoecious model, only the ratio of penetrances matters, not their absolute values. Defaults to seq(0.05, 0.95, 0.1).
#' @param lineweight How heavy should each line be? This passes a number to `lwd` for plotting. Defaults to 2.
#'	
#' @returns 
#' The function returns a matrix of the p_hat values: one row for each k2, one column for each value of S from 0 to 0.99
#' It also draws a plot. The lines are colored black to red as the penetrance ratio increases.
#' @examples TheoryPhat(residents = c(0.2,0.8))
#' @export
#' 

TheoryPhat <- function(residents = seq(0.05,0.95,0.1) , lineweight = 2){
	if(min(residents) < 0 | max(residents) > 1) stop("resident penetrances should be between 0 and 1.")
	colfunc <- colorRampPalette(c("red", "black"))
	rs <- 1/residents

	S <- seq(0,0.99,0.01)
	
	r <- rs[1]
	par(mar = c(4,4,1,1))
	plot((S*(r-3)+2-2*r + sqrt((1+r^2)*((S-2)^2) +2*r*(S^2+4*S -4))) / (4*(r-1)*(S-1)) ~ S, col = colfunc(1001)[1+ residents[1]*1000], ty = "l", ylim = c(0,.5), lwd = lineweight, ylab = "p_hat", xlab = "S")

	for(j in 2:length(rs)){
		r <- rs[j]
		points((S*(r-3)+2-2*r + sqrt((1+r^2)*((S-2)^2) +2*r*(S^2+4*S -4))) / (4*(r-1)*(S-1)) ~ S, col = colfunc(1001)[1+ residents[j]*1000], ty = "l", lwd = lineweight)		
	}
	par(mar = c(5, 4, 4, 2) + 0.1)
	thresh <- matrix(ncol= length(S), nrow = length(residents))
	for(a in 1:length(residents)){
		for(b in 1:length(S)){
	thresh[a,b] <- (S[b]*(rs[a]-3)+2-2*rs[a] + sqrt((1+rs[a]^2)*((S[b]-2)^2) +2*rs[a]*(S[b]^2+4*S[b] -4))) / (4*(rs[a]-1)*(S[b]-1))
		}
	}
	row.names(thresh) <- residents
	colnames(thresh) <- S	
	return(thresh)
}

#' NumericPhat
#'
#' @description
#' `NumericPhat` plots the numeric estimate of p_hat, the unstable internal equilibrium for antagonistic Medeas or peels,
#' as a function of selfing rate, for one or more ratios of the penetrances of the alleles. 
#'
#' @details
#' Go get a coffee -- this function is SLOW!
#' This function uses a grid-search to find the unstable equilibrium p_hat as a function of selfing rate, 
#' under any of the mating-system and allele-type models described in Rockman 2024, then it plots p_hat, as in Figure 7.
#' For monoecious models, it's much quicker to use `TheoryPhat`. 
#' Note that `NumericPhat` finds the equilibrium under the conditions that male and hermaphrodite initial allele frequencies
#' are identical, and further that the starting heterozygosity is zero. So it's really a model of a one-time invasion of a
#' population fixed for one allele by migrants from a population fixed for the other allele. 
#' 
#' 
#' @param k1 Penetrance of the invading allele. Defaults to 1.
#' @param k2 Penetrance of the resident allele. This can be a vector of values, each getting its own line in the plot. Defaults to c(0.05,0.35,0.65,0.95).
#' @param b.value The ratio of the number of hermaphrodite offspring from an outcross to the number from a self-fertilization. Defaults to 1,
#' implying that an outcross results in twice as many total offspring (half male, half hermaphrodite).
#' @param mod The name of the function that will iterate the recursion equations for evolution of the system. This is one of the 
#' following: AndroMedeaYp, AndroMedeaXYZ, AndroPeelYp, AndroPeelXYZ, MedPeelYp, MedPeelXYZ, PeelMedYp, PeelMedXYZ, MedeaYp, or 
#' MedeaXYZ. Note that the last two can be plotted much more efficiently using the theoretical results, using `TheoryPhat`. Defaults to AndroMedeaYp.
#' @param k.scale This is a number between zero and 1 that multiplies the specified k1 and k2 values to change their absolute magnitudes. Defaults to 1.
#' @param add If TRUE, the function will add lines to an existing plot. Defaults to FALSE.
#' @param ymax Specify the maximum value for the y-axis (value of p_hat). Defaults to 0.5.
#' @param weight How heavy should each line be? This passes a number to `lwd` for plotting. Defaults to 1.
#' @param linestyle Solid lines are 1, dashed and dotted lines are higher integers. This passes a value to `lty`. Defaults to 1.
#' @param addgray If TRUE, and also add is TRUE, the function will add additional lines to an existing plot, in gray rather than red-black. Defaults to F.
#' 
#' @returns The function returns a matrix of the plotted p_hat values: one row for each k2, one column for each value of S from 0 to 0.99. 
#' It also draws a plot. The lines are colored black to red as the penetrance ratio increases, plus gray if specified.
#' @examples NumericPhat(mod = AndroPeelYp, weight = 2)
#' @export
#' 

NumericPhat <- function(k1 = 1, k2 = c(0.05,0.35,0.65,0.95), b.value =1, mod = AndroMedeaYp, k.scale = 1, add = F, ymax = 0.5, linestyle = 1, weight = 1.5, addgray = F){
	
	if(k.scale < 0 | k.scale > 1) stop("k.scale should be between 0 and 1.")
	if(k1 < 0 | k1 > 1) stop("k1 should be between 0 and 1.")
	if(min(k2) < 0 | max(k2) > 1) stop("k2 should be between 0 and 1.")

	S <- seq(0.0,0.99, 0.01)
	ps <- seq(0.001,0.999,0.001)	
	k2s <- k2
	invaderk <- k1
	pthresh <- matrix(nrow = length(k2s), ncol =length(S))
	model <- get("mod")
	for(i in 1:(length(k2s))){
		residentk <- k2s[i]
		for(j in 1:length(S)){
			selfs <- S[j]
			updown <- NULL
			for(l in 1:999){
				updown[l] <- model(k1 = invaderk*k.scale, k2 = residentk*k.scale, n.gens = 101, b = b.value, selfing = selfs, start = c(ps[l],0))$p[100]-ps[l]	
			}
			if(updown[1] > 0){
				pthresh[i,j] <- 0
			}else{
			pthresh[i,j] <- ps[which(c(0,diff(sign(updown))) !=0)][1]	
			}
		}
	}
	row.names(pthresh) <- k2s
	colnames(pthresh) <- S

	# Plot the lines
	par(mar = c(4,4,1,1))
	colfunc <- colorRampPalette(c("red", "black"))
	if(add == F){
	plot(pthresh[1,] ~ S, ylim = c(0,ymax), ty = "l", ylab = "p_hat", xlab = "S", col = colfunc(length(k2s))[1], lwd = 1, xaxs = "i", yaxs = "i", lty = linestyle)
		if(!is.na(k2[2])){
		for(k in 2:length(k2s)){
		points(pthresh[k,]~S, col = colfunc(length(k2s))[k], lwd = 1, ty = "l", lty = linestyle)
		}
		}
	}
	if(add ==T){
		for(k in 1:length(k2s)){
			if(addgray == F){
		points(pthresh[k,]~S, col = colfunc(length(k2s))[k], lwd = 1, ty = "l", lty = linestyle)
			}else{
		points(pthresh[k,]~S, col = "gray", lwd = 1, ty = "l", lty = linestyle)
				}
		}	
	}
	par(mar = c(5, 4, 4, 2) + 0.1)
	
	return(pthresh)
}


#' FreqByTimePlot
#'
#' @description
#' `FreqByTimePlot` plots the allele frequency of a Medea or peel allele through time. It will plot multiple trajectories that differ 
#' only in the selfing rate, to illustrate the effect of selfing rate when all else is held constant. 
#'
#' @details
#' This function simply plots allele frequencies output by the recursion equations from Rockman 2024. 
#' 
#' 
#' @param start A vector of two values giving the initial allele frequency, p, and the initial heterozygosity, Y. Defaults to (0.04, 0).
#' @param k1 Penetrance of the invading allele. Defaults to 1.
#' @param k2 Penetrance of the resident allele. Defaults to 0. 
#' @param b The ratio of the number of hermaphrodite offspring from an outcross to the number from a self-fertilization. Defaults to 1,
#' implying that an outcross results in twice as many total offspring (half male, half hermaphrodite).
#' @param S Selfing rate. This can be a vector of multiple values. Defaults to seq(0,1,0.1).
#' @param plotgens The number of generations to plot. Defaults to 400.
#' @param mod The name of the function that will iterate the recursion equations for evolution of the system. This is one of the 
#' following: AndroMedeaYp, AndroPeelYp, MedPeelYp, PeelMedYp, or MedeaYp. Defaults to AndroMedeaYp, androdioecious Medea-Medea antagonism.
#' @param sexes Which sexes should be plotted? Under androdioecy, male and hermaphrodite frequencies can differ. Defaults to hermaphrodites only.
#' Specify herms, males, or both. Usually the male frequencies will so similar to hermaphrodite as to be invisible in the plot.
#' @param weights How heavy should each line be? This passes a number to `lwd` for plotting. Defaults to 2.
#' @param add If TRUE, the function will add lines to an existing plot. Defaults to FALSE.
#' @param ylim Specify the minimum and maximum value for the y-axis (allele frequency). Defaults to (0,1).
#' @param lty Solid lines are 1, dashed and dotted lines are higher integers. Defaults to 1.
#' 
#' @returns The function doesn't return an object-- it just draws a plot. The lines are colored orange to blue as a function of the selfing rate. If `sexes = "both"`, 
#' the male frequencies will be drawn at half the thickness of the hermaphrodite frequencies. 
#' @examples 
#' FreqByTimePlot(S = c(0,.5,.9), weights = 4, mod = MedeaYp)
#' FreqByTimePlot(S = c(0.5,.9), add = T, mod = AndroMedeaYp, lty = 2)
#' FreqByTimePlot(S = c(0.5,.9), add = T, mod = AndroPeelYp, lty = 3)
#' @export


FreqByTimePlot <- function(start = c(0.04, 0), k1 = 1, k2 = 0, b = 1, S = seq(0,1,1/10), plotgens = 400, mod = AndroMedeaYp, sexes = c("herms","males", "both"), weights = 2, add = F, ylim = c(0,1), lty = 1){
	
	if(start[2] - 2*(0.5-abs(start[1]-0.5)) > 1e-10) stop("starting heterozygosity is impossible")
	if(k1 < 0 | k1 > 1) stop("k1 should be between 0 and 1.")
	if(k2 < 0 | k2 > 1) stop("k2 should be between 0 and 1.")
	if(min(S) < 0 | max(S) > 1) stop("S should be between 0 and 1.")

	colfunc <- colorRampPalette(c("orange", "blue"))
	
	model <- get("mod")
	par(mar = c(4,4,1,1))
	if(add == F){
	plot(mod(n.gens = plotgens)$p ~ seq(1:plotgens), col = 0, ylab = "p", xlab = "generation", ylim = ylim)	
	}
	
	if(sexes[1] %in% c("herms", "both")){
		for(s in 1:length(S)){
			if("ph" %in% names(mod())){
				points(mod(n.gens = plotgens, selfing = S[s], start = start, k1 = k1, k2 = k2, b = b)$ph, col = colfunc(1001)[1+S[s]*1000], ty = "l", lwd = weights, lty = lty)		
			}else{
				points(mod(n.gens = plotgens, selfing = S[s], start = start, k1 = k1, k2 = k2)$p, col = colfunc(1001)[1+S[s]*1000], ty = "l", lwd = weights, lty = lty)		

			}
		}
	}
	if(sexes[1] %in% c("males","both")){
		for(s in 1:length(S)){
			points(mod(n.gens = plotgens, selfing = S[s], start = start, k1 = k1, k2 = k2, b = b)$pm, col = colfunc(1001)[1+S[s]*1000], ty = "l", lwd = weights/2, lty = lty)		
		}
	}
	par(mar = c(5, 4, 4, 2) + 0.1)
	
}


#' makegrid
#'
#' @description
#' `makegrid` is a helper function for the DeFinetti function. `makegrid` generates a list of starting genotype frequencies that
#' systematically cover genotype space. 
#' @details
#' This function returns a list of data frames. Each data frame is a set of x, y, and z frequencies that add to 1. 
#' 
#' 
#' @param point.density This is an integer that specifies the number of intervals from 0 to 1. For example, point.density = 4
#' will cause the function to generate a data frame of every combination of genotype values with x = c(0, .25, .5, .75, 1) and
#' the same for y and z, conditioned on them adding up to 1. Defaults to 10. 
#' @returns The function generates a list of three data frames, All, Homs, and Boundary. The All data frame includes every 
#' combination of x, y, and z that adds up to 1. Homs includes only the combinations where y = 0; the result is a list of 
#' every combination of homozygous genotypes only. Boundary gives a data frame of all combinations where one of the three
#' genotypes is at frequency 0; that is, it's the list of frequencies on the boundary of the DeFinetti plot. 

#' @examples 
#' makegrid(point.density = 10)
#' @export

makegrid <- function(point.density = 10){
nsteps <- point.density
xyz <- expand.grid(x = 0: nsteps, y= 0: nsteps, z = 0: nsteps)
xyz <- xyz/nsteps
All <- xyz[which(apply(xyz,1,sum) == 1),] # Positions evenly spaced across the whole plot
Homs <- All[which(All[,2] == 0),] # Positions evenly spaced along the Y=0 axis; i.e., homozygotes only
Boundary <- All[which(apply(All, 1, min) == 0),] # Positions evenly spaced along all three borders of the plot
return(list("All" = All,"Homs" = Homs, "Boundary" = Boundary))
}


#' steps
#'
#' @description
#' `steps` is a helper function for the DeFinetti function. `steps` iterates through a number of generations of 
#' recursion equations and plots the trajectories of genotype frequencies in the DeFinetti state space. 
#' @details
#' This adds genotype frequency trajectories to a DeFinetti plot. 
#' 
#' 
#' @param triplets a vector or data frame of x, y, z values that add up to 1. These are generated by `makegrid`. Each 
#' plotted trajectory will start from one of these triplets of genotype frequencies. No default. 
#' @param plotgens The number of generations to plot in each trajectory. No default.
#' @param kM1 The penetrance of allele 1 (x represents the homozygote frequency of this allele). Defaults to 1.
#' @param kM2 The penetrance of allele 2 (z represents the homozygote frequency of this allele). Defaults to 0.5
#' @param b.value The ratio of the number of hermaphrodite offspring from an outcross to the number from a self-fertilization. Defaults to 1, implying that an outcross results in twice as many total offspring (half male, half hermaphrodite).
#' @param weights How heavy should each line be? This passes a number to `lwd` for plotting. Defaults to 0.5
#' @param mod The name of the function that will iterate the recursion equations for evolution of the system. This is one of the 
#' following: AndroMedeaXYZ, AndroPeelXYZ, MedPeelXYZ, PeelMedXYZ, or MedeaXYZ. Defauts to AndroMedeaXYZ, which models two Medea alleles under androdioecy. 
#' @param sexes Which sexes should be plotted? Under androdioecy, male and hermaphrodite frequencies can differ. Defaults to hermaphrodites only. Specify `herms`, `males`, or `both`.
#' @returns The function does not return an object. It just adds lines and arrows to a DeFinetti plot, each showing `plotgens` generations of evolution. The color
#' of the line and arrow for hermaphrodites will correspond to the selfing rate in orange-blue space, as in `FreqByTimePlot`. 
#' If "males" or "both" are plotted, the males will be plotted in gray. 

#' @examples 
#' TernaryPlot(alab = "y", blab = "x", clab = "z")
#' steps(c(.3,.3,.4), plotgens=10, weights = 3, sexes = "both")
#' @export

steps <- function(triplets, plotgens, S = 0.5, kM1 = 1, kM2 = 0.5, bvalue = 1, weights = 0.5, mod = AndroMedeaXYZ, sexes = c("herms","males", "both")){
	colfunc <- colorRampPalette(c("orange", "blue"))
	initialfreqs <- as.numeric(triplets)
	model <- get("mod")
			if("ph" %in% names(mod(n.gens =2))){
			shifts <- model(start = initialfreqs, n.gens= plotgens, k1 = kM1, k2= kM2, b = bvalue, selfing = S)
			}else{
			shifts <- model(start = initialfreqs, n.gens= plotgens, k1 = kM1, k2= kM2, selfing = S)
			
			}
	thiscolor <- colfunc(1001)[1+round(S*1000)]

	for(i in 1:(plotgens-2)){
		if("Xh" %in% names(shifts)){
			if(sexes[1] %in% c("males", "both")){
			suppressWarnings(Ternary::TernaryArrows(shifts[i,c(5,4,6)], shifts[(i+1),c(5,4,6)], col = "gray", length = 0, lwd = weights))
				}
			}
		if(sexes[1] %in% c("herms","both")){	
		suppressWarnings(Ternary::TernaryArrows(shifts[i,c(2,1,3)], shifts[(i+1),c(2,1,3)], col = thiscolor, length = 0, lwd = weights))
			}
		}
		if("Xh" %in% names(shifts)){
			if(sexes[1] %in% c("males", "both")){
			suppressWarnings(Ternary::TernaryArrows(shifts[(plotgens-1),c(5,4,6)], shifts[plotgens,c(5,4,6)], col = "gray", length = 0.1, lwd = weights))
				}
			}
		if(sexes[1] %in% c("herms","both")){
			suppressWarnings(Ternary::TernaryArrows(shifts[(plotgens-1),c(2,1,3)], shifts[plotgens,c(2,1,3)], col = thiscolor, length = 0.1, lwd = weights))
			}
}

#' DeFinetti
#'
#' @description
#' `DeFinetti` plots the trajectories of genotype frequencies in the DeFinetti state space. 
#' @details
#' This function plots the evolution of Medea/peel alleles in genotype space, for any of the mating systems and  
#' allele types, selfing rates, penetrances, and so forth. The result is a figure like 4B from Rockman 2024.
#' 
#' @param startingpoints One of `"All"`, `"Boundary"`, or `"Homs"`. The function will then plot trajectories that start at points
#' equally spaced across the whole DeFinetti plot (All), or just the borders of the plot (Boundary), or just the bottom (Homs). Defaults to All.
#' @param pointnumber This is an integer that specifies how densely the trajectories should be plotted. It specifies the number of 
#' intervals from 0 to 1. For example, point.density = 4 will cause the function to plot trajectories from genotype frequencies 
#' spaced at 0, 0.25, 0.5, 0.75, and 1.  Defaults to 10. 
#' @param grid If a grid of lines in the background of the plot is desired, how dense should the lines be? Defaults to 0, no grid. 
#' `grid = 2` looks nice. 
#' @param HWE Plot a solid line to indicate genotype frequencies at Hardy-Weinberg equilibrium. Defaults to TRUE.
#' @param eHWE Plot a dashed line to indicate genotype frequencies at extended HWE, incorporating the selfing rate. Defaults to TRUE.
#' @param kscale This is a number between zero and 1 that multiplies the specified k1 and k2 values to change their absolute magnitudes. Defaults to 1.
#' @param k1 The penetrance of allele 1 (x represents the homozygote frequency of this allele). Defaults to 1.
#' @param k2 The penetrance of allele 2 (z represents the homozygote frequency of this allele). Defaults to 0.65.
#' @param S Selfing rate. Defaults to 0.57. 
#' @param plotgens The number of generations to plot in each trajectory. Defaults to 10.
#' @param b.value The ratio of the number of hermaphrodite offspring from an outcross to the number from a self-fertilization. Defaults to 1, implying that an outcross results in twice as many total offspring (half male, half hermaphrodite).
#' @param mod The name of the function that will iterate the recursion equations for evolution of the system. This is one of the 
#' following: AndroMedeaXYZ, AndroPeelXYZ, MedPeelXYZ, PeelMedXYZ, or MedeaXYZ. Defaults to AndroMedeaXYZ, which models two Medea alleles under androdioecy.
#' @param sexes One of `"herms"`, `"males"`, or `"both"`. Which sexes should be plotted? Under androdioecy, male and hermaphrodite frequencies can differ. 
#' Defaults to hermaphrodites only.
#' @param weights How heavy should each line be? This passes a number to `lwd` for plotting. Defaults to 1.
#' @param add If TRUE, the function will add lines to an existing plot. Defaults to FALSE.
#' @returns The function does not return an object; it just plots lines and arrows in a DeFinetti plot, each showing `plotgens` generations of evolution. The color
#' of the line and arrow for hermaphrodites will correspond to the selfing rate in orange-blue space, as in `FreqByTimePlot`. 
#' If "males" or "both" are plotted, the males will be plotted in gray. 
#' @examples DeFinetti(starting = "Boundary", grid = 2, S = .4, plotgens = 10, mod = MedPeelXYZ, sexes = "both", weights = 3)
#' @export
#' 


DeFinetti <- function(startingpoints = "All", pointnumber = 10, grid = 0, HWE = T, eHWE = T, kscale = 1, k1 = 1, k2 = 0.65, S = 0.57, plotgens = 10, b = 1, mod = AndroMedeaXYZ, sexes = "herms", weights = 1, add = F){
		
	if(k1 <0 | k1 >1 | k2 <0| k2>1 | S <0 | S >1) stop("k or S parameters are out of range")
	if(kscale < 0 | kscale > 1) stop("kscale should be between 0 and 1.")
	
	k1 <- k1*kscale
	k2 <- k2*kscale
	
	# Make the dataframe of starting genotype frequencies using the makegrid function
	startat <- makegrid(pointnumber)[[startingpoints[1]]]

	if(add == F){
	# Set up the plot
	par(mar = c(0,0,0,0))
	Ternary::TernaryPlot(alab = "y", blab = "x", clab = "z", grid.lines = grid)
	Ternary::TernaryPoints(startat[c(2,1,3)], pch = 19, cex= 0.2)
	}
	
	#Add HWE lines	
	p <- seq(0,1,1/100)
	selfrate <- S 
	f <- selfrate/(2-selfrate)
	if(eHWE == T){
	Ternary::TernaryLines(cbind( (1-f)*2*p*(1-p), ((1-f)*p^2 + p*f),  ((1-f)*(1-p)^2 + (1-p)*f)), col = 1, lty =2, lwd =.6) #Extended HWE under neutrality
	}
	if(HWE == T){
		Ternary::TernaryLines(cbind( 2*p*(1-p), (p^2),  ((1-p)^2)), col = 1, lwd = .6) # HWE
	}
	par(mar = c(5, 4, 4, 2) + 0.1)
	
	# Calculate and plot all the trajectories using the steps function
	suppressWarnings(apply(startat, 1, steps, kM1 = k1, kM2 = k2, S=S, b=b, plotgens = plotgens, mod = mod, sexes = sexes, weights = weights))
	return(invisible(NULL))
	
}


#' AddTheoryEquilibrium
#'
#' @description
#' `AddTheoryEquilibrium` adds the equilibrium values of p and Y under monoecy to a DeFinetti plot. 
#' @details
#' This function uses equations 5 and 6 from Rockman 2024 to find the equilibrium values of allele frequency (p) and heterozygosity (Y) 
#' and it adds them to an already-plotted DeFinetti plot. The theoretical values apply in the monoecious case only. 
#' 
#' @param k1 The penetrance of allele 1 (x represents the homozygote frequency of this allele). Defaults to 0.9.
#' @param k2 The penetrance of allele 2 (z represents the homozygote frequency of this allele). Defaults to 0.9.
#' @param S Selfing rate. Defaults to 0.3. 
#' @param lineweight How heavy should each line be? This passes a number to `lwd` for plotting. Defaults to 2.
#' @param pointsize How large should the dot representing Y_hat be? This passes to `cex` for plotting. Defaults to 1.
#' @returns The function adds a red line indicating p_hat and a red point indicating Y_hat. And it returns a list with three
#' elements: xyz is a dataframe of the X,Y,Z values of the line, at increments of 0.005 in heterozygote frequency; p_hat is the theoretical
#' value of p_hat; and y_hat is the theoretical value of y_hat. 
#' @examples DeFinetti(startingpoints = "Boundary", grid = 2, k1 = 1, k2 = 0.65, S = 0.57, plotgens = 10, mod = MedeaXYZ, weights = 2)
#' @examples AddTheoryEquilibrium(k1 = 1, k2 = 0.65, S = 0.57)
#' @export
#' 


AddTheoryEquilibrium <- function(k1 = .9, k2 = .9, S = 0.3, lineweight = 2, pointsize = 1){
	
	if(k1 <0 | k1 >1 | k2 <0| k2>1 | S <0 | S >1) stop("k or S parameters are out of range")

	r <- k1/k2
	if(r ==1){
		phat <- 0.5
		}else{
		phat <- (S*(r-3)+2-2*r + sqrt((1+r^2)*((S-2)^2) +2*r*(S^2+4*S -4))) / (4*(r-1)*(S-1))
		}
	K <- k1*(S/2 + (1-phat)-S*(1-phat)) + k2*(S/2 + phat - S*phat)
	yhat <- (1-(S/2) - sqrt(.25*S^2 + 1-S+(S-1)*(4*phat*(1-phat)*K))) / K

	yline <- seq(0,(2*phat), (2*phat)/100)
	xline <- phat - yline/2
	zline <- 1 - xline - yline
	Ternary::TernaryLines(cbind( yline, xline, zline), col = 2, lwd = lineweight) 
	Ternary::TernaryPoints(c(yhat, phat - yhat/2, (1-phat - yhat/2)),   cex = pointsize, pch = 19, col =2)
	PhatYhat <- list(data.frame(cbind(xline, yline,zline)), phat, yhat)
	names(PhatYhat) <- c("xyz", "p_hat", "Y_hat")
	return(PhatYhat)
}


#' AddNumericEquilibrium
#'
#' @description
#' `AddNumericEquilibrium ` adds the numeric estimates of equilibrium values of p and Y to a DeFinetti plot. 
#' @details
#' This function uses a grid search to find the equilibrium values of allele frequency (p) and heterozygosity (Y) 
#' and it adds them to an already-plotted DeFinetti plot. 
#' This function works for any of the mating-system and allele-type models described in Rockman 2024, but 
#' for monoecious models, it's much quicker to use `AddTheoryEquilibrium `. 
#' Note that `AddNumericEquilibrium` finds the equilibrium under the conditions that male and hermaphrodite initial allele frequencies are identical.
#' @param mod The name of the function that will iterate the recursion equations for evolution of the system. This is one of the 
#' following: AndroMedeaXYZ, AndroPeelXYZ, MedPeelXYZ, PeelMedXYZ, or MedeaXYZ. Defaults to AndroMedeaXYZ, which models two Medea alleles under androdioecy.
#' @param b The ratio of the number of hermaphrodite offspring from an outcross to the number from a self-fertilization. Defaults to 1, implying that an outcross results in twice as many total offspring (half male, half hermaphrodite).
#' @param S Selfing rate. Defaults to 0.57. 
#' @param kscale This is a number between zero and 1 that multiplies the specified k1 and k2 values to change their absolute magnitudes. Defaults to 1.
#' @param k1 The penetrance of allele 1 (x represents the homozygote frequency of this allele). Defaults to 1.
#' @param k2 The penetrance of allele 2 (z represents the homozygote frequency of this allele). Defaults to 0.65.
#' @param lineweight How heavy should each line be? This passes a number to `lwd` for plotting. Defaults to 2.
#' @returns The function adds a red line indicating p_hat, Y_hat combinations that form the unstable equilibrium. And it returns a data frame
#' of the X, Y, Z values of the line, at increments of 0.005 in heterozygote frequency. 
#' @examples DeFinetti(startingpoints = "Boundary", grid = 2, k1 = 1, k2 = 0.65, S = 0.57, plotgens = 10, mod = AndroMedeaXYZ, weights = 2)
#' @examples AddNumericEquilibrium(mod = AndroMedeaXYZ, k1 = 1, k2 = 0.65, S = 0.57)
#' @export
#' 


AddNumericEquilibrium <- function(mod = AndroMedeaXYZ, b = 1, S = 0.57, kscale =1, k1 =1, k2= 0.65, lineweight = 2){
	if(kscale < 0 | kscale > 1) stop("k.scale should be between 0 and 1.")
	k1 <- k1*kscale
	k2 <- k2*kscale
	ys <- seq(0,1,0.005)
	ps <- seq(0.001,0.999,0.001)
	model <- get("mod")
	if(k1 <0 | k1 >1 | k2 <0| k2>1 | S <0 | S >1) stop("k or S parameters are out of range")
	
	UpOrDown <- matrix(nrow = length(ys), ncol= length(ps))
	for(i in 1:length(ys)){ # for every value of y
		minp <- (ys[i]/2)
		maxp <- 1-minp	
		for(j in 1:length(ps)) { # for a bunch of values of Y
			if(ps[j] > minp && ps[j] <= maxp){
				if("ph" %in% names(mod(n.gens =2))){ # 
					if("Zh" %in% names(mod(n.gens =2))){
							startstring <- c(ps[j]-ys[i]/2, ys[i], 1-ps[j]-ys[i]/2)					
							UpOrDown[i,j] <- model(k1 = k1, k2 = k2, n.gens = 101, b = b, selfing = S, start = startstring)$p[100]-ps[j]		
						}else{
							UpOrDown[i,j] <- model(k1 = k1, k2 = k2, n.gens = 101, b = b, selfing = S, start = c(ps[j],ys[i]))$p[100]-ps[j]		
						}
					}else if("Z" %in% names(mod(n.gens=2))){
					startstring <- c(ps[j]-ys[i]/2, ys[i], 1-ps[j]-ys[i]/2)	
					UpOrDown[i,j] <- model(k1 = k1, k2 = k2, n.gens = 101, selfing = S, start = startstring)$p[100]-ps[j]			
					}else{
					UpOrDown[i,j] <- model(k1 = k1, k2 = k2, n.gens = 101, selfing = S, start = c(ps[j],ys[i]))$p[100]-ps[j]						
					}
			}
		}
	}
	Equi <- data.frame(nrow = length(ys), ncol =2)
	for(i in 1:length(ys)){
		Equi[i,1] <- ps[which(c(0,diff(sign(UpOrDown[i,]))) !=0)][1]	
		Equi[i,2] <- ys[i]
		}	

	Equilibrium <- Equi[which(!is.na(Equi[,1])),]
	names(Equilibrium) <- c("p", "Y")
	EquiXYZ <- data.frame(Equilibrium$Y, Equilibrium$p - Equilibrium$Y/2, 1-(Equilibrium$p +Equilibrium$Y/2))
	names(EquiXYZ) <- c("Y", "X","Z")
	date()	
	Ternary::TernaryLines(EquiXYZ, col = 2, lwd = lineweight) 
	return(data.frame(EquiXYZ)[c("X","Y","Z")])
	
}


#' MedeaYp
#'
#' @description
#' `MedeaYp` iterates recursion equations and outputs allele frequency, heterozygosity, and mean fitness for each generation. 
#' @details
#' This function iterates equation 2 from Rockman 2024 to calculate allele frequencies, heterozygosities, and mean fitnesses over time,
#' for any initial frequencies, selfing rates, and Medea penetrances. 
#' These equations apply to monoecious mating systems and to Medeas and peels identically. With S = 0 they also apply to dioecious obligate outcrossing.
#' @param start A vector of two numbers representing the initial allele frequency p and heterozygosity Y. Defaults to c(0.05, 0).
#' @param k1 The penetrance of allele 1 (p represents the frequency of this allele). Defaults to 0.
#' @param k2 The penetrance of allele 2. Defaults to 0.
#' @param selfing Selfing rate. Defaults to 0. 
#' @param n.gens Number of generations to iterate. Defaults to 200.
#' @returns Returns a data frame with one row for each generation, with columns p, Y (heterozygosity), and W (population mean fitness). 
#' @examples MedeaYp(start = c(0.4,0.48), selfing = 0.3, k1 = 0.4, k2 = 0.2, n.gens = 10)
#' @export
#' 


MedeaYp <- function(start = c(0.05, 0), selfing = 0, k1 = 0, k2 = 0,  n.gens = 200){

	if(start[2] - 2*(0.5-abs(start[1]-0.5)) > 1e-10) stop("starting heterozygosity is impossible")
	if(k1 <0 | k1 >1 | k2 <0| k2>1 | selfing <0 | selfing >1) stop("k or S parameters are out of range")

	p0 <- start[1]
	Y0 <- start[2] 

	p <- c(p0,rep(0,n.gens-1))
	Y <- c(Y0,rep(0,n.gens-1))	
	W <- c(1, rep(0,n.gens))		
		
	for(t in 1:n.gens){
		
		W[t] <- 1-(Y[t]/2)*(selfing*(k1+k2)/2 + (1-selfing)*(p[t]*k2 + (1-p[t])*k1))					
		p[t+1] <- (p[t] - (Y[t]*k2/2)*(selfing/2 + (1-selfing)*p[t]))/W[t]
		Y[t+1] <- (selfing*Y[t]/2 + (1-selfing)*2*p[t]*(1-p[t]))/W[t]
						
		}
		
	return(data.frame(p, Y, W)[1:n.gens,])

}


#' AndroMedeaYp
#'
#' @description
#' `AndroMedeaYp ` iterates recursion equations and outputs sex-specific allele frequency, heterozygosity, and mean fitness for each generation. 
#' @details
#' This function iterates equations from the supplement to Rockman 2024 to calculate allele frequencies, heterozygosities, and mean fitnesses over time, for any selfing rates, Medea penetrances, and outcross/self hermaphrodite offspring ratios (b). Initial allele frequency and heterozygosity is constrained to be identical for males and hermaphrodites. 
#' These equations apply to Medea elements under a Caenorhabditis-type androdioecious mating system. With S = 0 they also apply to dioecious obligate outcrossing.
#' @param start A vector of two numbers representing the initial allele frequency p and heterozygosity Y. Defaults to c(0.05, 0).
#' @param selfing Selfing rate. Defaults to 0. 
#' @param k1 The penetrance of allele 1 (p represents the frequency of this allele). Defaults to 0.
#' @param k2 The penetrance of allele 2. Defaults to 0.
#' @param n.gens Number of generations to iterate. Defaults to 200.
#' @param b The ratio of the number of hermaphrodite offspring from an outcross to the number from a self-fertilization. Defaults to 1, implying that an outcross results in twice as many total offspring (half male, half hermaphrodite).
#' @returns Returns a data frame with one row for each generation, with columns ph, Yh, pm, Ym, p, Wh, and Wm, representing first the hermaphrodite-specific ph (allele frequency), and Yh (heterozygosity), then the male-specific pm, Ym, then the global sex-weighted allele frequency p, and finally the sex-specific mean fitnesses. 
#' @examples AndroMedeaYp(start = c(0.4,0.48), selfing = 0.3, k1 = 0.4, k2 = 0.2, n.gens = 10, b = 1)
#' @export
#' 


AndroMedeaYp <- function(start = c(0.05, 0), selfing = 0, k1=0, k2=0, n.gens = 200, b = 1){

	if(start[2] - 2*(0.5-abs(start[1]-0.5)) > 1e-10) stop("starting heterozygosity is impossible")
	if(k1 <0 | k1 >1 | k2 <0| k2>1 | selfing <0 | selfing >1) stop("k or S parameters are out of range")
	
	p0 <- start[1]
	Y0 <- start[2] 

	ph <- c(p0,rep(0,n.gens-1))
	Yh <- c(Y0,rep(0,n.gens-1))	
	Wh <- c(1, rep(0,n.gens))		
	pm <- c(p0,rep(0,n.gens-1))
	Ym <- c(Y0,rep(0,n.gens-1))	
	Wm <- c(1, rep(0,n.gens))		
	p <- c(p0,rep(0,n.gens-1))

	for(t in 1:n.gens){

		Wh[t] <- selfing + b - selfing*b - (Yh[t]/2)*(k1*(selfing/2 + b*(1-pm[t]) - b*selfing*(1-pm[t])) + k2*(selfing/2 + b*pm[t] - b*selfing*pm[t]))
		ph[t+1] <- (ph[t]*(selfing+(b/2)-(selfing*b)/2) + pm[t]*(b-selfing*b)/2  - (Yh[t]*k2/2)*(selfing/2 + b*pm[t] - b*pm[t]*selfing))/Wh[t]
		Yh[t+1] <- (selfing*Yh[t]/2 + b*(1-selfing)*(ph[t]*(1-pm[t]) + (1-ph[t])*pm[t]))/Wh[t]
		
		Wm[t] <- 1-(Yh[t]/2)*(pm[t]*k2 + (1-pm[t])*k1)					
		pm[t+1] <- (1/2)*(ph[t] + pm[t] - (Yh[t]*k2*pm[t]))/Wm[t]
		Ym[t+1] <- (ph[t]*(1-pm[t]) + (1-ph[t])*pm[t])/Wm[t]		
		
		# sex-averaged allele frequency:
		p[t+1] <- (ph[t+1]*(selfing+(b/2)-(selfing*b)/2) + pm[t+1]*(b-selfing*b)/2) / (selfing + b*(1-selfing)) 
		
		}

	return(data.frame(ph, Yh, pm, Ym, p, Wh, Wm)[1:n.gens,])
	
}


#' AndroPeelYp
#'
#' @description
#' `AndroPeelYp` iterates recursion equations and outputs sex-specific allele frequency, heterozygosity, and mean fitness for each generation. 
#' @details
#' This function iterates equations from the supplement to Rockman 2024 to calculate allele frequencies, heterozygosities, and mean fitnesses over time, for any selfing rates, peel penetrances, and outcross/self hermaphrodite offspring ratios (b). Initial allele frequency and heterozygosity is constrained to be identical for males and hermaphrodites. 
#' These equations apply to peel elements under a Caenorhabditis-type androdioecious mating system. With S = 0 they also apply to dioecious obligate outcrossing.
#' @param start A vector of two numbers representing the initial allele frequency p and heterozygosity Y. Defaults to c(0.05, 0).
#' @param selfing Selfing rate. Defaults to 0. 
#' @param k1 The penetrance of allele 1 (p represents the frequency of this allele). Defaults to 0.
#' @param k2 The penetrance of allele 2. Defaults to 0.
#' @param n.gens Number of generations to iterate. Defaults to 200.
#' @param b The ratio of the number of hermaphrodite offspring from an outcross to the number from a self-fertilization. Defaults to 1, implying that an outcross results in twice as many total offspring (half male, half hermaphrodite).
#' @returns Returns a data frame with one row for each generation, with columns ph, Yh, pm, Ym, p, Wh, and Wm, representing first the hermaphrodite-specific ph (allele frequency), and Yh (heterozygosity), then the male-specific pm, Ym, then the global sex-weighted allele frequency p, and finally the sex-specific mean fitnesses. 
#' @examples AndroPeelYp(start = c(0.4,0.48), selfing = 0.3, k1 = 0.4, k2 = 0.2, n.gens = 10, b = 1)
#' @export
#' 


AndroPeelYp <- function(start = c(0.05, 0), selfing = 0, k1=0, k2=0, n.gens = 200, b=1){
	
	if(start[2] - 2*(0.5-abs(start[1]-0.5)) > 1e-10) stop("starting heterozygosity is impossible")
	if(k1 <0 | k1 >1 | k2 <0| k2>1 | selfing <0 | selfing >1) stop("k or S parameters are out of range")
	
	p0 <- start[1]
	Y0 <- start[2] 

	ph <- c(p0,rep(0,n.gens-1))
	Yh <- c(Y0,rep(0,n.gens-1))	
	Wh <- c(1, rep(0,n.gens))		
	pm <- c(p0,rep(0,n.gens-1))
	Ym <- c(Y0,rep(0,n.gens-1))	
	Wm <- c(1, rep(0,n.gens))		
	p <- c(p0,rep(0,n.gens-1))


	for(t in 1:n.gens){

		Wh[t] <- selfing + b - selfing*b - ( (k1/2)*(Yh[t]*selfing*.5 + Ym[t]*(b*(1-ph[t]) - selfing*b*(1-ph[t])))  + (k2/2)*(Yh[t]*selfing*0.5 + Ym[t]*(b*ph[t]-selfing*b*ph[t])) )
				
		ph[t+1] <- (ph[t]*(selfing+(b/2)-(selfing*b)/2) + pm[t]*(b-selfing*b)/2  - (k2/2)*(Yh[t]*selfing/2 + Ym[t]*(b*ph[t] - b*ph[t]*selfing))) /Wh[t]
		Yh[t+1] <- (selfing*Yh[t]/2 + b*(1-selfing)*(ph[t]*(1-pm[t]) + (1-ph[t])*pm[t]))/Wh[t]
		
		Wm[t] <- 1-(Ym[t]/2)*(ph[t]*k2 + (1-ph[t])*k1)					
		pm[t+1] <- (ph[t]/2 + pm[t]/2 - (Ym[t]*k2*ph[t])/2)/Wm[t]
		Ym[t+1] <- (ph[t]*(1-pm[t]) + (1-ph[t])*pm[t])/Wm[t]		

		# sex-averaged allele frequency:
		p[t+1] <- (ph[t+1]*(selfing+(b/2)-(selfing*b)/2) + pm[t+1]*(b-selfing*b)/2) / (selfing + b*(1-selfing)) 
				
		}
		
	return(data.frame(ph, Yh, pm, Ym, p, Wh, Wm)[1:n.gens,])
	
}


#' MedPeelYp
#'
#' @description
#' `MedPeelYp` iterates recursion equations and outputs sex-specific allele frequency, heterozygosity, and mean fitness for each generation. 
#' @details
#' This function iterates equations from the supplement to Rockman 2024 to calculate allele frequencies, heterozygosities, and mean fitnesses over time, for any selfing rates, peel penetrances, and outcross/self hermaphrodite offspring ratios (b). Initial allele frequency and heterozygosity is constrained to be identical for males and hermaphrodites. 
#' This function tracks the frequency of a Medea element when the alternate allele is a peel element,
#' under a Caenorhabditis-type androdioecious mating system. With S = 0 they also apply to dioecious obligate outcrossing.
#' @param start A vector of two numbers representing the initial allele frequency p and heterozygosity Y. Defaults to c(0.05, 0).
#' @param selfing Selfing rate. Defaults to 0. 
#' @param k1 The penetrance of allele 1, a Medea element (p represents the frequency of this allele). Defaults to 0.
#' @param k2 The penetrance of allele 2, a peel element. Defaults to 0.
#' @param n.gens Number of generations to iterate. Defaults to 200.
#' @param b The ratio of the number of hermaphrodite offspring from an outcross to the number from a self-fertilization. Defaults to 1, implying that an outcross results in twice as many total offspring (half male, half hermaphrodite).
#' @returns Returns a data frame with one row for each generation, with columns ph, Yh, pm, Ym, p, Wh, and Wm, representing first the hermaphrodite-specific ph (Medea allele frequency), and Yh (heterozygosity), then the male-specific pm, Ym, then the global sex-weighted Medea allele frequency p, and finally the sex-specific mean fitnesses. 
#' @examples MedPeelYp(start = c(0.4,0.48), selfing = 0.3, k1 = 0.4, k2 = 0.2, n.gens = 10, b = 1)
#' @export
#' 

MedPeelYp <- function(start = c(0.05, 0), selfing = 0, k1=0, k2=0, n.gens = 200, b =1){
	#k1 is k_Medea
	#k2 is k_Peel
	
	if(start[2] - 2*(0.5-abs(start[1]-0.5)) > 1e-10) stop("starting heterozygosity is impossible")
	if(k1 <0 | k1 >1 | k2 <0| k2>1 | selfing <0 | selfing >1) stop("k or S parameters are out of range")
	
	p0 <- start[1]
	Y0 <- start[2] 
	
	ph <- c(p0,rep(0,n.gens-1))
	Yh <- c(Y0,rep(0,n.gens-1))	
	Wh <- c(1, rep(0,n.gens))		
	pm <- c(p0,rep(0,n.gens-1))
	Ym <- c(Y0,rep(0,n.gens-1))	
	Wm <- c(1, rep(0,n.gens))		
	p <- c(p0, rep(0,n.gens-1))

	for(t in 1:n.gens){

		Wh[t] <- selfing + b -selfing*b - ((k2/2)*(Yh[t]*selfing/2 + Ym[t]*(b*ph[t] -selfing*b*ph[t])) + (k1/2)*Yh[t]*(selfing/2 + (b*(1-pm[t]) - selfing*b*(1-pm[t])))) 					
		ph[t+1] <- (ph[t]*(selfing + b/2 -selfing*b/2) + pm[t]*(b/2 - selfing*b/2) - (k2/2)*(Yh[t]*selfing/2 + Ym[t]*(b*ph[t] -selfing*b*ph[t]))) /Wh[t]
		Yh[t+1] <- (selfing*Yh[t]/2 + b*(1-selfing)*(ph[t]*(1-pm[t]) + (1-ph[t])*pm[t]))/Wh[t]
		
		Wm[t] <- 1-(Ym[t]*ph[t]*k2 + Yh[t]*(1-pm[t])*k1)/2					
		pm[t+1] <- (1/2)*( ph[t] + pm[t] - (Ym[t]*k2*ph[t]) ) /Wm[t]
		Ym[t+1] <- (ph[t]*(1-pm[t]) + (1-ph[t])*pm[t])/Wm[t]

		# sex-averaged allele frequency:
		p[t+1] <- (ph[t+1]*(selfing+(b/2)-(selfing*b)/2) + pm[t+1]*(b-selfing*b)/2) / (selfing + b*(1-selfing)) 
		
		}
		
	return(data.frame(ph, Yh, pm, Ym, p, Wh, Wm)[1:n.gens,])
	
}

#' PeelMedYp
#'
#' @description
#' `PeelMedYp` iterates recursion equations and outputs sex-specific allele frequency, heterozygosity, and mean fitness for each generation. 
#' @details
#' This function iterates equations from the supplement to Rockman 2024 to calculate allele frequencies, heterozygosities, and mean fitnesses over time, for any selfing rates, penetrances, and outcross/self hermaphrodite offspring ratios (b). Initial allele frequency and heterozygosity is constrained to be identical for males and hermaphrodites. 
#' This function tracks the frequency of a peel element when the alternate allele is a Medea element,
#' under a Caenorhabditis-type androdioecious mating system. With S = 0 they also apply to dioecious obligate outcrossing.
#' @param start A vector of two numbers representing the initial allele frequency p and heterozygosity Y. Defaults to c(0.05, 0).
#' @param selfing Selfing rate. Defaults to 0. 
#' @param k1 The penetrance of allele 1, a peel element (p represents the frequency of this allele). Defaults to 0.
#' @param k2 The penetrance of allele 2, a Medea element. Defaults to 0.
#' @param n.gens Number of generations to iterate. Defaults to 200.
#' @param b The ratio of the number of hermaphrodite offspring from an outcross to the number from a self-fertilization. Defaults to 1, implying that an outcross results in twice as many total offspring (half male, half hermaphrodite).
#' @returns Returns a data frame with one row for each generation, with columns ph, Yh, pm, Ym, p, Wh, and Wm, representing first the hermaphrodite-specific ph (peel allele frequency), and Yh (heterozygosity), then the male-specific pm, Ym, then the global sex-weighted peel allele frequency p, and finally the sex-specific mean fitnesses. 
#' @examples PeelMedYp(start = c(0.4,0.48), selfing = 0.3, k1 = 0.4, k2 = 0.2, n.gens = 10, b = 1)
#' @export

PeelMedYp <- function(start = c(0.05, 0), selfing = 0, k1=0, k2=0, n.gens = 200, b =1){
	#k1 is k_Peel
	#k2 is k_Medea
	#Here, p is the frequency of the peel allele!
	
	if(start[2] - 2*(0.5-abs(start[1]-0.5)) > 1e-10) stop("starting heterozygosity is impossible")
	if(k1 <0 | k1 >1 | k2 <0| k2>1 | selfing <0 | selfing >1) stop("k or S parameters are out of range")

	p0 <- start[1]
	Y0 <- start[2] 
	
	ph <- c(p0,rep(0,n.gens-1))
	Yh <- c(Y0,rep(0,n.gens-1))	
	Wh <- c(1, rep(0,n.gens))		
	pm <- c(p0,rep(0,n.gens-1))
	Ym <- c(Y0,rep(0,n.gens-1))	
	Wm <- c(1, rep(0,n.gens))		
	p <- c(p0, rep(0,n.gens-1))

	for(t in 1:n.gens){

		Wh[t] <- selfing*(1-Yh[t]*k2/4 - Yh[t]*k1/4) + b*(1-selfing)*(1-(Yh[t]*k2*pm[t]/2 + Ym[t]*k1*(1-ph[t])/2)) 					
		ph[t+1] <- (selfing*(ph[t]-Yh[t]*k2/4) + b*(1-selfing)*(ph[t]+pm[t]-Yh[t]*k2*pm[t])/2) /Wh[t]
		Yh[t+1] <- (selfing*Yh[t]/2 + b*(1-selfing)*(ph[t]*(1-pm[t]) + (1-ph[t])*pm[t]))/Wh[t]
		
		Wm[t] <- 1-(Yh[t]*k2*pm[t]/2 + Ym[t]*k1*(1-ph[t])/2)		
		pm[t+1] <- (ph[t]+pm[t]-Yh[t]*k2*pm[t])/2 /Wm[t]
		Ym[t+1] <- (ph[t]*(1-pm[t]) + (1-ph[t])*pm[t])/Wm[t]


		# sex-averaged allele frequency:
		p[t+1] <- (ph[t+1]*(selfing+(b/2)-(selfing*b)/2) + pm[t+1]*(b-selfing*b)/2) / (selfing + b*(1-selfing)) 
		
		}
		
	return(data.frame(ph, Yh, pm, Ym, p, Wh, Wm)[1:n.gens,])
	
}

#' MedeaXYZ
#'
#' @description
#' `MedeaXYZ` iterates recursion equations and outputs allele frequency, heterozygosity, and mean fitness for each generation. 
#' @details
#' This function iterates equation 1 from Rockman 2024 to calculate genotype frequencies and mean fitnesses over time,
#' for any initial frequencies, selfing rates, and Medea penetrances. 
#' These equations apply to monoecious mating systems and to Medeas and peels identically. With S = 0 they also apply to dioecious obligate outcrossing.
#' @param start A vector of three numbers representing the initial genotype frequency X, Y, and Z (genotypes M1M1, M1M2, and M2M2). Defaults to c(0.05, 0, 0.95).
#' @param k1 The penetrance of allele 1 (X represents the homozygote frequency of this allele). Defaults to 0.
#' @param k2 The penetrance of allele 2 (Z represents the homozygote frequency of this allele). Defaults to 0.
#' @param selfing Selfing rate. Defaults to 0. 
#' @param n.gens Number of generations to iterate. Defaults to 200.
#' @returns Returns a data frame with one row for each generation, with columns X, Y, Z, p, and W (population mean fitness). 
#' @examples MedeaXYZ(selfing = 0.5, k1 = 0.8, k2 = 0.2)
#' 
#' @export

MedeaXYZ <- function(start = c(0.05, 0, 0.95), selfing = 0, k1 = 0, k2 = 0,  n.gens = 200){

	if(abs(sum(start) -1) > 1e-10) stop("starting frequencies do not sum to 1")
	if(k1 <0 | k1 >1 | k2 <0| k2>1 | selfing <0 | selfing >1) stop("k or S parameters are out of range")

	X0 <- start[1] 
	Y0 <- start[2]
	Z0 <- start[3]
			
	X <- c(X0,rep(0,n.gens))
	Y <- c(Y0,rep(0,n.gens))
	Z <- c(Z0,rep(0,n.gens))
	W <- rep(0,(n.gens+1))
	p <- c((start[1] + start[2]/2), rep(0,n.gens))
	
	for(t in 1:n.gens){
		
		W[t] <- selfing * (X[t] + (1-k2)*Y[t]/4 + Y[t]/2 + (1-k1)*Y[t]/4 + Z[t]) + (1-selfing) * (X[t] + (1-k2/2)*Y[t]*X[t] + (1-(k1+k2)/4)*Y[t]^2 + (1-k1/2)*Y[t]*Z[t] + Z[t])

		X[t+1] <- (selfing*(X[t] + (1-k2)*Y[t]/4) + (1-selfing)*(X[t]^2 + (1-k2/2)*X[t]*Y[t] + (1-k2)*(1/4)*Y[t]^2)) / W[t]		
		Y[t+1] <- (selfing*Y[t]/2 + (1-selfing)*(2*X[t]*Z[t] +(1-Y[t]/2)*Y[t])) / W[t]
		Z[t+1] <- (selfing*(Z[t] + (1-k1)*Y[t]/4) + (1-selfing)*(Z[t]^2 + (1-k1/2)*Z[t]*Y[t] + (1-k1)*(1/4)*Y[t]^2)) / W[t]

		p[t+1] <- X[t+1] + Y[t+1]/2

		}
		
return(data.frame(X, Y, Z, p, W)[1:n.gens,])

}


#' AndroMedeaXYZ
#'
#' @description
#' `AndroMedeaXYZ` iterates recursion equations and outputs sex-specific allele frequency, heterozygosity, and mean fitness for each generation. 
#' @details
#' This function iterates equations from the supplement to Rockman 2024 to calculate genotype frequencies and mean fitnesses over time, for any selfing rates, Medea penetrances, and outcross/self hermaphrodite offspring ratios (b). Initial genotype frequencies are constrained to be identical for males and hermaphrodites. 
#' These equations apply to Medea elements under a Caenorhabditis-type androdioecious mating system. With S = 0 they also apply to dioecious obligate outcrossing.
#' @param start A vector of three numbers representing the initial genotype frequency X, Y, and Z (genotypes M1M1, M1M2, and M2M2). Defaults to c(0.05, 0, 0.95).
#' @param selfing Selfing rate. Defaults to 0. 
#' @param k1 The penetrance of allele M1 (X represents the homozygote frequency of this allele). Defaults to 0.
#' @param k2 The penetrance of allele M2 (Z represents the homozygote frequency of this allele). Defaults to 0.
#' @param n.gens Number of generations to iterate. Defaults to 200.
#' @param b The ratio of the number of hermaphrodite offspring from an outcross to the number from a self-fertilization. Defaults to 1, implying that an outcross results in twice as many total offspring (half male, half hermaphrodite).
#' @returns Returns a data frame with one row for each generation, with columns Xh, Yh, Zh, Xm, Ym, Zm, ph, pm, p, Wh, and Wm, representing first the hermaphrodite-specific genotype frequencies, then the male-specific genotype frequencies, then hermaphrodite- and male-specific allele frequencies for the M1 allele, then the global sex-weighted M1 allele frequency p, and finally the sex-specific mean fitnesses. 
#' @examples AndroMedeaXYZ(selfing = 0.5, k1 = 0.8, k2 = 0.2)
#' @export
#' 

AndroMedeaXYZ <- function(start = c(0.05, 0, 0.95), selfing = 0, k1 = 0, k2 = 0, b = 1, n.gens =200){
	
	if(abs(sum(start) -1) > 1e-10) stop("starting frequencies do not sum to 1")
	if(k1 <0 | k1 >1 | k2 <0| k2>1 | selfing <0 | selfing >1) stop("k or S parameters are out of range")
	
	Xh0 <- start[1] 
	Xm0 <- start[1] 
	Yh0 <- start[2]
	Ym0 <- start[2]
	Zh0 <- start[3] 
	Zm0 <- start[3]
	
	Xh <- c(Xh0,rep(0,n.gens))	
	Xm <- c(Xm0,rep(0,n.gens))
	
	Yh <- c(Yh0,rep(0,n.gens))
	Ym <- c(Ym0,rep(0,n.gens))
		
	Zh <- c(Zh0,rep(0,n.gens))	
	Zm <- c(Zm0,rep(0,n.gens))
	
	Wh <- rep(0,(n.gens+1))	
	Wm <- rep(0,(n.gens+1))
	
	ph <- c(start[1] + start[2]/2, rep(0,n.gens))
	pm <- c(start[1] + start[2]/2, rep(0,n.gens))
	pmean <- c(start[1] + start[2]/2, rep(0,n.gens))
	
	for(t in 1:n.gens){
		
		Wh[t] <- (selfing*(1-Yh[t]*(1/4)*(k1+k2)) + b*(1-selfing)*(1 - (Yh[t]/2)*(Xm[t]*k2 +Ym[t]*(k1+k2)/2 + Zm[t]*k1)))
		
		Xh[t+1] <- (selfing*(Xh[t] + (1-k2)*Yh[t]/4) + b*(1-selfing)*(Xh[t]*Xm[t] +Xh[t]*Ym[t]/2 + (1-k2)*Yh[t]*Xm[t]/2 + (1-k2)*Yh[t]*Ym[t]/4))/Wh[t]		
		Yh[t+1] <- (selfing*Yh[t]/2 + b*(1-selfing)*((Ym[t]+Yh[t])/2 - (Yh[t]*Ym[t])/2 +Xh[t]*Zm[t]+Zh[t]*Xm[t]))/Wh[t]
		Zh[t+1] <- 1- Xh[t+1] - Yh[t+1]	

		Wm[t] <- 1 - Yh[t]*k2*Xm[t]/2 - Yh[t]*(k1+k2)*Ym[t]/4 - Yh[t]*k1*Zm[t]/2
	
		if(Wm[t]>0){
			Xm[t+1] <- (Xh[t]*Xm[t] + Xh[t]*Ym[t]/2 + (1-k2)*Yh[t]*Xm[t]/2 + (1-k2)*Yh[t]*Ym[t]/4)/Wm[t]
			Ym[t+1] <- (Ym[t]/2 + Yh[t]*(Xm[t]+Zm[t])/2 +Xh[t]*Zm[t]+Zh[t]*Xm[t])/Wm[t]
			Zm[t+1] <- 1 - Xm[t+1] - Ym[t+1]	
		}else{
			Xm[t+1] <- 0
			Ym[t+1] <- 0 
			Zm[t+1] <- 0
		}
		
	# Allele frequencies
		ph[t+1] <- Xh[t+1] + Yh[t+1]/2
		pm[t+1] <- Xm[t+1] + Ym[t+1]/2
		pmean[t+1] <- (ph[t+1]*(selfing+(b/2)-(selfing*b)/2) + pm[t+1]*(b-selfing*b)/2) / (selfing + b*(1-selfing)) 		
		}
		p <- pmean
	return(data.frame(Xh, Yh, Zh, Xm, Ym, Zm, ph, pm, p, Wh, Wm)[1:n.gens,])

}

#' AndroPeelXYZ
#'
#' @description
#' `AndroPeelXYZ` iterates recursion equations and outputs sex-specific allele frequency, heterozygosity, and mean fitness for each generation. 
#' @details
#' This function iterates equations from the supplement to Rockman 2024 to calculate genotype frequencies and mean fitnesses over time, for any selfing rates, peel penetrances, and outcross/self hermaphrodite offspring ratios (b). Initial genotype frequencies are constrained to be identical for males and hermaphrodites. 
#' These equations apply to peel elements under a Caenorhabditis-type androdioecious mating system. With S = 0 they also apply to dioecious obligate outcrossing.
#' @param start A vector of three numbers representing the initial genotype frequency X, Y, and Z (genotypes P1P1, P1P2, and P2P2). Defaults to c(0.05, 0, 0.95).
#' @param selfing Selfing rate. Defaults to 0. 
#' @param k1 The penetrance of allele P1 (X represents the homozygote frequency of this allele). Defaults to 0.
#' @param k2 The penetrance of allele P2 (Z represents the homozygote frequency of this allele). Defaults to 0.
#' @param n.gens Number of generations to iterate. Defaults to 200.
#' @param b The ratio of the number of hermaphrodite offspring from an outcross to the number from a self-fertilization. Defaults to 1, implying that an outcross results in twice as many total offspring (half male, half hermaphrodite).
#' @returns Returns a data frame with one row for each generation, with columns Xh, Yh, Zh, Xm, Ym, Zm, ph, pm, p, Wh, and Wm, representing first the hermaphrodite-specific genotype frequencies, then the male-specific genotype frequencies, then hermaphrodite- and male-specific allele frequencies for the P1 allele, then the global sex-weighted P1 allele frequency p, and finally the sex-specific mean fitnesses. 
#' @examples AndroPeelXYZ(selfing = 0.5, k1 = 0.8, k2 = 0.2)
#' @export
#' 

AndroPeelXYZ <- function(start = c(0.05, 0, 0.95), selfing = 0, k1 = 0, k2 = 0, b = 1, n.gens =200){

	if(abs(sum(start) -1) > 1e-10) stop("starting frequencies do not sum to 1")
	if(k1 <0 | k1 >1 | k2 <0| k2>1 | selfing <0 | selfing >1) stop("k or S parameters are out of range")
	
	Xh0 <- start[1] 
	Xm0 <- start[1] 
	Yh0 <- start[2]
	Ym0 <- start[2]
	Zh0 <- start[3] 
	Zm0 <- start[3]
	
	Xh <- c(Xh0,rep(0,n.gens))	
	Xm <- c(Xm0,rep(0,n.gens))
	
	Yh <- c(Yh0,rep(0,n.gens))
	Ym <- c(Ym0,rep(0,n.gens))
		
	Zh <- c(Zh0,rep(0,n.gens))	
	Zm <- c(Zm0,rep(0,n.gens))
	
	Wh <- rep(0,(n.gens+1))	
	Wm <- rep(0,(n.gens+1))
	
	ph <- c(start[1] + start[2]/2, rep(0,n.gens))
	pm <- c(start[1] + start[2]/2, rep(0,n.gens))
	pmean <- c(start[1] + start[2]/2, rep(0,n.gens))
	
	for(t in 1:n.gens){
		
		Wh[t] <- (selfing*(1-Yh[t]*(1/4)*(k1+k2)) + b*(1-selfing)*(1 - (Ym[t]/2)*(Xh[t]*k2 +Yh[t]*(k1+k2)/2 + Zh[t]*k1)))
		
		Xh[t+1] <- (selfing*(Xh[t] + (1-k2)*Yh[t]/4) + b*(1-selfing)*(Xh[t]*Xm[t] +Yh[t]*Xm[t]/2 + (1-k2)*Xh[t]*Ym[t]/2 + (1-k2)*Yh[t]*Ym[t]/4))/Wh[t]		
		Yh[t+1] <- (selfing*Yh[t]/2 + b*(1-selfing)*((Ym[t]+Yh[t])/2 - (Yh[t]*Ym[t])/2 +Xh[t]*Zm[t]+Zh[t]*Xm[t]))/Wh[t]
		Zh[t+1] <- 1 - Xh[t+1] - Yh[t+1]	

		Wm[t] <- 1 - Ym[t]*k2*Xh[t]/2 - Ym[t]*(k1+k2)*Yh[t]/4 - Ym[t]*k1*Zh[t]/2
	
		if(Wm[t]>0){
			Xm[t+1] <- (Xh[t]*Xm[t] + Yh[t]*Xm[t]/2 + (1-k2)*Xh[t]*Ym[t]/2 + (1-k2)*Yh[t]*Ym[t]/4)/Wm[t]
			Ym[t+1] <- ((Ym[t]+Yh[t])/2 - (Yh[t]*Ym[t])/2 +Xh[t]*Zm[t]+Zh[t]*Xm[t])/Wm[t]
			Zm[t+1] <- 1 - Xm[t+1] - Ym[t+1]	
		}else{
			Xm[t+1] <- 0
			Ym[t+1] <- 0 
			Zm[t+1] <- 0
		}
		
	# Allele frequencies
		ph[t+1] <- Xh[t+1] + Yh[t+1]/2
		pm[t+1] <- Xm[t+1] + Ym[t+1]/2
		pmean[t+1] <- (ph[t+1]*(selfing+(b/2)-(selfing*b)/2) + pm[t+1]*(b-selfing*b)/2) / (selfing + b*(1-selfing)) 
		p <- pmean
				
		}

	return(data.frame(Xh, Yh, Zh, Xm, Ym, Zm, ph, pm, p, Wh, Wm)[1:n.gens,])

}


#' MedPeelXYZ
#'
#' @description
#' `MedPeelXYZ` iterates recursion equations and outputs sex-specific allele frequency, heterozygosity, and mean fitness for each generation. 
#' @details
#' This function iterates equations from the supplement to Rockman 2024 to calculate genotype frequencies and mean fitnesses over time, for any selfing rates, penetrances, and outcross/self hermaphrodite offspring ratios (b). Initial genotype frequencies are constrained to be identical for males and hermaphrodites. 
#' This function tracks the frequency of a Medea element when the alternate allele is a peel element,
#' under a Caenorhabditis-type androdioecious mating system. With S = 0 it also applies to dioecious obligate outcrossing.
#' @param start A vector of three numbers representing the initial genotype frequency X, Y, and Z (genotypes MM, MP, and PP). Defaults to c(0.05, 0, 0.95).
#' @param selfing Selfing rate. Defaults to 0. 
#' @param k1 The penetrance of allele 1, a Medea element (X represents the homozygote frequency of this allele). Defaults to 0.
#' @param k2 The penetrance of allele 2, a peel element. Defaults to 0.
#' @param n.gens Number of generations to iterate. Defaults to 200.
#' @param b The ratio of the number of hermaphrodite offspring from an outcross to the number from a self-fertilization. Defaults to 1, implying that an outcross results in twice as many total offspring (half male, half hermaphrodite).
#' @returns Returns a data frame with one row for each generation, with columns Xh, Yh, Zh, Xm, Ym, Zm, ph, pm, p, Wh, and Wm, representing first the hermaphrodite-specific genotype frequencies, then the male-specific genotype frequencies, then hermaphrodite- and male-specific allele frequencies for the Medea allele, then the global sex-weighted Medea allele frequency p, and finally the sex-specific mean fitnesses. 
#' @examples MedPeelXYZ(selfing = 0.5, k1 = 0.8, k2 = 0.2)
#' 
#' @export


MedPeelXYZ <- function(start = c(0.05, 0, 0.95), selfing = 0, k1 = 0, k2 = 0, b = 1, n.gens =200){
	#k1 is k_Medea
	#k2 is k_Peel

	if(abs(sum(start) -1) > 1e-10) stop("starting frequencies do not sum to 1")
	if(k1 <0 | k1 >1 | k2 <0| k2>1 | selfing <0 | selfing >1) stop("k or S parameters are out of range")

	Xh0 <- start[1] 
	Xm0 <- start[1] 
	Yh0 <- start[2]
	Ym0 <- start[2]
	Zh0 <- start[3] 
	Zm0 <- start[3]
	
	Xh <- c(Xh0,rep(0,n.gens))	
	Xm <- c(Xm0,rep(0,n.gens))
	
	Yh <- c(Yh0,rep(0,n.gens))
	Ym <- c(Ym0,rep(0,n.gens))
		
	Zh <- c(Zh0,rep(0,n.gens))	
	Zm <- c(Zm0,rep(0,n.gens))
	
	Wh <- rep(0,(n.gens+1))	
	Wm <- rep(0,(n.gens+1))
	
	ph <- c(start[1] + start[2]/2, rep(0,n.gens))
	pm <- c(start[1] + start[2]/2, rep(0,n.gens))
	pmean <- c(start[1] + start[2]/2, rep(0,n.gens))
	
	for(t in 1:n.gens){
		
		Wh[t] <- (selfing*(1-Yh[t]*(1/4)*(k1+k2)) + b*(1-selfing)*(1 - (Ym[t]*k2/2)*(Xh[t] +Yh[t]/2) - (Yh[t]*k1/2)*(Zm[t]+Ym[t]/2)))
		
		Xh[t+1] <- (selfing*(Xh[t] + (1-k2)*Yh[t]/4) + b*(1-selfing)*(Xh[t]*Xm[t] +Yh[t]*Xm[t]/2 + (1-k2)*Xh[t]*Ym[t]/2 + (1-k2)*Yh[t]*Ym[t]/4))/Wh[t]		
		Yh[t+1] <- (selfing*Yh[t]/2 + b*(1-selfing)*((Ym[t]+Yh[t])/2 - (Yh[t]*Ym[t])/2 +Xh[t]*Zm[t]+Zh[t]*Xm[t]))/Wh[t]
		Zh[t+1] <- 1 - Xh[t+1] - Yh[t+1]	

		Wm[t] <- (1 - (Ym[t]*k2/2)*(Xh[t] +Yh[t]/2) - (Yh[t]*k1/2)*(Zm[t]+Ym[t]/2))
	
		if(Wm[t]>0){
			Xm[t+1] <- (Xh[t]*Xm[t] + Yh[t]*Xm[t]/2 + (1-k2)*Xh[t]*Ym[t]/2 + (1-k2)*Yh[t]*Ym[t]/4)/Wm[t]
			Ym[t+1] <- ((Ym[t]+Yh[t])/2 - (Yh[t]*Ym[t])/2 +Xh[t]*Zm[t]+Zh[t]*Xm[t])/Wm[t]
			Zm[t+1] <- 1 - Xm[t+1] - Ym[t+1]
		}else{
			Xm[t+1] <- 0
			Ym[t+1] <- 0 
			Zm[t+1] <- 0
		}
		
	# Allele frequencies
		ph[t+1] <- Xh[t+1] + Yh[t+1]/2
		pm[t+1] <- Xm[t+1] + Ym[t+1]/2
		pmean[t+1] <- (ph[t+1]*(selfing+(b/2)-(selfing*b)/2) + pm[t+1]*(b-selfing*b)/2) / (selfing + b*(1-selfing)) 				
		}
		p <- pmean

	return(data.frame(Xh, Yh, Zh, Xm, Ym, Zm, ph, pm, p, Wh, Wm)[1:n.gens,])

}


#' PeelMedXYZ
#'
#' @description
#' `PeelMedXYZ` iterates recursion equations and outputs sex-specific allele frequency, heterozygosity, and mean fitness for each generation. 
#' @details
#' This function iterates equations from the supplement to Rockman 2024 to calculate genotype frequencies and mean fitnesses over time, for any selfing rates, penetrances, and outcross/self hermaphrodite offspring ratios (b). Initial genotype frequencies are constrained to be identical for males and hermaphrodites. 
#' This function tracks the frequency of a peel element when the alternate allele is a Medea element,
#' under a Caenorhabditis-type androdioecious mating system. With S = 0 it also applies to dioecious obligate outcrossing.
#' @param start A vector of three numbers representing the initial genotype frequency X, Y, and Z (genotypes PP, PM, and MM). Defaults to c(0.05, 0, 0.95).
#' @param selfing Selfing rate. Defaults to 0. 
#' @param k1 The penetrance of allele 1, a peel element (X represents the homozygote frequency of this allele). Defaults to 0.
#' @param k2 The penetrance of allele 2, a Medea element. Defaults to 0.
#' @param n.gens Number of generations to iterate. Defaults to 200.
#' @param b The ratio of the number of hermaphrodite offspring from an outcross to the number from a self-fertilization. Defaults to 1, implying that an outcross results in twice as many total offspring (half male, half hermaphrodite).
#' @returns Returns a data frame with one row for each generation, with columns Xh, Yh, Zh, Xm, Ym, Zm, ph, pm, p, Wh, and Wm, representing first the hermaphrodite-specific genotype frequencies, then the male-specific genotype frequencies, then hermaphrodite- and male-specific allele frequencies for the peel allele, then the global sex-weighted peel allele frequency p, and finally the sex-specific mean fitnesses. 
#' @examples PeelMedXYZ(selfing = 0.5, k1 = 0.8, k2 = 0.2)
#' @export


PeelMedXYZ <- function(start = c(0.05, 0, 0.95), selfing = 0, k1 = 0, k2 = 0, b = 1, n.gens =200){
	#k1 is k_peel
	#k2 is k_Medea

	if(abs(sum(start) -1) > 1e-10) stop("starting frequencies do not sum to 1")
	if(k1 <0 | k1 >1 | k2 <0| k2>1 | selfing <0 | selfing >1) stop("k or S parameters are out of range")

	Xh0 <- start[1] 
	Xm0 <- start[1] 
	Yh0 <- start[2]
	Ym0 <- start[2]
	Zh0 <- start[3] 
	Zm0 <- start[3]
	
	Xh <- c(Xh0,rep(0,n.gens))	
	Xm <- c(Xm0,rep(0,n.gens))
	
	Yh <- c(Yh0,rep(0,n.gens))
	Ym <- c(Ym0,rep(0,n.gens))
		
	Zh <- c(Zh0,rep(0,n.gens))	
	Zm <- c(Zm0,rep(0,n.gens))
	
	Wh <- rep(0,(n.gens+1))	
	Wm <- rep(0,(n.gens+1))
	
	ph <- c(start[1] + start[2]/2, rep(0,n.gens))
	pm <- c(start[1] + start[2]/2, rep(0,n.gens))
	pmean <- c(start[1] + start[2]/2, rep(0,n.gens))
	
	for(t in 1:n.gens){
		
		Wh[t] <- (selfing*(1-Yh[t]*(1/4)*(k1+k2)) + b*(1-selfing)*(1 - (Yh[t]*k2/2)*(Xm[t] +Ym[t]/2) - (Ym[t]*k1/2)*(Zh[t]+Yh[t]/2)))
		
		Xh[t+1] <- (selfing*(Xh[t] + (1-k2)*Yh[t]/4) + b*(1-selfing)*(Xh[t]*Xm[t] +Xh[t]*Ym[t]/2 + (1-k2)*Yh[t]*Xm[t]/2 + (1-k2)*Yh[t]*Ym[t]/4))/Wh[t]		
		Yh[t+1] <- (selfing*Yh[t]/2 + b*(1-selfing)*((Ym[t]+Yh[t])/2 - (Yh[t]*Ym[t])/2 +Xh[t]*Zm[t]+Zh[t]*Xm[t]))/Wh[t]
		Zh[t+1] <- 1 - Xh[t+1] - Yh[t+1]	

		Wm[t] <- (1 - (Yh[t]*k2/2)*(Xm[t] +Ym[t]/2) - (Ym[t]*k1/2)*(Zh[t]+Yh[t]/2))
	
		if(Wm[t]>0){
			Xm[t+1] <- (Xh[t]*Xm[t] +Xh[t]*Ym[t]/2 + (1-k2)*Yh[t]*Xm[t]/2 + (1-k2)*Yh[t]*Ym[t]/4)/Wm[t]
			Ym[t+1] <- ((Ym[t]+Yh[t])/2 - (Yh[t]*Ym[t])/2 +Xh[t]*Zm[t]+Zh[t]*Xm[t])/Wm[t]
			Zm[t+1] <- 1 - Xm[t+1] - Ym[t+1]
		}else{
			Xm[t+1] <- 0
			Ym[t+1] <- 0 
			Zm[t+1] <- 0
		}
		
	# Allele frequencies
		ph[t+1] <- Xh[t+1] + Yh[t+1]/2
		pm[t+1] <- Xm[t+1] + Ym[t+1]/2
		pmean[t+1] <- (ph[t+1]*(selfing+(b/2)-(selfing*b)/2) + pm[t+1]*(b-selfing*b)/2) / (selfing + b*(1-selfing)) 				
		}
		p <- pmean

	return(data.frame(Xh, Yh, Zh, Xm, Ym, Zm, ph, pm, p, Wh, Wm)[1:n.gens,])

}
```
