## Supplementary material for "Parental-effect gene-drive elements under partial selfing, or why do *Caenorhabditis* genomes have hyperdivergent regions?": MedeaFight R package and guide: PlotManuscriptFigures.R

```
# FIGURE 2:
par(mfrow = c(1,2))
#left panel
FreqByTimePlot(mod = MedeaYp, weight = 1)
FreqByTimePlot(mod = MedeaYp, weight = 1, start = c(0.9,0), ylim = c(0.9,1), plotgens = 30)

# FIGURE 3:
par(mfrow = c(1,2))
FreqByTimePlot(mod = MedeaYp, k1 = 0.9, k2 = 0.9, start = c(0.45,0), weights=1, plotgens = 100, ylim=c(0,0.5))
DeFinetti(start = "All", mod = MedeaXYZ, kscale = 1, k1 = 0.9, k2 = 0.9, S = 0.3, plotgens = 6, grid =4, pointnumber = 20)
AddTheoryEquilibrium(k1 = 0.9, k2 = 0.9, S = 0.3)

# FIGURE 4:
par(mfrow = c(1,2))
FreqByTimePlot(mod = MedeaYp, k1 = 0.9, k2 = 0.5, start = c(0.2,0), weights=1, plotgens = 150, ylim=c(0,1))
DeFinetti(start = "All", mod = MedeaXYZ, kscale = 1, k1 = 0.9, k2 = 0.5, S = 0.6, plotgens = 6, grid =4, pointnumber = 20)
AddTheoryEquilibrium(k1 = 0.9, k2 = 0.5, S = 0.6)

# FIGURE 5:
par(mfrow = c(1,1))
TheoryPhat()

# FIGURE 6:
par(mfrow = c(1,2))
#left panel
FreqByTimePlot(S = c(0,0.5,0.9), weights = 4, mod = MedeaYp)
FreqByTimePlot(S = c(0.5,0.9), add = T, mod = AndroMedeaYp, lty = 2)
FreqByTimePlot(S = c(0.5,0.9), add = T, mod = AndroPeelYp, lty = 3)
#right panel
FreqByTimePlot(S = c(0,0.5,0.9), weights = 4, mod = MedeaYp) # plots monoecious results
FreqByTimePlot(S = c(0.5,0.9), add = T, b = 0.5, mod = AndroMedeaYp, lty = 2) # plots same with AndroMedeaYp
FreqByTimePlot(S = c(0.5,0.9), add = T, b = 0.5, mod = AndroPeelYp, lty = 3) # plots same with AndroPeelYp

# FIGURE 7:
par(mfrow = c(2,2))
TheoryPhat(residents = seq(0.05,0.95,0.3), lineweight = 0.5)
# Medea - Medea
NumericPhat(mod = AndroMedeaYp, add = T, weight = 2)
NumericPhat(mod = AndroMedeaYp, k.scale = 0.1, add = T, weight = 1.5, linestyle= 3)
# peel - peel
NumericPhat(mod = AndroPeelYp, weight = 2)
NumericPhat(mod = AndroPeelYp, k.scale = 0.1, add = T, weight = 1.5,  linestyle= 3)
# peel - Medea
NumericPhat(mod = PeelMedYp, weight = 2)
NumericPhat(mod = PeelMedYp, weight = 2, k2 =1, add = T, addgray = T)
NumericPhat(mod = PeelMedYp, k.scale = 0.1, add = T, weight = 1.5, linestyle= 3)
NumericPhat(mod = PeelMedYp, k.scale = 0.1, k2 =1, add = T, addgray = T, weight = 1.5, linestyle= 3)
# Medea - peel
NumericPhat(mod = MedPeelYp, weight = 2, ymax= 0.8)
NumericPhat(mod = MedPeelYp, weight = 2, k2 =1, add = T, addgray = T)
NumericPhat(mod = MedPeelYp, k.scale = 0.1, add = T, weight = 1.5, linestyle= 3)
NumericPhat(mod = MedPeelYp, k.scale = 0.1, k2 =1, add = T, addgray = T, weight = 1.5, linestyle= 3)


# FIGURE S1:
par(mfrow = c(4,4))
figurek2 <- 0.65
figureS <- 0.57
# Medea-Medea, b = 1
FreqByTimePlot(mod = AndroMedeaYp, k1 = 1, k2 = figurek2, start = c(0.2,0), weights=1, plotgens = 150, ylim=c(0,1))
DeFinetti(start = "Boundary", mod = AndroMedeaXYZ, kscale = 1, k1 = 1, k2 = figurek2, S = figureS, plotgens = 15, grid =2, pointnumber = 10, sexes = "both", weights=2)
AddNumericEquilibrium(mod = AndroMedeaYp, k1 = 1, k2 = figurek2, S = figureS, lineweight = 4)
# Medea-Medea, b = 0.5
FreqByTimePlot(mod = AndroMedeaYp, k1 = 1, k2 = figurek2, start = c(0.2,0), weights=1, plotgens = 150, ylim=c(0,1), b = 0.5)
DeFinetti(start = "Boundary", mod = AndroMedeaXYZ, kscale = 1, k1 = 1, k2 = figurek2, S = figureS, plotgens = 15, grid =2, pointnumber = 10, sexes = "both", weights=2, b = 0.5)
AddNumericEquilibrium(mod = AndroMedeaYp, k1 = 1, k2 = figurek2, S = figureS, b = 0.5, lineweight = 4)
# peel-peel, b = 1
FreqByTimePlot(mod = AndroPeelYp, k1 = 1, k2 = figurek2, start = c(0.2,0), weights=1, plotgens = 150, ylim=c(0,1))
DeFinetti(start = "Boundary", mod = AndroPeelXYZ, kscale = 1, k1 = 1, k2 = figurek2, S = figureS, plotgens = 15, grid =2, pointnumber = 10, sexes = "both", weights=2)
AddNumericEquilibrium(mod = AndroPeelYp, k1 = 1, k2 = figurek2, S = figureS, lineweight = 4)
# peel-peel, b = 0.5
FreqByTimePlot(mod = AndroPeelYp, k1 = 1, k2 = figurek2, start = c(0.2,0), weights=1, plotgens = 150, ylim=c(0,1), b = 0.5)
DeFinetti(start = "Boundary", mod = AndroPeelXYZ, kscale = 1, k1 = 1, k2 = figurek2, S = figureS, plotgens = 15, grid =2, pointnumber = 10, sexes = "both", weights=2, b = 0.5)
AddNumericEquilibrium(mod = AndroPeelYp, k1 = 1, k2 = figurek2, S = figureS, b = 0.5, lineweight = 4)
# Medea-peel, b = 1
FreqByTimePlot(mod = MedPeelYp, k1 = 1, k2 = figurek2, start = c(0.2,0), weights=1, plotgens = 150, ylim=c(0,1))
DeFinetti(start = "Boundary", mod = MedPeelXYZ, kscale = 1, k1 = 1, k2 = figurek2, S = figureS, plotgens = 15, grid =2, pointnumber = 10, sexes = "both", weights=2)
AddNumericEquilibrium(mod = MedPeelYp, k1 = 1, k2 = figurek2, S = figureS, lineweight = 4)
# Medea-peel, b = 0.5
FreqByTimePlot(mod = MedPeelYp, k1 = 1, k2 = figurek2, start = c(0.2,0), weights=1, plotgens = 150, ylim=c(0,1), b = 0.5)
DeFinetti(start = "Boundary", mod = MedPeelXYZ, kscale = 1, k1 = 1, k2 = figurek2, S = figureS, plotgens = 15, grid =2, pointnumber = 10, sexes = "both", weights=2, b = 0.5)
AddNumericEquilibrium(mod = MedPeelYp, k1 = 1, k2 = figurek2, S = figureS, b = 0.5, lineweight = 4)
# peel-Medea, b = 1
FreqByTimePlot(mod = PeelMedYp, k1 = 1, k2 = figurek2, start = c(0.2,0), weights=1, plotgens = 150, ylim=c(0,1))
DeFinetti(start = "Boundary", mod = PeelMedXYZ, kscale = 1, k1 = 1, k2 = figurek2, S = figureS, plotgens = 15, grid =2, pointnumber = 10, sexes = "both", weights=2)
AddNumericEquilibrium(mod = PeelMedYp, k1 = 1, k2 = figurek2, S = figureS, lineweight = 4)
# peel-Medea, b = 0.5
FreqByTimePlot(mod = PeelMedYp, k1 = 1, k2 = figurek2, start = c(0.2,0), weights=1, plotgens = 150, ylim=c(0,1), b = 0.5)
DeFinetti(start = "Boundary", mod = PeelMedXYZ, kscale = 1, k1 = 1, k2 = figurek2, S = figureS, plotgens = 15, grid =2, pointnumber = 10, sexes = "both", weights=2, b = 0.5)
AddNumericEquilibrium(mod = PeelMedYp, k1 = 1, k2 = figurek2, S = figureS, b = 0.5, lineweight = 4)


# FIGURE S2:
par(mfrow = c(2,2))
# Medea - Medea
NumericPhat(mod = AndroMedeaYp, weight = 2, b = 0.5)
NumericPhat(mod = AndroMedeaYp, k.scale = 0.1, add = T, weight = 1.5, linestyle= 3, b = 0.5)
# peel - peel
NumericPhat(mod = AndroPeelYp, weight = 2, b = 0.5)
NumericPhat(mod = AndroPeelYp, k.scale = 0.1, add = T, weight = 1.5,  linestyle= 3, b = 0.5)
# peel - peel
NumericPhat(mod = PeelMedYp, weight = 2, b = 0.5)
NumericPhat(mod = PeelMedYp, weight = 2, k2 =1, add = T, addgray = T, b = 0.5)
NumericPhat(mod = PeelMedYp, k.scale = 0.1, add = T, weight = 1.5, linestyle= 3, b = 0.5)
NumericPhat(mod = PeelMedYp, k.scale = 0.1, k2 =1, add = T, addgray = T, weight = 1.5, linestyle= 3, b = 0.5)
# peel - peel
NumericPhat(mod = MedPeelYp, weight = 2, ymax= 1, b = 0.5)
NumericPhat(mod = MedPeelYp, weight = 2, k2 =1, add = T, addgray = T, b = 0.5)
NumericPhat(mod = MedPeelYp, k.scale = 0.1, add = T, weight = 1.5, linestyle= 3, b = 0.5)
NumericPhat(mod = MedPeelYp, k.scale = 0.1, k2 =1, add = T, addgray = T, weight = 1.5, linestyle= 3, b = 0.5)
```
